# Neurocognitive Vulnerability to Hypoxia Across Cognitive Domains: A Systematic Review and Meta-Analysis

**DOI:** 10.1101/2025.05.15.654374

**Authors:** Daniel J. McKeown, Douglas J. Angus, Ahmed A. Moustafa, Victor R. Schinazi

## Abstract

Hypoxia challenges brain function because of the high metabolic demands required to sustain cognition, yet its effects across cognitive abilities remain incompletely understood. We conducted a systematic review and meta-analysis of 46 studies comprising 152 effect sizes to quantify the impact of hypoxia on human cognitive performance and identify factors that moderate these effects. Overall, hypoxia was associated with a significant reduction in cognitive performance, Hedges’ *g* = −0.42, 95% CI [−0.53, −0.31]. Domain-specific analyses indicated the largest pooled impairments for processing speed (Gs; *g* = −0.62, 95% CI [−0.98, −0.26]), long-term storage and retrieval (Glr; *g* = −0.60, 95% CI [−1.03, −0.17]), and psychomotor speed (Gps; *g* = −0.50, 95% CI [−0.96, −0.05]), while smaller impairments for fluid reasoning (Gf; *g* = −0.36, 95% CI [−.55, −0.16]), short-term memory (Gsm; *g* = −0.36, 95% CI [−.56, −0.15]), and decision speed/reaction time (Gt; *g* = −0.33, 95% CI [−.55, −0.12]). Visual processing (Gv) was not significantly affected. Greater hypoxic severity, reflected by lower fraction of inspired oxygen (FiO□), was associated with larger cognitive impairments. In contrast, exposure duration, participant age, and hypoxic type were not consistently associated with cognitive effect size. Substantial heterogeneity remained across analyses, indicating considerable variability in cognitive responses to hypoxia. These findings demonstrate that hypoxia is associated with broad but heterogeneous reductions in cognitive performance, with processing speed, long-term storage and retrieval, and psychomotor speed showing the largest pooled effects. This synthesis provides a framework for investigating the neurophysiological mechanisms and individual factors underlying variability in cognitive responses to hypoxia.

---

One of the earliest real-world accounts of cognitive impairment in a high-altitude setting comes from meteorologist James Glaisher and co-pilot Henry Coxwell during their 1862 hot air balloon ascent to 11,000 m above sea level. As altitude increased, Coxwell recounts losing motor control of his hands, while Glaisher lost visual acuity and eventually consciousness. Interestingly, the occurrence of these symptoms of “balloon sickness” intensified progressively during ascent. While anecdotal, this account suggests that reductions in tissue oxygen availability (hypoxia) can induce cognitive impairments that may depend on the duration and severity of exposure. Indeed, hypoxia is a demanding physiological challenge that requires nervous, respiratory, renal, and cardiovascular adaptation. Despite advances in our understanding of physiological adaptations to hypoxia, the impact of hypoxia on cognition remains unclear, particularly in relation to exposure severity and duration.

The human brain is particularly vulnerable to reductions in oxygen availability due to its disproportionately high metabolic demand. Although representing only ∼2% of total body mass, the brain consumes approximately 20% of total oxygen at rest to sustain synaptic transmission, neural signalling, and large-scale network integration. Consequently, even transient reductions in oxygen availability can disrupt the neural processes underlying cognition ^1^.

Cognition is a complex process that involves integrating activity from multiple brain regions to interact with and respond to stimuli in the environment. Although valuable for understanding hypoxia’s impact on cognition, real-world assessments in high-altitude environments cannot fully control for changes in barometric pressure and temperature. Considering this, laboratory-based assessments that manipulate the oxygen concentration of inspired air (fraction of inspired oxygen, F_i_O_2_) allow for investigations of isolated hypoxic effects. In general, impairments in cognitive processes become evident at exposures of 2,500 to 3,000 m above sea level, but findings remain inconsistent, with hypoxia having a negligible-to-substantial impact on cognition ^2,3^. This inconsistency in findings may, in part, be due to the lack of standardisation of tasks to assess cognitive processes, with multiple tasks assessing the same process but yielding different performance outcomes. Variation in participant age, exposure severity, exposure type (e.g., normobaric, hypobaric, intermittent), exposure duration, and intrinsic brain activity (e.g., periodic and aperiodic components of electroencephalographic [EEG] activity) may also contribute to inconsistent findings ^4,5^.

Importantly, cognitive systems are unlikely to exhibit uniform vulnerability to hypoxia. Cognitive abilities differ substantially in their metabolic requirements, distributed neural architecture, and reliance on large-scale cortical integration ^6^. Differences in these underlying neural and physiological demands may therefore contribute to variability in cognitive responses to reductions in oxygen availability ^7^. Understanding this differential vulnerability is critical for identifying the neurophysiological mechanisms through which hypoxia disrupts cognition across environmental and operational settings.

Attentional tasks require the ability to sustain focus on a relevant stimulus while perceiving and appropriately ignoring irrelevant stimuli. Attention has been shown to be impaired during high-altitude exposure, with impairment increasing with the severity of exposure and the attentional demand. Attentional performance is maintained at moderate altitudes (2,200 – 3,800 m above sea level) but reduced at extreme altitudes (5, 800 m above sea level) ^8,9^, in some cases persisting once returning to sea level ^9^. Interestingly, attentional performance is more affected by hypobaric hypoxia than by normobaric hypoxia ^8^. As attentional impairments appear to be most evident under conditions of high task demand ^10^, task characteristics may be an important determinant of the observed effects of hypoxia on attention.

Tasks requiring retention and learning, such as those assessing memory-specific processes, have demonstrated similar results. Here, impairments in learning and retention of memory tasks can occur at high altitude and persist once returning to sea level ^11^. Studies on memory processing in pilots also demonstrate that increasing hypoxic severity and task complexity are associated with greater performance decrements ^12,13^. Alarmingly, individuals are mostly unaware of such impairments while actively engaged in the task. Memory systems may be particularly vulnerable to hypoxia due to the high metabolic requirements of hippocampal circuitry and the sensitivity of CA1 neurons to oxygen deprivation ^14–16^. Reductions in memory performance coincide with selective reductions in blood flow and delivery in the posterior cerebral artery, which is a primary supplier of oxygen to the hippocampus ^8^. These findings suggest that the vulnerability of hippocampal circuitry may contribute to hypoxia-induced impairments in memory-related processes.

Complex cognitive tasks, such as those requiring simultaneous processing of competing information, place substantial demands on executive control ^17,18^. Executive functions encompass multiple processes, including inhibition, cognitive flexibility, planning, and goal-directed behaviour, and therefore recruit distributed neural systems ^7^. Hypoxic exposure has been shown to impair performance on several executive tasks, particularly under more severe conditions ^19–22^. However, the diversity of tasks used to assess executive function illustrates a broader challenge within the hypoxia literature. That is, similar cognitive constructs may be assessed using tasks that place substantially different demands on underlying cognitive systems. ^20–22^.

Processing speed reflects the efficiency and rapidity with which relatively simple information can be perceived, discriminated, and processed, whereas psychomotor speed involves the rapid translation of perceptual information into an appropriate motor response. Reaction and decision speed additionally involves the speed with which an individual selects and initiates a response. Hypoxia has been shown to affect each of these processes, although the magnitude of impairment may depend on the sensory modality, task demands, and exposure conditions^10,24^. Hypoxia’s impact on processing speed is dependent on the type of stimulus being processed. Processing speed is reduced when presented with visual and auditory stimuli ^10,24,25^. However, visual processing may be more impaired due to visual receptors inherently having a greater metabolic demand ^26,27^ and involving a more diverse neural network ^28^. Considering this, the type of cognitive task is an important factor in assessing hypoxia’s impact on processing speed. Hypoxia also reduces an individual’s ability to execute motor commands in response to a stimulus ^29–32^. This is not surprising, as exposure to hypoxia has consistently been shown to reduce motor cortical and spinal output ^33–36^. Psychomotor performance is reduced with increased reaction time during perceptual-motor tasks ^29,32,37,38^. However, this may be primarily due to reductions in processing speed (e.g., reduced visual sensitivity; ^32,37^.

Emerging evidence further suggests that intrinsic brain dynamics may influence vulnerability to hypoxia. Alterations in oscillatory and aperiodic EEG activity during hypoxia have been associated with disrupted excitation-inhibition balance, altered neural efficiency, and increased neural noise ^4,5^. These neurophysiological changes may contribute to individual variability in cognitive tolerance to hypoxic exposure yet remain poorly integrated within current models of hypoxia-related cognitive impairment.

Despite decades of research ^2,3,19,39–43^, the neurocognitive mechanisms underlying hypoxia-related cognitive impairment remain poorly integrated. Previous reviews have typically focused on isolated cognitive domains, lacked quantitative synthesis across exposure characteristics, or examined hypoxia primarily from an environmental physiology perspective. In addition, previous systematic reviews have been limited by small study sizes ^3,40,41^, the inclusion of concurrent exercise interventions ^19^, or a lack of meta-analytical approaches ^2,3,39,40,42,43^, all of which have hindered the standardization of hypoxic effects between studies ^42^. Consequently, it remains unclear whether hypoxia produces generalised cognitive decline or disproportionately impairs specific neural systems and cognitive domains.

To address the heterogeneity of cognitive tasks used to assess the effects of hypoxia, theoretical frameworks can provide a consistent basis for classifying cognitive abilities and identifying commonalities across outcome measures and tasks. One such framework is the Cattell–Horn–Carroll (CHC) Theory of Cognitive Abilities ^44,45^. The CHC theory provides a hierarchical model of cognitive functioning, distinguishing general cognitive ability from broad and narrow cognitive abilities. The psychometric structure of the CHC framework has been supported through factor-analytic and cross-battery research, demonstrating that diverse cognitive tests can be meaningfully mapped onto broad cognitive abilities ^46–48^. This provides a theoretically and psychometrically grounded approach for organising the heterogeneous cognitive tasks used across studies of hypoxia and enables examination of whether hypoxic effects vary across distinct cognitive abilities.

The primary objective of this systematic review and meta-analysis was to quantify the effects of hypoxia on overall cognitive performance and specific broad cognitive abilities in humans. Cognitive domains were classified according to relevant broad CHC abilities, including fluid reasoning (*G*f), short-term memory (*G*sm), long-term storage and retrieval (*G*lr), visual processing (*G*v), processing-speed (*G*s), psychomotor speed (*G*ps), and reaction and decision speed (*G*t). A secondary objective was to determine whether participant characteristics (e.g., age) and exposure characteristics (e.g., severity, duration, and hypoxia type) moderated these effects Finally, we sought to interpret these findings within a neurophysiological framework to better understand potential mechanisms underlying differential cognitive vulnerability during hypoxia.

## RESULTS

### Data Extraction

Two authors, DJM and VRS, independently assessed the extracted data items and agreed on 98.95% of items (848/857; κ = .904), with disagreements identified for 1.05% of items (*n* = 9). Of the disputed items, eight were ultimately included following adjudication by DJA. In total, 46 data items were identified as suitable for inclusion, comprising information on author, year of publication, study design, intervention, sample characteristics, hypoxic exposure duration and severity, cognitive outcomes, and main findings pertaining to cognitive processes (Table 1 in the Supplementary Material 1).

**Table 1.** Number of studies and extracted effect sizes used in the meta-analyses of each CHC broad category

| CHC category | Studies | Effect sizes | Task measures |
| --- | --- | --- | --- |
| *Gf* | 14 | 30 | Reasoning; logical relations; Stroop/response inhibition; cognitive flexibility; tower task; decision-making/risk-taking |
| *Glr* | 6 | 11 | RAVLT; verbal/visual memory; long-term and associative memory; verbal/non-verbal fluency |
| *Gps* | 7 | 11 | Psychomotor speed; grooved pegboard; hand movement; motor speed |
| *Gs* | 16 | 27 | DSST/coding; Trail Making; processing-speed measures; Stroop/attention-response inhibition; proofreading; number comparison |
| *Gsm* | 22 | 44 | Digit Span; n-back; working-memory tasks; operational span; memory search; mathematical processing; VSC/accuracy and throughput |
| *Gt* | 9 | 21 | Reaction-time measures; PVT; Go/No-Go; Stroop/response inhibition; anticipation/reaction-time tasks |
| *Gv* | 4 | 8 | Map-and-compass; pattern recognition; spatial tracking; visual/spatial processing measures |

*Gf*, Fluid reasoning; *Glr*, Long-term storage and retrieval; *Gps*, Psychomotor speed; *Gs*, Processing speed; *Gsm*, Short-term memory; *Gt*, Decision speed/ reaction time; *Gv*, Visual processing; RAVLT, Rey Auditory Verbal Learning Test; DSST, Digit Symbol Substitution Test; VSC, Visual Sequence Comparison Test; PVT, Psychomotor Vigilance Test

Six included studies were judged to have moderate or serious risk of bias and were therefore excluded from the quantitative meta-analysis (Table 2 in Supplementary Material 1). This risk was predominantly attributable to Domain 1 (bias due to confounding; Table 3 in Supplementary Material 1), where studies either included a concomitant exercise protocol, lacked a control comparison condition (i.e., normoxia/sea level or low-lander control), or had large participant age variability within groups. These studies were retained in the qualitative synthesis but were not included in the quantitative meta-analyses. The remaining studies were judged to have low or moderate risk across the ROBINS-I domains. A further six eligible studies were retained in the qualitative synthesis but excluded from the quantitative meta-analysis because insufficient information was available to calculate a compatible effect size. Where an eligible study reported statistical significance or test statistics without sufficient descriptive or other information to derive a compatible effect size, it was not included in the corresponding meta-analysis ^30,49–52^. Of the 46 studies included in the systematic review, 34 contributed data to the quantitative meta-analysis, yielding 152 effect sizes. Studies were included in the quantitative meta-analysis where sufficient data were available to calculate heteroscedastic standardised mean differences (SMDH), expressed as Hedges’ *g*, including group means, standard deviations, and sample sizes, or where these statistics could be derived from the reported data. The Preferred Reporting Items for Systematic reviews and Meta-Analyses (PRISMA) flow diagram provides a summary of the inclusion process (Figure 1). The PRISMA checklist, a reference list of excluded data items, and the seed references used in the search are presented in Supplementary Material 2.

**Figure 1.**
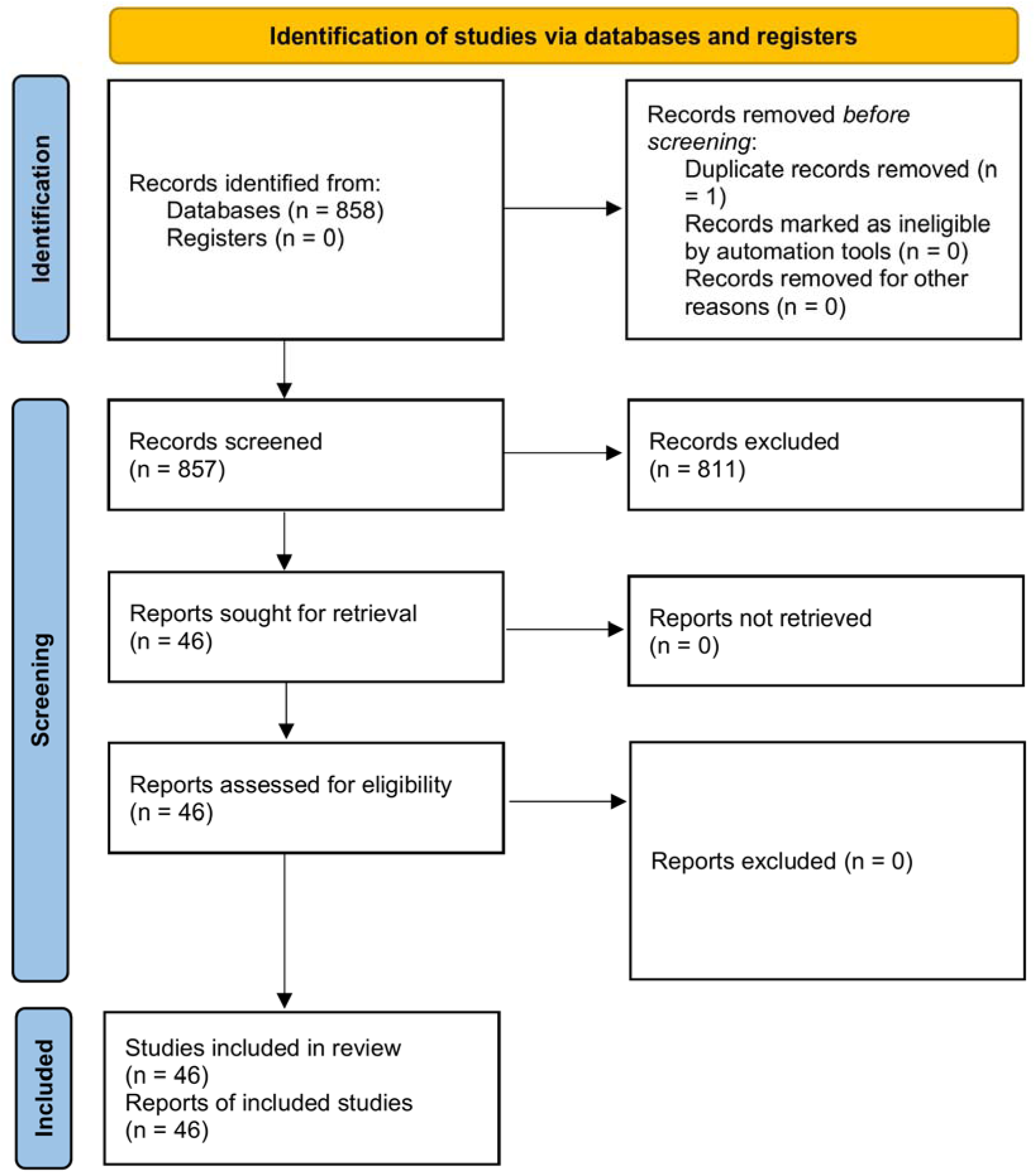
The PRISMA flow diagram summarises the identification, screening, eligibility assessment, and inclusion of studies in the systematic review. A total of 858 records were identified, with 1 duplicate removed before screening. Following title and abstract screening, 46 reports were assessed for eligibility, resulting in 46 studies included in the systematic review.

For the general cognitive ability and individual cognitive ability analyses, statistically significant effects are reported in the main text. Full results, including non-significant coefficients, are presented in the Supplementary Material 3. Funnel plots and trim-and-fill analyses were used to explore the potential influence of reporting bias. Although some funnel plots suggested possible asymmetry, pooled effect estimates were generally similar between the original random-effects models and trim-and-fill-adjusted models (Supplementary Material 3). Therefore, the primary random-effects estimates are reported in the main text. For overall cognitive ability, results from the multivariable mixed-effects moderator model are additionally reported. For individual CHC broad abilities, exploratory univariable mixed-effects moderator models are reported where sufficient data were available and including one moderator at a time (severity, duration, type, or age), whereas moderator analyses were not conducted for visual processing because of the limited number of contributing studies. In all moderator models, significant residual heterogeneity remained (see Supplementary Material 3 for model-specific heterogeneity statistics).

### General Cognitive Ability

Hypoxia was associated with a moderate negative effect on general cognitive ability, Hedge’s *g* = -.42, SE = .05, 95% CI = -.53, -.31, indicating that reductions in oxygen availability impair overall cognitive performance across domains. Effect sizes demonstrated substantial heterogeneity across studies, I^2^ = 87.12, τ² = .38, PI = -1.63, .80, suggesting that exposure- and participant-related factors contributed to variability in cognitive impairment (forest plot is presented as Figure 1 in Supplementary Material 1).

In the moderator model, hypoxia severity was significantly associated with the magnitude of the cognitive effect, with lower FiO□ (greater hypoxic severity) associated with more negative cognitive effects, β = 8.32, SE = 2.54, 95% CI = 3.34, 13.30, *p* = .001; Figure 2. Exposure duration, β = -.01, SE = .05, 95% CI = -.12, .10, *p* = .833, and participant age, β = -.02, SE = .015, 95% CI = -.05, .01, *p* = .263, were not significantly associated with cognitive effects. The coefficients for cognitive domain and hypoxia type represent differences relative to their respective reference categories and are presented in Supplementary Material 3.

**Figure 2.**
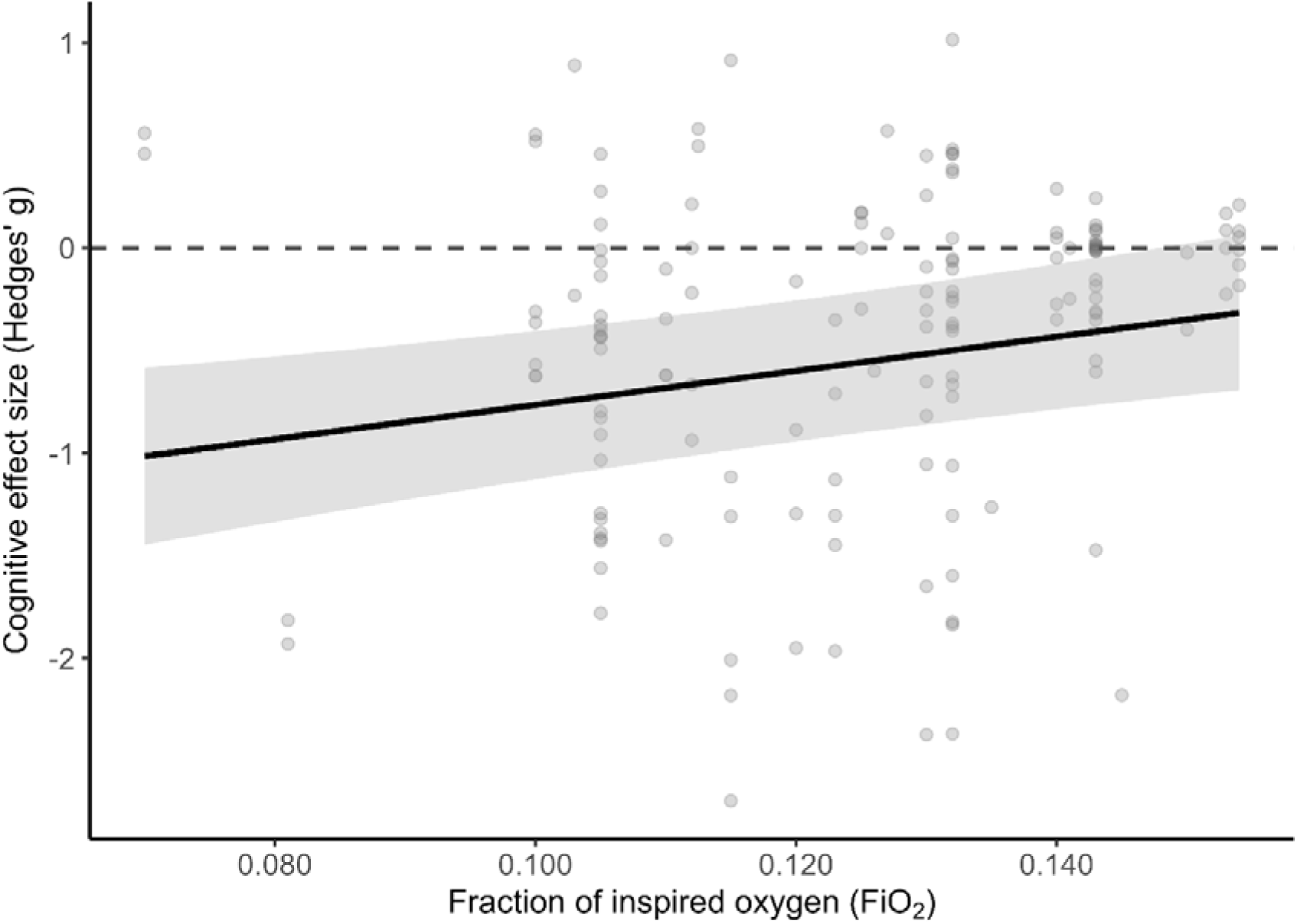
Meta-regression of the association between fraction of inspired oxygen (FiO□) and cognitive effect size. The solid line represents predicted effect sizes from the multivariable mixed-effects meta-regression, with duration and age held at their mean values and categorical moderators held at their reference levels. Shaded regions represent 95% confidence intervals. Points represent individual observed effect sizes.

### Fluid reasoning (Gf)

Hypoxia was associated with a small-to-moderate negative effect on fluid reasoning, Hedge’s *g* = -.36, SE = .1, 95% CI = -.55, -.16, *p* = .001. Effect sizes demonstrated substantial heterogeneity across studies, I^2^ = 83.09, τ² = .23, PI = -1.33, .61, suggesting that exposure-and participant-related factors contributed to variability in cognitive impairment. Funnel-plot asymmetry was explored using trim-and-fill, which imputed an additional three effect sizes.

However, the effect of hypoxia on fluid reasoning remained, Hedge’s *g* = -.27, SE = .11, 95% CI = -.48, -.06, *p* = .013. Hypoxia severity was significantly associated with the magnitude of the fluid reasoning effect, with lower FiO□ (greater hypoxic severity) associated with more negative cognitive effects, β = 10.48, SE = 4.43, 95% CI 1.79, 19.17, *p* = .018. Exposure duration, β = -.03, SE = .04, 95% CI = -.11, .05, *p* = .471, participant age, β = .01, SE = .02, 95% CI = -.03, .04, *p* = .765, and hypoxia type, β = -.01, SE = .17, 95% CI = -.33, .33, *p* = .993 were not significantly associated with fluid reasoning effects.

### Long-term storage and retrieval (Glr)

Hypoxia was associated with a moderate-to-large negative effect on long-term storage and retrieval, Hedge’s *g* = -.60, SE = .22, 95% CI = -1.03, -.17, *p* = .007. Effect sizes demonstrated substantial heterogeneity across studies, I^2^ = 83.35, τ² = .41, PI = -1.92, .72, suggesting that exposure- and participant-related factors contributed to variability in cognitive impairment. Hypoxia severity, β = 9.74, SE = 10.34, 95% CI = -10.54, 30.01, *p* = .347, duration, β = .17, SE = .16, 95% CI = -.14, .48, *p* = .279, and age, β = -.03, SE = .03, 95% CI = -.09, .03, *p* = .361, were not significantly associated with the magnitude of the long-term storage and retrieval effect.

### Psychomotor speed (Gps)

Hypoxia was associated with a moderate negative effect on psychomotor speed, Hedge’s *g* = -.50, SE = .23, 95% CI = -.96, -.05, *p* = .029. Effect sizes demonstrated substantial heterogeneity across studies, I^2^ = 86.88, τ² = .49, PI = -1.94, .94, suggesting that exposure-and participant-related factors contributed to variability in cognitive impairment. Funnel-plot asymmetry was explored using trim-and-fill, which imputed one additional effect size. However, the effect of hypoxia on psychomotor speed increased, Hedge’s *g* = -.60, SE = .23, 95% CI = -1.05, -.14, *p* = .010. Hypoxia severity, β =.91, SE = 12.79, 95% CI = -25.97, 24.15, *p* = .943, and age, β = -.02, SE = .03, 95% CI = -.08, .03, *p* = .418, were not significantly associated with the magnitude of the psychomotor speed effect.

### Processing speed (Gs)

Hypoxia was associated with a moderate-to-large negative effect on processing speed, Hedge’s *g* = -.62, SE = .18, 95% CI = -.98, -.26, *p* = .001. Effect sizes demonstrated substantial heterogeneity across studies, I^2^ = 92.07, τ² = .80, PI = -2.41, 1.18, suggesting that exposure- and participant-related factors contributed to variability in cognitive impairment. Hypoxia severity, β = 11.33, SE = 10.54, 95% CI = -9.33, 31.99, *p* = .283, duration, β = -.05, SE = .06, 95% CI = -.17, .08, *p* = .478, and age, β < .01, SE = .02, 95% CI = -.03, .03, *p* = .984, were not significantly associated with the magnitude of the processing speed effect.

### Short-term memory (Gsm)

Hypoxia was associated with a small-to-moderate negative effect on short-term memory, Hedge’s *g* = -.36, SE = .10, 95% CI = -.56, -.15, *p* = .001. Effect sizes demonstrated substantial heterogeneity across studies, I^2^ = 87.23, τ² = .38, PI = -1.59, .87, suggesting that exposure- and participant-related factors contributed to variability in cognitive impairment. Hypoxia severity, β = 4.67, SE = 4.32, 95% CI = -3.81, 13.14, *p* = .281, duration, β = -.02, SE = .03, 95% CI = -.09, .05, *p* = .551, hypoxia type, β = .15, SE = .14, 95% CI = -.13, .43, *p* = .292, and age, β = -.02, SE = .02, 95% CI = -.06, .02, *p* = .263, were not significantly associated with the magnitude of the short-term memory effect.

### Decision speed/ reaction time (Gt)

Hypoxia was associated with a small-to-moderate negative effect on decision speed/ reaction time, Hedge’s *g* = -.33, SE = .11, 95% CI = -.55, -.11, *p* = .002. Effect sizes demonstrated substantial heterogeneity across studies, I^2^ = 78.77, τ² = .19, PI = -1.21, .55, suggesting that exposure- and participant-related factors contributed to variability in cognitive impairment. Funnel-plot asymmetry was explored using trim-and-fill, which imputed an additional six effect sizes. Once imputation had occurred, the effect of hypoxia on decision speed/ reaction was no longer significant, Hedge’s *g* = -.14, SE = .11, 95% CI = -.36, .08, *p* = .211. Hypoxia severity, β = 7.76, SE = 3.97, 95% CI = -.03, 15.55, *p* = .051, duration, β < -.01, SE = .04, 95% CI = -.09, .08, *p* = .911, hypoxia type, β = -.18, SE = .15, 95% CI = - .48, .11, *p* = .215, were not significantly associated with the magnitude of the decision speed/ reaction time effect. However, age was significantly associated with the magnitude of the decision speed/ reaction time effect with older age being associated with slower decision speed/reaction time, β = .01, SE < .01, 95% CI = .00, .03, *p* = .044.

### Visual processing (Gv)

Hypoxia was not associated with visual processing, Hedge’s *g* = -.29, SE = .25, 95% CI = -.78, .21, *p* = .251. However, effect sizes demonstrated substantial heterogeneity across studies, I^2^ = 89.24, τ² = .45, PI = -1.69, 1.13. Moderator analyses were not conducted because visual processing included only 4 independent studies. Meta-regression with this number of studies would be substantially underpowered and likely to produce unstable estimates.

## DISCUSSION

In this systematic review and meta-analysis, we provide evidence that hypoxia is associated with a reduction in general cognitive ability, but that the magnitude of impairment varies across cognitive abilities. The largest pooled effects were observed for processing speed (Gs), long-term storage and retrieval (Glr), and psychomotor speed (Gps), whereas smaller effects were observed for fluid reasoning (Gf), short-term memory (Gsm), and decision speed/reaction time (Gt), and visual processing (Gv) was not significantly impaired. Greater hypoxic severity, reflected by lower FiO□, was associated with larger impairments in general cognitive ability and fluid reasoning, while exposure duration and participant age were generally not associated with the magnitude of cognitive effects. Collectively, these findings suggest that cognitive vulnerability during hypoxia varies according to the cognitive abilities engaged and the severity of oxygen reduction, while substantial residual heterogeneity indicates that additional physiological and neurobiological factors contribute to individual differences in hypoxia tolerance.

### Hypoxia differentially affects broad cognitive abilities

Hypoxia was associated with a moderate reduction in general cognitive ability, indicating that reduced oxygen availability can adversely affect cognitive performance across a range of tasks. However, the CHC-based analyses demonstrated substantial variability in the magnitude of effects across broad cognitive abilities. The largest pooled effects were observed for process speed (*G*s), long-term storage and retrieval (G*lr*), and psychomotor speed (*G*ps), whereas smaller effects were observed for fluid reasoning (G*f*), short-term memory (G*sm*), and decision speed/reaction time (*G*t). Visual processing (G*v*) was not significantly affected. This pattern suggests that cognitive abilities involving rapid information processing, memory retrieval, and sensorimotor responses may be particularly sensitive to hypoxic exposure.

Processing speed reflects the efficiency with which individuals perceive, discriminate, and response to relatively simple information, and is commonly assessed using tasks requiring rapid visual or motor responses ^23,53^. The substantial impairment observed for processing speed suggests that hypoxia may reduce the efficiency of basic information processing even when tasks place relatively limited demands on higher-order reasoning. This finding is consistent with evidence that hypoxia can alter neural processes supporting the rapid transmission and integration of information ^24,25^. However, the wide prediction interval and residual heterogeneity indicate that there is considerable variability in the magnitude and direction of effects across studies included in our analysis. Considering this, while processing speed showed the largest pooled effect, this should not be interpreted as evidence that all individuals experience substantial slowing under hypoxia, and underlying physiological factors may contribute to some individuals being more resilient to hypoxic effects ^1,5,34,36^.

Psychomotor speed was also substantially impaired, consistent with the importance of efficient sensorimotor integration for maintaining performance during hypoxia ^54^. Psychomotor speed reflects the efficiency with which perceptual information is translated into a voluntary motor response and therefore depends on coordinated activity across perceptual, cortical, spinal, and motor systems ^43^. Previous research indicates that psychomotor performance can be impaired even during relatively mild hypoxia, with greater reductions in motor speed and accuracy reported at more severe levels of exposure. At moderate-to-severe altitudes, hypoxia has been associated with impaired sensory integration ^25,55^, altered physiological tremor ^56^, reduced voluntary motor output ^35,57^, and changes in cortical and spinal motoneuron excitability ^34,36^. These physiological changes provide potential mechanisms through which hypoxia may compromise the rapid coordination of perceptual information and motor responses.

Importantly, the concurrent impairment of processing speed and psychomotor speed suggests that the effect observed for psychomotor performance cannot necessarily be attributed solely to peripheral or motor mechanisms. Although H-reflex and M-wave latencies, which provide indices of spinal and peripheral nerve conduction, typically remain relatively unchanged under hypoxic conditions ^58–60^, reductions in psychomotor performance may instead arise from disruption across multiple levels of sensorimotor processing and integration. The substantial heterogeneity in both processing and psychomotor speed further suggests that task demands and individual physiological responses may influence the extent to which these processes are affected.

Long-term storage and retrieval were similarly impaired. This finding indicates that cognitive processes involving the acquisition, storage, and retrieval of previously learned information may also be vulnerable to hypoxia. Memory processes are strongly dependent on the integrity of hippocampal formation, which is particularly susceptible to oxygen deprivation because of its high metabolic activity and dense synaptic connectivity ^61–68^. Experimental evidence further indicates that hypoxia can alter hippocampal function through changes in CA3-CA1 synaptic activity, NMDA receptor signalling, neurotransmitter systems, and the balance between excitatory and inhibitory neuronal activity ^14,69–75^. Animal studies demonstrating impaired performance on maze and passive-avoidance tasks following hypoxic exposure ^76–78^ provide additional evidence for the sensitivity of hippocampal-dependent memory processes to reduced oxygen availability. Although the present meta-analysis cannot establish a specific hippocampal mechanism, these findings provide a plausible neurobiological basis for the observed impairment in long-term storage and retrieval.

The remaining CHC abilities showed smaller pooled effects. Fluid reasoning and short-term memory were both associated with small-to-moderate impairments, while decision speed/reaction time showed a smaller negative effect. The effect for fluid reasoning is particularly relevant because hypoxic severity was also associated with the magnitude of fluid-reasoning impairment. This suggests that higher-order reasoning processes may become increasingly vulnerable as the severity of oxygen deprivation increases ^11,12^. In contrast, the absence of significant moderation by severity for short-term memory and decision speed/reaction time suggests that the factors determining vulnerability may differ across cognitive abilities. However, the decision speed/reaction time estimates should be interpreted cautiously because once accounting for interpolated effect sizes, the effect was no longer significant. This suggests that the meta-analysis was not sensitive enough, due to the limited study size or asymmetry in reporting, to detect reliable small effects of hypoxia on decision speed/reaction time.

Visual processing was the only CHC broad ability for which the pooled effect was not statistically significant. This may indicate relative preservation of some visual processing function during hypoxia, although the analysis was based on only four independent studies and therefore provides limited evidence regarding the absence of an effect. Furthermore, the prediction interval was wide and encompassed both potential determinants and improvements. Future research should aim to further elucidate the effects hypoxia has on visual processing abilities.

### Hypoxic severity and exposure characteristics

The meta-analysis included studies conducted under normobaric, hypobaric, and intermittent hypoxic conditions. Although hypoxia type was not significantly associated with the magnitude of cognitive effects in the present analyses, differences in the physiological responses elicited by these exposure conditions may contribute to the substantial heterogeneity observed across studies. For instance, hypobaric hypoxia is accompanied by reductions in atmospheric pressure and temperature, which can alter respiratory, haematological, circulatory, and metabolic responses^79^. Intermittent hypoxia may also produce distinct physiological adaptations, including hormetic responses that influence pulmonary and cardiovascular function and may affect cognitive performance, particularly in older adults ^80^. Environmental stressors accompanying hypoxic exposure may further contribute to variability in cognitive performance. For example, cold stress can impact executive functioning and memory performance, with effects depending on task complexity and attentional demand ^3^. Together these factors suggest that cognitive responses to hypoxia may be shaped not only by oxygen availability itself, but also by the broader physiological and environmental context in which hypoxia occurs.

Hypoxic severity was the clearest exposure-related moderator identified in the present meta-analysis. Lower FiO□, indicating greater hypoxic severity, was associated with more negative effects on general cognitive ability and fluid reasoning. This finding is consistent with evidence that reductions in oxygen availability and increasing hypoxic severity can exacerbate cognitive impairments, particularly when cognitive demands are high ^3^. The association between severity and fluid reasoning is particularly noteworthy, suggesting that impairments in higher-order reasoning may become more pronounced as the severity of oxygen deprivation increases. However, severity was not significantly associated with the magnitude of effects for the other CHC abilities examined, indicating that the relationship between oxygen availability and cognitive performance may vary across cognitive abilities. Thus, although greater hypoxic severity appears to increase the likelihood or magnitude of cognitive impairment, severity alone does not account for the substantial variability observed across cognitive outcomes.

In contrast, exposure duration was not significantly associated with the magnitude of cognitive effects in the present analyses. One possible explanation is that the relationship between duration and cognitive impairment may be influenced by physiological adaptation. Prolonged exposure to hypoxia can induce longer-term physiological adaptations ^79,81^, including responses mediated by hypoxia-inducible factors (HIFs) ^39,82,83^, which may mitigate some consequences of sustained reductions in oxygen availability. However, this interpretation remains speculative, as duration was not a significant moderator of general cognitive ability or the CHC-specific effects examined. The absence of a consistent duration effect may instead reflect substantial variation in exposure protocols, the timing of cognitive assessment, and the extent of physiological adaptation across studies. Collectively, these findings suggest that the severity of oxygen reduction may be more consistently related to cognitive impairment than exposure duration or hypoxia type, although considerable residual heterogeneity indicates that additional exposure, physiological, and participant-level factors contribute to individual differences in cognitive responses to hypoxia.

### Neurophysiological mechanisms of differential cognitive vulnerability during hypoxia

Differences in cognitive vulnerability may also reflect the capacity of neural systems to compensate for reduced oxygen availability. Hypoxia can alter cerebral blood flow, cortical excitability, and the balance between excitatory and inhibitory neural processes, potentially affecting the efficiency with which distributed neural networks support cognitive performance ^1,36,53^. Such compensatory mechanisms may help maintain performance under moderate hypoxic stress but may become insufficient as oxygen availability declines. This possibility is consistent with the present finding that greater hypoxic severity was associated with larger impairments in general cognitive ability and fluid reasoning. Similarly, the relatively large, pooled effect observed for long-term storage and retrieval may reflect the sensitivity of hippocampal circuitry to metabolic stress, given its high metabolic requirements and vulnerability to oxygen deprivation ^14,75^. However, the absence of significant moderation by severity for long-term storage and retrieval indicates that this proposed mechanism should not be interpreted as evidence that hippocampal-dependent processes are uniformly more sensitive to increasing hypoxic severity. These findings suggest that the magnitude of cognitive impairment during hypoxia depends not only on oxygen availability itself, but also on the extent to which a cognitive process relies on metabolically demanding neural integration. Together, these findings support a framework of differential neurocognitive vulnerability during hypoxia, whereby cognitive systems with high metabolic and integrative demands demonstrate greater susceptibility to oxygen deprivation.

The substantial unexplained heterogeneity remaining across several models further suggests that individual neurophysiological differences contribute to variability in hypoxia tolerance. Residual heterogeneity likely reflects additional physiological and neurobiological factors contributing to individual variability in hypoxia tolerance. Alterations in cortical excitability, neurovascular regulation, excitation-inhibition balance, and intrinsic brain dynamics may all influence susceptibility to cognitive impairment during hypoxia ^1,36,53^. Emerging electrophysiological evidence demonstrating altered oscillatory and aperiodic EEG activity during hypoxic exposure supports the possibility that hypoxia induces widespread changes in neural efficiency and cortical network organisation ^4,5^. These neurophysiological alterations may provide an important mechanistic framework for understanding why some individuals exhibit greater cognitive resilience during hypoxia than others.

### Electrophysiological Insights and Future Directions

Individual alpha peak frequency of the EEG reflects changes in cognitive load ^84^ and has been linked to variability in processing speed ^85^, memory ^86^, and attention ^87^. Similarly, aperiodic EEG activity, often conceptualised as reflecting neural noise and excitation-inhibition balance, has been associated with cognitive performance across younger and older populations ^53,84,88^. Individuals exhibiting steeper aperiodic exponents and greater offsets, reflective of altered excitation-inhibition dynamics and neural population activity, often demonstrate better performance across processing speed ^53^, memory ^89,90^, and executive function ^91^ tasks. Considering the substantial unexplained heterogeneity observed across several models in the present review, these intrinsic electrophysiological characteristics may contribute to variability in cognitive vulnerability during hypoxia.

Although the review focused on behavioural outcomes, interpretation within the context of neurophysiological frameworks may provide insight into the neural systems most vulnerable to hypoxia. Electrophysiological research in hypoxia has primarily focused on oscillatory activity ^5^. Hypoxic exposure has been shown to reduce peak alpha frequency, characterised by reductions in alpha power alongside increases in lower-frequency theta activity ^92^, although findings remain inconsistent ^93^. Emerging evidence further suggests that hypobaric hypoxia reduces the aperiodic exponent, indicative of increased neural noise and altered excitation-inhibition balance ^4^. Interestingly, these electrophysiological alterations resemble non-pathological age-related changes in intrinsic brain activity, potentially contributing to the greater cognitive vulnerability observed in older individuals during hypoxia.

Together, these findings suggest that alterations in intrinsic brain dynamics may contribute to domain-specific cognitive vulnerability during hypoxia. Future research should therefore integrate behavioural and electrophysiological approaches to determine how oscillatory and aperiodic EEG activity relate to individual differences in hypoxia tolerance and cognitive resilience.

### Limitations

Several aspects of the review process should be considered when interpreting these findings. The review followed PRISMA 2020 guidelines and incorporated duplicate screening, independent data extraction, and independent risk-of-bias assessment to maximise methodological rigour. However, the search was limited to English-language publications and did not include a systematic search of grey literature or unpublished studies, which may have limited the identification of all relevant evidence. Nevertheless, assessment of publication bias using funnel plots and trim-and-fill analyses suggested that the overall conclusions were robust to potential reporting bias. Although a formal certainty-of-evidence framework (e.g., GRADE) was not applied, confidence in the findings was supported through comprehensive risk-of-bias assessment, moderator analyses, and evaluation of statistical heterogeneity.

Despite accounting for cognitive ability and key study and participant characteristics, substantial unexplained heterogeneity remained across the meta-analyses. Cognitive tasks often recruit overlapping neural and cognitive processes, which may limit the extent to which broad cognitive abilities can be separated empirically. Furthermore, comparisons between paradigms assessing a common cognitive ability remain challenging because tasks can differ substantially in their cognitive demands and construct specificity^42^. The broad range of paradigms used across studies may therefore have introduced additional variability, even when tasks were classified within the same CHC broad ability. Although standardised effect sizes and multilevel meta-analytic models were used to account for differences in study-level effects and reduce the influence of methodological variability, considerable residual heterogeneity remained. This heterogeneity indicates that factors not captured by the present moderator analyses, including differences in task characteristics, participant physiology, exposure protocols, and individual responses to hypoxia—may contribute to variability in cognitive effects.

Older adults remain comparatively underrepresented within the hypoxia literature, despite age-related changes in cognitive and cerebrovascular function that may influence responses to reduced oxygen availability ^84,94^. In the present meta-analysis, however, participant age was generally not significantly associated with the magnitude of hypoxia-related cognitive effects. Age was not a significant moderator of general cognitive ability or the majority of CHC broad abilities examined, although a significant association was observed for decision speed/reaction time. Given the relatively small effect and the sensitivity of this association to potential reporting asymmetry, this finding should be interpreted cautiously. The absence of consistent age-related moderation suggests that chronological age alone may be insufficient to explain individual differences in cognitive responses to hypoxia. Instead, age-related differences in cerebrovascular function, baseline cognitive capacity, physiological responses to hypoxia, and compensatory neural mechanisms may contribute to individual variability in hypoxia tolerance ^95,96^. Together with the substantial residual heterogeneity observed across the meta-analyses, these findings highlight the importance of investigating physiological and neurobiological factors beyond chronological age when examining individual susceptibility to hypoxia-related cognitive impairment.

### Conclusion

Collectively, these findings provide evidence that hypoxia is associated with reduced cognitive performance, but that the magnitude of impairment varies across cognitive abilities. The largest pooled effects were observed for processing speed, long-term storage and retrieval, and psychomotor speed, while visual processing was not significantly impaired. Greater hypoxic severity, reflected by lower FiO□, was associated with greater impairment in general cognitive ability and fluid reasoning, whereas exposure duration, hypoxia type, and participant age were generally not associated with effect magnitude. Substantial residual heterogeneity across analyses indicates that cognitive responses to hypoxia are influenced by factors beyond oxygen availability alone, including potentially individual physiological and neurophysiological differences. Future research should therefore investigate the mechanisms underlying individual variability in hypoxia tolerance, including neurophysiological markers that may help identify individuals at greater risk of cognitive impairment in low-oxygen environments.

## METHOD

This systematic and meta-analytical review aligned with the 2020 PRISMA guidelines ^97^. The PRISMA checklist accompanying this article can be found in the Supplementary Material 2. This systematic review was not prospectively registered. No formal review protocol was prepared prior to commencement of the study.

### Eligibility Criteria

Eligibility criteria conformed to the Population, Intervention, Comparator, Outcome, Setting, Types (PICOST) framework. Data items included all relevant randomised-controlled trials, observational studies, longitudinal studies, cohort studies, case-control studies, before-after studies, and cross-sectional studies or surveys involving healthy humans exposed to either environmental (high altitude) or simulated (laboratory-based) hypoxia. Studies were included if they used normal oxygen, sea-level conditions, or lowlanders as a comparator and measured outcomes related to memory, attention, executive function, processing speed, or psychomotor speed. Data items with hypoxic exposure due to pathology (e.g., chronic obstructive pulmonary disease), with interventions that were pharmaceutical or exercise-based, which did not have suitable comparators, or were conducted in a hospital-based setting, were excluded from the review. Included studies were grouped according to cognitive outcome, with effect sizes synthesised separately for overall cognitive ability and the predefined cognitive domains of attention, memory, executive function, processing speed, and psychomotor speed.

### Data Sources & Search Strategy

The Evidence Research Accelerator ^98^ suite was used to aid in the data source selection and search strategy, and in the data screening and selection process. The TERA SearchRefiner tool was used to aid in data sourcing and to develop the search strategy. A literature search querying SCOPUS, PubMed, Medline, and Cochrane Library from inception to 6^th^ January 2025 was performed. Searches were conducted using combinations of keywords and Boolean operators relating to hypoxia and cognition. The core search strategy combined terms for hypoxia (“hypoxia” OR “altitude” OR “hypobaric”) with terms for cognition (“cognition” OR “attention” OR “memory” OR “executive function” OR “processing speed” OR “psychomotor speed”). Database-specific subject headings and syntax were used where appropriate. The complete search strategies for each database and the 13 seed PMIDs that were used as “relevant citations” to aid in refining the literature search, are provided in Supplementary Material 2. In total, 858 data items were identified in the literature search. Using the Deduplicator tool ^99^ of TERA, one data item was identified as ‘likely being a duplicate’ and was removed ^100^, bringing the total number of data items to 857.

### Selection & Data Collection Process

Authors DJM and VRS independently screened the citation search, titles/abstracts, full texts, and trial registries for inclusion of data items against the inclusion and exclusion criteria. Any disagreements regarding data item inclusion were resolved by referring to the third author, DJA. No automation/AI was used for study eligibility (other than TERA for deduplication/search refinement). Data extraction was performed by one reviewer (DJM) using a standardised data extraction form developed a priori. A second reviewer (VRS) independently verified the extracted data for accuracy and completeness. Any discrepancies were resolved through discussion with a third reviewer (DJA) until consensus was reached. Where necessary, the original publications were re-examined to clarify unclear or incomplete information. No study investigators were contacted for additional information.

### Data Items

The primary outcome was cognitive performance during hypoxic exposure compared with a normoxic or sea-level comparator. Tasks were classified according to their primary cognitive process and the nature of the performance outcome, using the Cattell–Horn–Carroll (CHC) broad-category framework ^44^. These categories consisted of fluid reasoning (*G*f), short-term memory (*G*sm), long-term storage and retrieval (*G*lr), visual processing (*G*v), processing-speed (*G*s), psychomotor speed (*G*ps), and reaction and decision speed (*G*t). Where a task involved multiple cognitive processes, it was assigned to the CHC broad ability most directly targeted by the task and its primary outcome measure. Where studies reported multiple tasks assessing the same cognitive domain, all eligible outcomes were extracted to maximise available information for the multilevel meta-analysis. When a study reported outcomes across multiple cognitive domains, effect sizes were extracted for each relevant domain. If multiple time points were reported, data corresponding to the hypoxic exposure period were extracted. Table 1 contains the number of studies and number of extracted effect sizes for each cognitive domain used in the meta-analyses.

The following study characteristics were extracted from each eligible study: first author, publication year, study design, sample size, participant age, sex (where reported), health status, hypoxia exposure type (normobaric or hypobaric), simulated altitude or inspired oxygen fraction (FiO□), exposure duration, comparator condition, cognitive task(s), cognitive domain classification, outcome statistics required for effect size calculation, and study quality information for risk-of-bias assessment.

Where information was incomplete or unclear, data were extracted from tables, figures, or supplementary materials when possible. No assumptions were made regarding missing outcome data beyond those required for effect size calculation, and studies lacking sufficient statistical information were excluded from quantitative synthesis.

### Risk of Bias

Risk of bias was assessed using the ROBINS-I (Risk Of Bias In Non-randomized Studies of Interventions) tool, which is recommended for evaluating non-randomised intervention studies. Bias was evaluated across seven domains: bias due to confounding (Domain 1), bias in the classification of interventions (Domain 2), bias in the selection of participants (Domain 3), bias due to deviations from intended interventions (Domain 4), bias due to missing data (Domain 5), bias in the measurement of outcomes (Domain 6), and bias in the selection of the reported result (Domain 7). For Domain 1, potential confounding factors included participant-, task-, environmental-, and methodology-related variables (Supplementary Table 3). Risk-of-bias assessments were performed independently by authors DJM and VRS. Any disagreements were resolved through discussion and, where consensus could not be reached, were adjudicated by the third author (DJA). No automation tools were used during the risk-of-bias assessment. Studies judged to have moderate or severe risk of bias are summarised in Table 2 of Supplementary Material 1 (n = 6).

### Certainty of Evidence

A formal certainty-of-evidence assessment (e.g., GRADE) was not performed because the review synthesised heterogeneous behavioural outcomes across multiple cognitive domains using multilevel meta-analytic models. Confidence in the findings was instead evaluated through assessment of study quality using ROBINS-I, statistical heterogeneity, moderator analyses, and reporting bias assessments.

### Meta-analysis

Meta-analyses were conducted in *R* (v. 4.3.1) using the *metafor* package (v. 4.8-0). Effect sizes were expressed as Hedges’ *g*, with positive values indicating better cognitive performance under hypoxic conditions and negative values indicating worse cognitive performance. Sampling variances (*vi*) were used to weight effect sizes, and all models were estimated using restricted maximum likelihood (REML).

Prior to analysis, effect sizes from Malle et al. (2013) were excluded because they were identified as an outlier. Hypoxic exposure duration was log-transformed using log(duration + 1) to reduce skewness and account for substantial variability in exposure duration across studies. Continuous moderators, including hypoxia severity, log-transformed duration, and participant age, were mean-centred before meta-regression analyses. Hypoxia type and Cattell–Horn–Carroll (CHC) broad cognitive ability classifications were treated as categorical variables.

A general cognitive ability analysis was conducted using all eligible cognitive effect sizes. Cognitive ability was additionally examined according to CHC broad abilities, including fluid reasoning (Gf), long-term storage and retrieval (Glr), psychomotor speed (Gps), processing speed (Gs), short-term memory (Gsm), decision speed/reaction time (Gt), and visual processing (Gv). The number of effect sizes and independent studies contributing to each cognitive ability analysis was documented.

For each outcome, an initial reduced random-effects model without moderators was estimated using the rma() function. These models provided pooled estimates of the overall effect of hypoxia on cognitive performance. Moderator analyses were subsequently conducted using mixed-effects multilevel meta-regression models implemented using the rma.mv() function, with study included as a random effect to account for the clustering of multiple effect sizes within studies.

For the overall cognitive ability analysis, a multivariable mixed-effects model simultaneously included hypoxia severity, log-transformed duration, participant age, CHC broad cognitive ability, and hypoxia type as moderators. This model was used to examine whether the overall effects of hypoxia differed according to characteristics of the hypoxic exposure, participants, and cognitive outcomes.

For individual CHC broad abilities, the number of independent studies contributing to each analysis was insufficient to support similarly complex multivariable models. Therefore, exploratory univariable mixed-effects meta-regressions were conducted separately for each moderator where there was sufficient data representation. Depending on the availability and distribution of data within each cognitive ability, these models examined the effects of hypoxia severity, log-transformed duration, participant age, and hypoxia type. For moderator analyses involving duration, effect sizes with missing duration data were excluded only from the relevant duration model and were retained in the primary meta-analysis and other moderator analyses.

Hypoxia type was examined only where meaningful comparisons between adequately represented categories were possible. For some cognitive abilities, analyses were restricted to hypobaric and normobaric hypoxia because other exposure categories were sparsely represented or absent. Hypoxia type was not examined for outcomes where the distribution of effect sizes across exposure categories was insufficient to support a stable comparison. Similarly, moderator analyses were not conducted for visual processing because only four independent studies contributed data, making meta-regression estimates highly unstable and underpowered.

Statistical heterogeneity in the reduced random-effects models was assessed using Cochran’s *Q* statistic and quantified using the *I*² statistic. Moderator models were interpreted as exploratory, particularly for individual cognitive abilities with relatively small numbers of independent studies. No formal sensitivity analyses were conducted.

Potential reporting bias was assessed visually using funnel plots and exploratorily using the trim-and-fill procedure implemented in the *metafor* package. Funnel plots were inspected for evidence of asymmetry consistent with potential small-study or reporting effects. Where applied, the trim-and-fill procedure was used to estimate the potential influence of asymmetry on the pooled effect size. Funnel plots and trim-and-fill estimates were interpreted with caution for cognitive abilities represented by a small number of independent studies, as these methods have limited reliability under such conditions. To visualise the relationship between hypoxia severity and general cognitive performance, predictions were generated from the multivariable mixed-effects meta-regression model across the observed range of hypoxia severity. Other continuous moderators were held at their mean values, while categorical moderators were held at their respective reference levels.

## AVAILABILITY OF DATA AND MATERIALS

The dataset supporting the conclusions of this article and additional files are available in the GitHub repository, https://github.com/MindSpaceLab/Hypoxia_Cognition_Review

## AUTHOR CONTRIBUTIONS

Conceptualisation: D.J.M., D.J.A, and V.R.S.; Data curation: D.J.M., D.J.A, and V.R.S.; Formal analysis: D.J.M., D.J.A, A.A.M., and V.R.S; Project administration: D.J.M.; Visualisation: D.J.M.; Writing - original draft: D.J.M., D.J.A, A.A.M., and V.R.S.; Writing - review & editing: D.J.M., D.J.A, A.A.M., and V

