## Supplementary Material 1 for "Neurocognitive Vulnerability to Hypoxia Across Cognitive Domains: A Systematic Review and Meta-Analysis"

**TABLES**

*Table 1. Characteristics of included data items (n = 46).*

| Author & Year | Design | Intervention | Sample characteristics | Duration | Severity | Outcome measures | Main cognitive findings |
|---|---|---|---|---|---|---|---|
| (Aebi et al., 2020) | Non-randomised cross-over | NN, HH, NH, HN | 16 healthy pilot trainees, 26 ± 4 yr | 30 min | NN, HN = 440m<br>HH, NH = 5500m | KLT<br>KSS | ↑ KLT error in NH & HH<br>↓Calculations in HH |
| (Ando et al., 2023) | Non-randomised cross-over | NN, HH | 12 males, 23 ± 2 yr | 7 hr | NN = 0.21 $F_iO_2$<br>HH = 0.12 $F_iO_2$ | Go/No-Go<br>Memory task accuracy | ↓ Memory task accuracy in HH<br>↔ Go/No-Go RT & accuracy |
| (Asmaro et al., 2013) | Repeated measures within subject | NN, HH | 35 aviation industry employees, 19-69 yr | 30 min | NN = SL<br>HH1 = 5335m<br>HH2 = 7620m | ST<br>DST Forward & Back<br>TMT A & B | ↓ Performance during ST and DST Forward & Backward, & TMT A&B in HH |

| | | | | | | | |
|---|---|---|---|---|---|---|---|
| (Bailey et al., 2019) | Cross-sectional observational | - | 23 male highlander 17 male lowlanders | - | Highlanders = 3640m<br>Lowlanders = 80m | RAVL<br>RDF & RDB<br>TMT A & B<br>DSST<br>GPDT | ↓ RAVLT, RDB, RDF, TMT B, DSST, ST in highlanders compared to lowlanders |
| (Bakharev, 1981) | Repeated measures within subject | LA, HA | 7 young men | 8 days | HA = 3400-3600m | Visual, short term, long term, operative, figurative, associative, involuntary, and auditory memory (RAVLT-like) | ↓ Memory at day 2<br>Those of central Asia (highlanders) had less severe reductions compared to Pskov Region (lowlanders). |
| (Blogg & Gennser, 2006) | RCT | NN, HH | 13 subjects, 29.5 ± 6.4 | 10 min | 21kPa, 15kPa, 10kPa $P_iO_2$ | FCRT<br>Mannequin test<br>Finger-tapping test<br>Medium-tapping test | ↔ Psychomotor performance at 15kPa<br>↑ FCRT and medium-tapping test error at 10kPa. Although |

| | | | | | | | |
|---|---|---|---|---|---|---|---|
| | | | | | | | participants perceived being more focused |
| (Bloomfield et al., 2023) | RCT | NN, PCH, ICH | 12 adults, 26 ± 6.2 yr | 30 min | 10% oxygen gas (5500m) | CNSVS test battery | ↓ Neurocognitive index score, cognitive flexibility, complex attention, composite memory, and visual memory in PCH<br>↓ Neurocognitive index score, composite memory, psychomotor speed and ↑ RT in ICH |
| (Bonnon et al., 1995) | Comparative | NN, HH | 6 subject experimental and unspecified control group | 60 hr | 4383m | Cognitive-motor task | ↑ Cognitive-motor performance in control group<br>↔ Performance in experimental group comparable once at 60 hrs. |
| (Bonnon et al., 1999) | Comparative | NN, HH | 7 subject experimental and unspecified control group | 17 days | 9 days at 3500-4400m, 8 days climbing 500m/day to 5400m | Cognitive motor task | ↔ Difference between experimental and control group. |

| | | | | | | | |
|---|---|---|---|---|---|---|---|
| (Bouquet et al., 1999) | Comparative | NN, HH | 8 male climbers, 24 – 37 yr<br>10 controls, 22 – 40 yr | 31 days | Simulated climb to 8848m | Purdue pegboard, recognition task | ↓ Purdue pegboard performance > 8000m<br>↓Recognition performance in from 5500m<br>↓Performance in complex task vs simpler task |
| (Champod et al., 2013) | Non-randomised cross-over | NN, IH | 17 healthy males | 6 hrs of 1min on/1 min off | - | *n*-back task | ↑RT during spatial but not verbal n-back task in IH across all levels of difficulty |
| (Chroboczek et al., 2023) | Non-randomised cross-over | NN, NH | 17 healthy males | 15 min | NH = 0.13 $F_iO_2$ | ST | ↔ ST reading and naming interference |

| | | | | | | | |
|---|---|---|---|---|---|---|---|
| (Crow & Kelman, 1971) | Repeated measures within subject | HH | 86 medical students, 19 – 30 years. Organised into groups of 4 or 6 | 30 min | HH = 609m, 2438m, or 3656m | Required to memorise sequences of 6 digits at 1Hz (n-back-like task) to assess short-term memory | ↔ error rate between conditions |
| (Davis et al., 2015) | Comparative | HH | 132 residents into three groups based on altitude of residence (0-1500m, 1501-3000m, >3000m) | - | Testing occurred at 4860m | Go/no-go test score<br>VFT<br>Hand movement test | ↔ VFT<br>↓ Go/no-go in >3000m group<br>↓ Correct hand movements in >3000m and 1500-3000m group compared to 1500m group |
| (Falla, van Veelen, et al., 2024) | RCT | NN, HH | 48 pre-hospital emergency service personnel | 30 min | NN = 200m<br>HH = 3000m or 5000m | PVT<br>BART<br>DSST<br>VAS | ↓ PVT at 5000m compared to 200m, no difference in other test<br>Participants did not detect ↓ PVT on VAS |

| | | | | | | | |
|---|---|---|---|---|---|---|---|
| (Falla, Hüfner, et al., 2024) | RCT | HH | 36 pre-hospital emergency service personnel | 30 min | Two trials at 4000m, one with supplement oxygen and one without (control) | PVT<br>DSST<br>*n*-back<br>GPDT | ↑ PVT with oxygen supplementation ↑ 0.37ms in PVT each % reduction in $S_pO_2$. |
| (Gerard et al., 2000) | Randomised cross-over | NH | 23 healthy adults | 2 hr | NH = 5000m with and without 6% $O_2$ enrichment | Story Memory<br>TMT A&B<br>DSST<br>Finger Tapping<br>Digit Vigilance<br>GPDT<br>PASA<br>Matrix Reasoning<br>KSS<br>Probed Memory<br>Recall<br>PVT | ↓ RT in all tasks with $O_2$ enrichment<br>↑ Hand-eye coordination, and mood with $O_2$ enrichment. |

| | | | | | | | |
|---|---|---|---|---|---|---|---|
| | | | | | | CPT<br>CTT<br>PERS<br>VAS | |
| (Guo et al., 2017) | Observation | NN, HH | 163 healthy adults | 24 hr | HH = 3700m | RT<br>Attention<br>DSST<br>DST<br>Pursuit aiming | ↔ Cognitive measures between those with headache and those without at HA |
| (Hill et al., 2014) | Comparative | LA, HA | 229 in total: children (4-10 yr, LA = 44, HA = 52), adolescents (13-16 yr, LA = 23, HA = 25), young adults (25-40yr, LA = 20, HA = 20), older adults (>50 | - | LA = 500m<br>HA = 3700m | Cognitive Drug Research battery | ↓ Processing speed, attention in HA participants compared to LA. |

| | | | | | | | |
|---|---|---|---|---|---|---|---|
| | | | years, LA = 22, HA = 23) | | | | |
| (Hohenauer et al., 2022) | Randomised cross-over | NN, NH, HH | 20 healthy participants, 27.6 yr | NH = 15 min HH1 = 1hr HH2 = 24 hr | NN = 550m NH = .144 $F_iO_2$ HH = 2980m | ANAM | ↔ RT, logical relations, math processing, switching, and *n* back tests in any condition |
| (Keeler et al., 2023) | Repeated measures within subject | NN, NH | 21 healthy adults, 27 ± 5 yr | 90 min | NN = 0.206 $F_iO_2$ NH = 0.14 $F_iO_2$ | Gridshot Capacity Multitracker | ↔ no difference in any gamified task. |
| (Koller et al., 1993) | Comparative | NH | 10 acclimatised mountaineers and 11 | - | NH = 6000m | Attentiveness | ↓ Attentiveness in lowlanders compared to highlanders >4000m |

| | | | | | | | |
|---|---|---|---|---|---|---|---|
| | | | non-acclimatised controls | | | | |
| (Latshang et al., 2013) | Randomised cross-over | NN, HH | 51 healthy men, 24 yr | 2 days at each HH level | NN = 490m<br>HH1 = 1630m<br>HH2 = 2590m | PVT, DASS, *n*-back tasks | ↔ All cognitive tasks across conditions. |
| (Legg et al., 2016) | Randomised cross-over | NN, HH | 36 air force personnel | 30 min | HH1 = 8000ft<br>HH2 = 12000ft | ANAM | ↔ All cognitive tasks across conditions |
| (Malle et al., 2013) | Comparative | NN, HH | 28 pilots in HH and 29 control pilots | 90 min | NN = SL<br>HH = 10000m | PASA | ↓ Working memory in HH group compared to NN |
| (Merz et al., 2013) | Comparative | NN, HH | 32 mountaineers, separated into two groups summitting at different rates | - | NN = SL<br>HH1 = 4497m<br>HH2 = 5533m<br>HH3 = 6265m | LB<br>VFT<br>RFFT<br>CT | ↔ All cognitive tasks across conditions |

| | | | | | | | |
|---|---|---|---|---|---|---|---|
| (Niedermeier et al., 2017) | RCT | NN, IH, HH | 52 participants into a sham or IH pre-acclimatisation group | NN, IH = 1hr HH = 12 hr | NN = 0.2 $F_iO_2$<br>IH = 0.13 $F_iO_2$<br>HH = 0.13 $F_iO_2$ | Game of Dice Task | ↓ Risky decision making in HH for IH group compared to sham |
| (Noble et al., 1993) | Comparative | NN, NH | 2 groups of 12 male subjects | 30 min | NN = SL<br>NH = 78% $S_pO_2$ | TMT<br>DSST<br>LRT<br>RT | ↓ Performance of TMT and no learning effect in NH group compared to sham.<br>↑ RT |
| (Parker et al., 2017) | Comparative | NN, NH | 10 orthopaedic surgeons into either NN or NH group | 45 min | NN = SL<br>NH = 0.14 $F_iO_2$ | VWMC | ↔ VWMC tasks across conditions |
| (Pighin et al., 2014) | Randomised cross-over | NN, NH | 26 university students, 23 ± 6.8 yr | 60 min | NN = 0.21 $F_iO_2$<br>NH = 0.14 $F_iO_2$ | Decision-making task | ↓ Loss aversion in NH compared to NN. |

| | | | | | | | |
|---|---|---|---|---|---|---|---|
| (Quackenbush et al., 2016) | Comparative | NN, HH | 1 group performed VIHT<br>1 group performed sham | 30 min | HH1 = 3048m<br>HH2 = 3658m | ST<br>SDMT<br>DST | ↑ ST and SDMT performance for VIHT group compared to sham |
| (Ramírez-delaCruz et al., 2025) | Randomised cross-over | NN, NH | 19 participants, 23.7 ± 3.9 yr | 45 min | NN = SL<br>NH1=0.15 $F_iO_2$<br>NH2=0.13 $F_iO_2$<br>NH3=0.11 $F_iO_2$ | Memory task, Go/No-go task (sustained attention), Ericksen Flanker test, BAT (anticipation), RT | ↑ sustained attention at NH1,<br>↑ anticipation at NH1 & NH3,<br>↑ RT at NH3<br>↔ no difference across conditions |
| (Rice et al., 2019) | Repeated measures within subject | NN, NH | 60 aviation students | 5 min/level | NN = SL<br>NH1 = 2438m<br>NH2 = 3658m<br>NH3 = 4877m<br>NH4 = 6096m | Visual Sequence Comparison<br>X-Plane | ↓ Visual Sequency Comparison & X-Plane performance in NH compared to NN |

| | | | | | | | |
|---|---|---|---|---|---|---|---|
| (Schega et al., 2016) | RCT | NN, IH | 34 participants, 60 – 75 yr | 90 min | IH = 90-85% $S_pO_2$ | ST | ↑ ST performance in IH group |
| (Schega et al., 2013) | RCT | NN, IH | 34 participants, 60 – 75 yr | 90 min | IH = 90-85% $S_pO_2$ | ZVT | ↑ ZVT performance in IH group |
| (Schlaepfer et al., 1992) | Repeated measures within subject | NN, HH, NH | 10 healthy subjects | - | HH = 3450m<br>NH = 0.145 $F_iO_2$ | Read briefly displayed letters | ↓ time needed to read letters in HH and NH |
| (Shukitt-Hale et al., 1998) | Repeated measures within subject | NN, HH | 23 army personnel, 18-29 yr | 4.5 hr | NN = 550m<br>HH1 = 4200m<br>HH2 = 4700m | Battery of cognitive tasks | ↓ Performance in battery of cognitive tasks only after a few hours of exposure in a grade dependent fashion |
| (Tesler et al., 2015) | Randomised cross-over | NN, HH | 26 participants, 26.9 yr | 2 days | NN = 490m<br>HH1 = 1630m<br>HH2 = 2590m | Visio-motor learning task | ↓ Sleep dependent memory performance in HH conditions |

| | | | | | | | |
|---|---|---|---|---|---|---|---|
| (Thomas et al., 2007) | Prospective | HH | 11 subjects, 27 ± 1.5 yr | 9 hr | HH1 = 7700ft<br>HH2 = 10000ft<br>HH3 = 13000ft | PVT<br>*n*-back | ↔ No difference in PVT & *n*-back |
| (Turner, Byblow, et al., 2015) | Randomised match-paired | NN, NH | 22 adults, 23 yr | 50 min | NN = SL<br>NH = 0.1 $F_iO_2$ | CNS Vital Signs | ↓ Performance in CNS Vital Signs |
| (Turner, Barker-Collo, et al., 2015) | Randomised match-paired | NN, NH | 15 adults, 21-55 yr | 90 min | NN = SL<br>NH = 0.1 $F_iO_2$ | CNS Vital Signs | ↓ Performance in CNS Vital Signs |
| (van der Post et al., 2002) | Randomised cross-over | NN, NH | 12 healthy subjects | 2 hr | NN = 97% $S_pO_2$<br>HH1 = 90% $S_pO_2$<br>HH2 = 80% $S_pO_2$ | Memory task<br>RT | ↑ RT in HH2 compared to HH1 and NN.<br>↑ Cognitive performance in HH1 compared to NN and HH2 |

| | | | | | | | |
|---|---|---|---|---|---|---|---|
| (Van Dorp et al., 2007) | RCT | NH | 22 healthy subjects | 45 min | Sham hypoxia = 103 mmHg $PetO_2$ without inspired $CO_2$<br>Isocapnic hypoxia = 40mmHg with 0.4 mmHg clamped $CO_2$<br>Poikilocapnic hypoxia = 40mmHg without $CO_2$ | PVT and MAT-bat | ↓ Performance in PVT and MAT-bat |
| (Wu et al., 2023) | RCT | NN<br>HH | 82 males 20.76 ± 1.6, into control or a RIPC group | 6 days | HH = 3680m | WHO-NCTB | ↑ Spatial memory and digit symbol performance in RIPC group compared to controls<br>↑ Spatial memory performance in RIPC group compared to RIPC at baseline |

| | | | | | | | |
|---|---|---|---|---|---|---|---|
| (Yan et al., 2011a) | Comparative | NN HH | 2 groups: 1 born at SL, the other in HA | - | - | *2*-back task | ↔ Working memory performance |
| (Yan et al., 2011b) | Comparative | NN HH | 2 groups: 1 born at SL, the other in HA | - | - | Verbal working memory | ↑RT in HA group<br>↓ Accuracy in HA group |

*Green arrows represent positive outcomes, yellow arrows represent no change in outcomes, red arrows represent negative outcomes. NN, normobaric normoxia; HH, hypobaric hypoxia; NH, normobaric hypoxia; HN, hypobaric normoxia;* KLT, Konzentrations Leistungs Test-Revidierte Fassung; KSS, Karonlinska Sleepiness Scale; $F_iO_2$, fraction of inspired oxygen; RT, reaction time; RAVL, Rey Auditory Verbal Learning; RDF, Repetition of Digits Forward; RDB, Repetition of Digits Backwards test; TMT, Trail Making Test, DSST, Digit Symbol Substitution Test; ST, Stroop test; GPDT, grooved pegboard dexterity test; MoCA, Montreal Cognitive Assessment; BDI, Becks Depression Inventory; DST, Digit Span test; LA, low altitude; HA, high altitude; RCT, randomised control trial; $P_iO_2$, pressure of inspired oxygen; PCH, poikilocapnic hypoxia; ICH, isocapnic hypoxia; CNSVS, Central Nervous System Vital Signs; IH, intermittent hypoxia; VFT, verbal fluency test; BART, balloon analogue risk task; VAS, visual analogue scale; PASA, Paced Auditory Serial Addition; PVT, Psychomotor Vigilance Task; CPT, Calculation Performance Task; Compensatory Tracking Task; Performance Effort and Rating Scales; ANAM, Automated Neuropsychological Assessment Metrics; DASS, divided attention steering simulator; LB, line bisection test; RFFT, Ruff figural fluency test; CT, modified Pegboard test the Chess test; RBANS, The Repeatable Battery for the Assessment of Neuropsychological Status; WAIS-III, Wechsler Adult Intelligence Scale; RVP, Rapid Visual Information Processing test; OTS, One Touch Stockings of Cambridge; AMS, Acute Mountain Sickness; LRT, Letter Rotation Test; VWMC, verbal working memory capacity; VIHT, Voluntary Isocapnic Hyperventilation Training; SDMT, Symbol Digit Modalities Test; BAT, Bassin Anticipation Timer; MMSE, Mini Mental State Examination; ZVT, Number Combination Test; MAT-bat, Multi-Attribute Task battery; RIPC, remote ischemic precondition; WHO-NCTB, World Health Organization Neurobehavioral Core Test Battery; SL, sea level; FCRT, four choice reaction time.

*Table 2.* ROBINS-I-V2 risk of bias assessment for data items.

| **Data item** | **Domain 1** | **Domain 2** | **Domain 3** | **Domain 4** | **Domain 5** | **Domain 6** | **Domain 7** | **Overall** |
|---|---|---|---|---|---|---|---|---|
| *Chroboczek et al., 2023* | Serious | Low | Low | Low | Moderate | Low | Moderate | Serious |
| *Crow et al., 1971* | Moderate | Moderate | Moderate | Low | Low | Low | Moderate | Moderate |
| *Gerard et al., 2000* | Serious | Low | Low | Low | Serious | Low | Moderate | Serious |
| *Guo et al., 2017* | Serious | Low | Moderate | Low | Low | Serious | Moderate | Serious |
| *Quackenbush et al., 2016* | Moderate | Serious | Moderate | Serious | Low | Low | Moderate | Serious |
| *Van Dorp et al., 2007* | Serious | Low | Low | Low | Low | Low | Moderate | Serious |

*Domain 1, Bias due to confounding; Domain 2, Bias in the selection of participants; Domain 3, Bias in the classification of interventions; Domain 4, Bias due to deviations from intended interventions; Domain 5, Bias due to missing data; Domain 6, Bias in the measurement of outcomes; Domain 7, Bias in the selection of the reported result.*

*Table 3. List of confounding factors taken into consideration of data items assessed against Domain 1 of the ROBINS-I V2.*

| **Participant factors** | **Task factors** | **Environmental factors** | **Methodological factors** |
|---|---|---|---|
| - Differences in baseline cognitive performance between conditions<br>- Age variability<br>- Physical fitness variability<br>- Health conditions<br>- Acclimatisation<br>- Genetic factors | - Complexity of task<br>- Familiarity to task | - Severity and duration of hypoxia<br>- Simulation or environmental hypoxia<br>- Temperature and pressure | - Sensitivity and reliability of tools used<br>- Timing of assessments<br>- Control groups used |

### Figure 1

Forest plot of the overall effect of hypoxia on cognitive ability

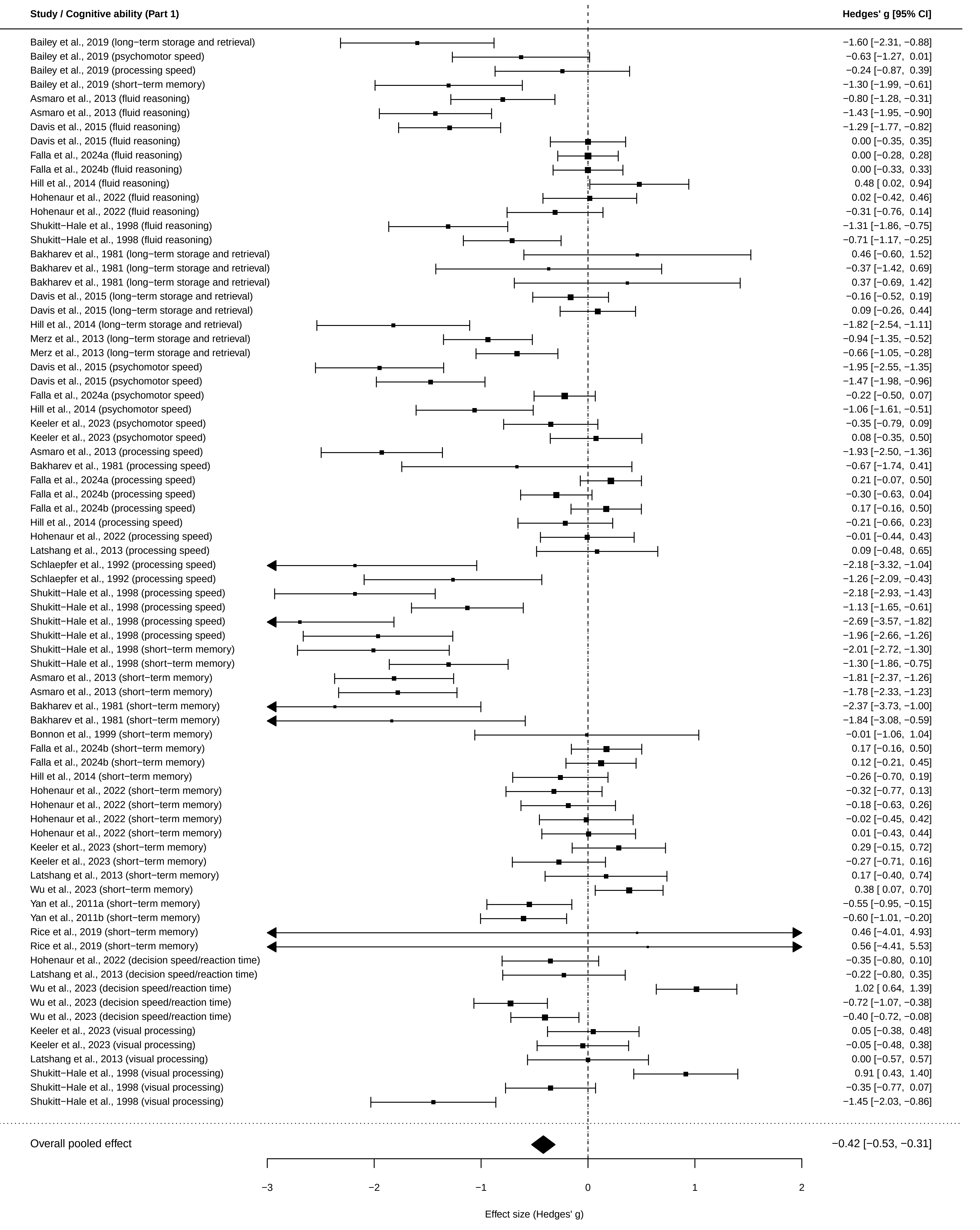
Study / Cognitive ability (Part 1)
Hedges' g [95% CI]
Bailey et al., 2019 (long−term storage and retrieval) −1.60 [−2.31, −0.88]
Bailey et al., 2019 (psychomotor speed) −0.63 [−1.27, 0.01]
Bailey et al., 2019 (processing speed) −0.24 [−0.87, 0.39]
Bailey et al., 2019 (short−term memory) −1.30 [−1.99, −0.61]
Asmaro et al., 2013 (fluid reasoning) −0.80 [−1.28, −0.31]
Asmaro et al., 2013 (fluid reasoning) −1.43 [−1.95, −0.90]
Davis et al., 2015 (fluid reasoning) −1.29 [−1.77, −0.82]
Davis et al., 2015 (fluid reasoning) 0.00 [−0.35, 0.35]
Falla et al., 2024a (fluid reasoning) 0.00 [−0.28, 0.28]
Falla et al., 2024b (fluid reasoning) 0.00 [−0.33, 0.33]
Hill et al., 2014 (fluid reasoning) 0.48 [ 0.02, 0.94]
Hohenaur et al., 2022 (fluid reasoning) 0.02 [−0.42, 0.46]
Hohenaur et al., 2022 (fluid reasoning) −0.31 [−0.76, 0.14]
Shukitt−Hale et al., 1998 (fluid reasoning) −1.31 [−1.86, −0.75]
Shukitt−Hale et al., 1998 (fluid reasoning) −0.71 [−1.17, −0.25]
Bakharev et al., 1981 (long−term storage and retrieval) 0.46 [−0.60, 1.52]
Bakharev et al., 1981 (long−term storage and retrieval) −0.37 [−1.42, 0.69]
Bakharev et al., 1981 (long−term storage and retrieval) 0.37 [−0.69, 1.42]
Davis et al., 2015 (long−term storage and retrieval) −0.16 [−0.52, 0.19]
Davis et al., 2015 (long−term storage and retrieval) 0.09 [−0.26, 0.44]
Hill et al., 2014 (long−term storage and retrieval) −1.82 [−2.54, −1.11]
Merz et al., 2013 (long−term storage and retrieval) −0.94 [−1.35, −0.52]
Merz et al., 2013 (long−term storage and retrieval) −0.66 [−1.05, −0.28]
Davis et al., 2015 (psychomotor speed) −1.95 [−2.55, −1.35]
Davis et al., 2015 (psychomotor speed) −1.47 [−1.98, −0.96]
Falla et al., 2024a (psychomotor speed) −0.22 [−0.50, 0.07]
Hill et al., 2014 (psychomotor speed) −1.06 [−1.61, −0.51]
Keeler et al., 2023 (psychomotor speed) −0.35 [−0.79, 0.09]
Keeler et al., 2023 (psychomotor speed) 0.08 [−0.35, 0.50]
Asmaro et al., 2013 (processing speed) −1.93 [−2.50, −1.36]
Bakharev et al., 1981 (processing speed) −0.67 [−1.74, 0.41]
Falla et al., 2024a (processing speed) 0.21 [−0.07, 0.50]
Falla et al., 2024b (processing speed) −0.30 [−0.63, 0.04]
Falla et al., 2024b (processing speed) 0.17 [−0.16, 0.50]
Hill et al., 2014 (processing speed) −0.21 [−0.66, 0.23]
Hohenaur et al., 2022 (processing speed) −0.01 [−0.44, 0.43]
Latshang et al., 2013 (processing speed) 0.09 [−0.48, 0.65]
Schlaepfer et al., 1992 (processing speed) −2.18 [−3.32, −1.04]
Schlaepfer et al., 1992 (processing speed) −1.26 [−2.09, −0.43]
Shukitt−Hale et al., 1998 (processing speed) −2.18 [−2.93, −1.43]
Shukitt−Hale et al., 1998 (processing speed) −1.13 [−1.65, −0.61]
Shukitt−Hale et al., 1998 (processing speed) −2.69 [−3.57, −1.82]
Shukitt−Hale et al., 1998 (processing speed) −1.96 [−2.66, −1.26]
Shukitt−Hale et al., 1998 (short−term memory) −2.01 [−2.72, −1.30]
Shukitt−Hale et al., 1998 (short−term memory) −1.30 [−1.86, −0.75]
Asmaro et al., 2013 (short−term memory) −1.81 [−2.37, −1.26]
Asmaro et al., 2013 (short−term memory) −1.78 [−2.33, −1.23]
Bakharev et al., 1981 (short−term memory) −2.37 [−3.73, −1.00]
Bakharev et al., 1981 (short−term memory) −1.84 [−3.08, −0.59]
Bonnon et al., 1999 (short−term memory) −0.01 [−1.06, 1.04]
Falla et al., 2024b (short−term memory) 0.17 [−0.16, 0.50]
Falla et al., 2024b (short−term memory) 0.12 [−0.21, 0.45]
Hill et al., 2014 (short−term memory) −0.26 [−0.70, 0.19]
Hohenaur et al., 2022 (short−term memory) −0.32 [−0.77, 0.13]
Hohenaur et al., 2022 (short−term memory) −0.18 [−0.63, 0.26]
Hohenaur et al., 2022 (short−term memory) −0.02 [−0.45, 0.42]
Hohenaur et al., 2022 (short−term memory) 0.01 [−0.43, 0.44]
Keeler et al., 2023 (short−term memory) 0.29 [−0.15, 0.72]
Keeler et al., 2023 (short−term memory) −0.27 [−0.71, 0.16]
Latshang et al., 2013 (short−term memory) 0.17 [−0.40, 0.74]
Wu et al., 2023 (short−term memory) 0.38 [ 0.07, 0.70]
Yan et al., 2011a (short−term memory) −0.55 [−0.95, −0.15]
Yan et al., 2011b (short−term memory) −0.60 [−1.01, −0.20]
Rice et al., 2019 (short−term memory) 0.46 [−4.01, 4.93]
Rice et al., 2019 (short−term memory) 0.56 [−4.41, 5.53]
Hohenaur et al., 2022 (decision speed/reaction time) −0.35 [−0.80, 0.10]
Latshang et al., 2013 (decision speed/reaction time) −0.22 [−0.80, 0.35]
Wu et al., 2023 (decision speed/reaction time) 1.02 [ 0.64, 1.39]
Wu et al., 2023 (decision speed/reaction time) −0.72 [−1.07, −0.38]
Wu et al., 2023 (decision speed/reaction time) −0.40 [−0.72, −0.08]
Keeler et al., 2023 (visual processing) 0.05 [−0.38, 0.48]
Keeler et al., 2023 (visual processing) −0.05 [−0.48, 0.38]
Latshang et al., 2013 (visual processing) 0.00 [−0.57, 0.57]
Shukitt−Hale et al., 1998 (visual processing) 0.91 [ 0.43, 1.40]
Shukitt−Hale et al., 1998 (visual processing) −0.35 [−0.77, 0.07]
Shukitt−Hale et al., 1998 (visual processing) −1.45 [−2.03, −0.86]
Overall pooled effect −0.42 [−0.53, −0.31]
−3
−2
−1
0
1
2
Effect size (Hedges' g)

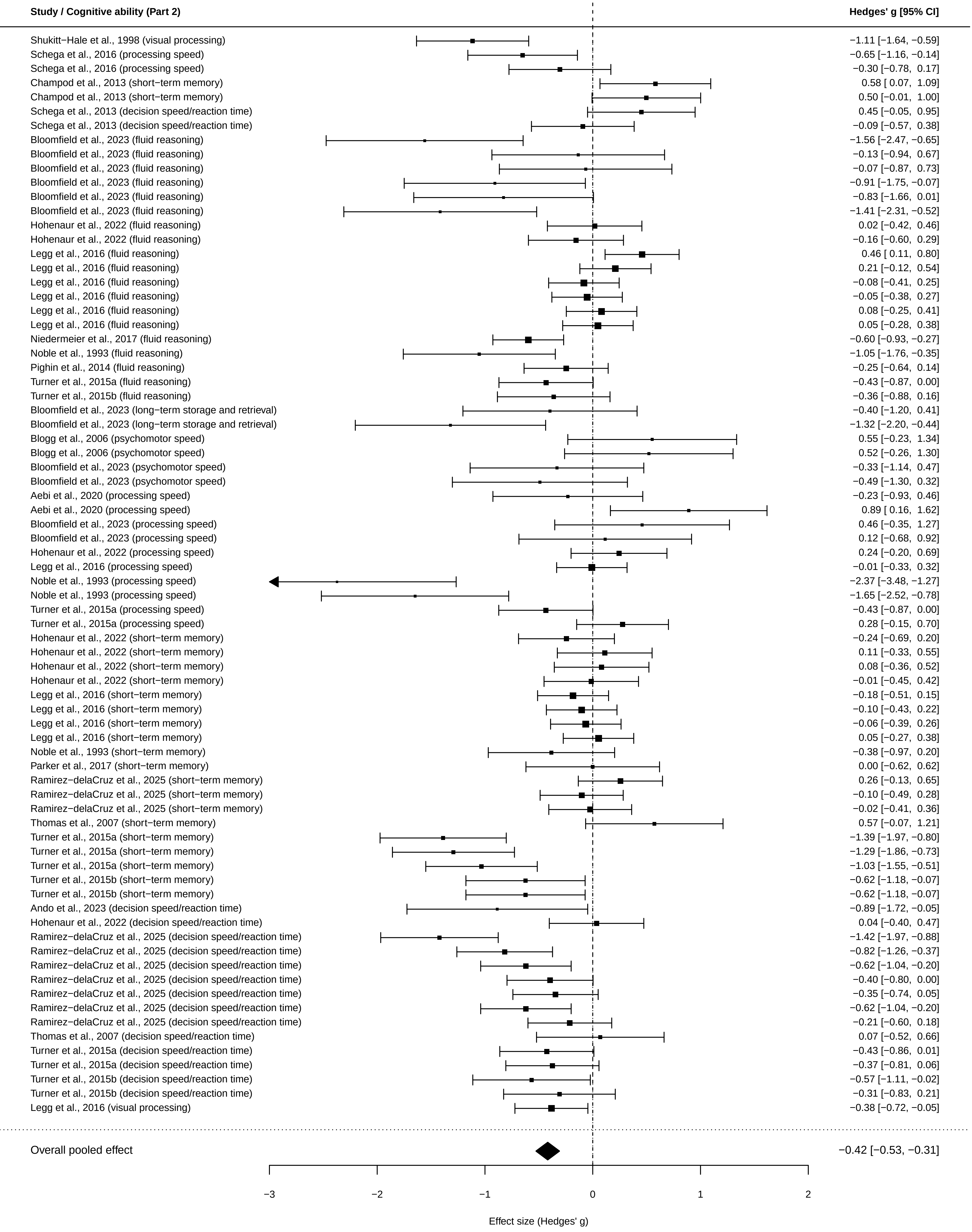

Study / Cognitive ability (Part 2)
Hedges' g [95% CI]
Shukitt−Hale et al., 1998 (visual processing) −1.11 [−1.64, −0.59]
Schega et al., 2016 (processing speed) −0.65 [−1.16, −0.14]
Schega et al., 2016 (processing speed) −0.30 [−0.78, 0.17]
Champod et al., 2013 (short−term memory) 0.58 [ 0.07, 1.09]
Champod et al., 2013 (short−term memory) 0.50 [−0.01, 1.00]
Schega et al., 2013 (decision speed/reaction time) 0.45 [−0.05, 0.95]
Schega et al., 2013 (decision speed/reaction time) −0.09 [−0.57, 0.38]
Bloomfield et al., 2023 (fluid reasoning) −1.56 [−2.47, −0.65]
Bloomfield et al., 2023 (fluid reasoning) −0.13 [−0.94, 0.67]
Bloomfield et al., 2023 (fluid reasoning) −0.07 [−0.87, 0.73]
Bloomfield et al., 2023 (fluid reasoning) −0.91 [−1.75, −0.07]
Bloomfield et al., 2023 (fluid reasoning) −0.83 [−1.66, 0.01]
Bloomfield et al., 2023 (fluid reasoning) −1.41 [−2.31, −0.52]
Hohenaur et al., 2022 (fluid reasoning) 0.02 [−0.42, 0.46]
Hohenaur et al., 2022 (fluid reasoning) −0.16 [−0.60, 0.29]
Legg et al., 2016 (fluid reasoning) 0.46 [ 0.11, 0.80]
Legg et al., 2016 (fluid reasoning) 0.21 [−0.12, 0.54]
Legg et al., 2016 (fluid reasoning) −0.08 [−0.41, 0.25]
Legg et al., 2016 (fluid reasoning) −0.05 [−0.38, 0.27]
Legg et al., 2016 (fluid reasoning) 0.08 [−0.25, 0.41]
Legg et al., 2016 (fluid reasoning) 0.05 [−0.28, 0.38]
Niedermeier et al., 2017 (fluid reasoning) −0.60 [−0.93, −0.27]
Noble et al., 1993 (fluid reasoning) −1.05 [−1.76, −0.35]
Pighin et al., 2014 (fluid reasoning) −0.25 [−0.64, 0.14]
Turner et al., 2015a (fluid reasoning) −0.43 [−0.87, 0.00]
Turner et al., 2015b (fluid reasoning) −0.36 [−0.88, 0.16]
Bloomfield et al., 2023 (long−term storage and retrieval) −0.40 [−1.20, 0.41]
Bloomfield et al., 2023 (long−term storage and retrieval) −1.32 [−2.20, −0.44]
Blogg et al., 2006 (psychomotor speed) 0.55 [−0.23, 1.34]
Blogg et al., 2006 (psychomotor speed) 0.52 [−0.26, 1.30]
Bloomfield et al., 2023 (psychomotor speed) −0.33 [−1.14, 0.47]
Bloomfield et al., 2023 (psychomotor speed) −0.49 [−1.30, 0.32]
Aebi et al., 2020 (processing speed) −0.23 [−0.93, 0.46]
Aebi et al., 2020 (processing speed) 0.89 [ 0.16, 1.62]
Bloomfield et al., 2023 (processing speed) 0.46 [−0.35, 1.27]
Bloomfield et al., 2023 (processing speed) 0.12 [−0.68, 0.92]
Hohenaur et al., 2022 (processing speed) 0.24 [−0.20, 0.69]
Legg et al., 2016 (processing speed) −0.01 [−0.33, 0.32]
Noble et al., 1993 (processing speed) −2.37 [−3.48, −1.27]
Noble et al., 1993 (processing speed) −1.65 [−2.52, −0.78]
Turner et al., 2015a (processing speed) −0.43 [−0.87, 0.00]
Turner et al., 2015a (processing speed) 0.28 [−0.15, 0.70]
Hohenaur et al., 2022 (short−term memory) −0.24 [−0.69, 0.20]
Hohenaur et al., 2022 (short−term memory) 0.11 [−0.33, 0.55]
Hohenaur et al., 2022 (short−term memory) 0.08 [−0.36, 0.52]
Hohenaur et al., 2022 (short−term memory) −0.01 [−0.45, 0.42]
Legg et al., 2016 (short−term memory) −0.18 [−0.51, 0.15]
Legg et al., 2016 (short−term memory) −0.10 [−0.43, 0.22]
Legg et al., 2016 (short−term memory) −0.06 [−0.39, 0.26]
Legg et al., 2016 (short−term memory) 0.05 [−0.27, 0.38]
Noble et al., 1993 (short−term memory) −0.38 [−0.97, 0.20]
Parker et al., 2017 (short−term memory) 0.00 [−0.62, 0.62]
Ramirez−delaCruz et al., 2025 (short−term memory) 0.26 [−0.13, 0.65]
Ramirez−delaCruz et al., 2025 (short−term memory) −0.10 [−0.49, 0.28]
Ramirez−delaCruz et al., 2025 (short−term memory) −0.02 [−0.41, 0.36]
Thomas et al., 2007 (short−term memory) 0.57 [−0.07, 1.21]
Turner et al., 2015a (short−term memory) −1.39 [−1.97, −0.80]
Turner et al., 2015a (short−term memory) −1.29 [−1.86, −0.73]
Turner et al., 2015a (short−term memory) −1.03 [−1.55, −0.51]
Turner et al., 2015b (short−term memory) −0.62 [−1.18, −0.07]
Turner et al., 2015b (short−term memory) −0.62 [−1.18, −0.07]
Ando et al., 2023 (decision speed/reaction time) −0.89 [−1.72, −0.05]
Hohenaur et al., 2022 (decision speed/reaction time) 0.04 [−0.40, 0.47]
Ramirez−delaCruz et al., 2025 (decision speed/reaction time) −1.42 [−1.97, −0.88]
Ramirez−delaCruz et al., 2025 (decision speed/reaction time) −0.82 [−1.26, −0.37]
Ramirez−delaCruz et al., 2025 (decision speed/reaction time) −0.62 [−1.04, −0.20]
Ramirez−delaCruz et al., 2025 (decision speed/reaction time) −0.40 [−0.80, 0.00]
Ramirez−delaCruz et al., 2025 (decision speed/reaction time) −0.35 [−0.74, 0.05]
Ramirez−delaCruz et al., 2025 (decision speed/reaction time) −0.62 [−1.04, −0.20]
Ramirez−delaCruz et al., 2025 (decision speed/reaction time) −0.21 [−0.60, 0.18]
Thomas et al., 2007 (decision speed/reaction time) 0.07 [−0.52, 0.66]
Turner et al., 2015a (decision speed/reaction time) −0.43 [−0.86, 0.01]
Turner et al., 2015a (decision speed/reaction time) −0.37 [−0.81, 0.06]
Turner et al., 2015b (decision speed/reaction time) −0.57 [−1.11, −0.02]
Turner et al., 2015b (decision speed/reaction time) −0.31 [−0.83, 0.21]
Legg et al., 2016 (visual processing) −0.38 [−0.72, −0.05]
Overall pooled effect −0.42 [−0.53, −0.31]
−3
−2
−1
0
1
2
Effect size (Hedges' g)
