## Supplementary Material 2 for "Neurocognitive Vulnerability to Hypoxia Across Cognitive Domains: A Systematic Review and Meta-Analysis"

### PRISMA 2020 Checklist

| Section and Topic | Item # | Checklist item | Location where item is reported |
|---|---|---|---|
| **TITLE** | | | |
| Title | 1 | Identify the report as a systematic review. | Page 1 |
| **ABSTRACT** | | | |
| Abstract | 2 | See the PRISMA 2020 for Abstracts checklist. | Page 2 |
| **INTRODUCTION** | | | |
| Rationale | 3 | Describe the rationale for the review in the context of existing knowledge. | Page 3 - 6 |
| Objectives | 4 | Provide an explicit statement of the objective(s) or question(s) the review addresses. | Page 6 |
| **METHODS** | | | |
| Eligibility criteria | 5 | Specify the inclusion and exclusion criteria for the review and how studies were grouped for the syntheses. | Page 16 |
| Information sources | 6 | Specify all databases, registers, websites, organisations, reference lists and other sources searched or consulted to identify studies. Specify the date when each source was last searched or consulted. | Page 16 - 17 |
| Search strategy | 7 | Present the full search strategies for all databases, registers and websites, including any filters and limits used. | Page 17 |
| Selection process | 8 | Specify the methods used to decide whether a study met the inclusion criteria of the review, including how many reviewers screened each record and each report retrieved, whether they worked independently, and if applicable, details of automation tools used in the process. | Page 17 |
| Data collection process | 9 | Specify the methods used to collect data from reports, including how many reviewers collected data from each report, whether they worked independently, any processes for obtaining or confirming data from study investigators, and if applicable, details of automation tools used in the process. | Page 17 |
| Data items | 10a | List and define all outcomes for which data were sought. Specify whether all results that were compatible with each outcome domain in each study were sought (e.g. for all measures, time points, analyses), and if not, the methods used to decide which results to collect. | Page 17 |
| | 10b | List and define all other variables for which data were sought (e.g. participant and intervention characteristics, funding sources). Describe any assumptions made about any missing or unclear information. | Page 18 |
| Study risk of bias assessment | 11 | Specify the methods used to assess risk of bias in the included studies, including details of the tool(s) used, how many reviewers assessed each study and whether they worked independently, and if applicable, details of automation tools used in the process. | Page 18 |
| Effect measures | 12 | Specify for each outcome the effect measure(s) (e.g. risk ratio, mean difference) used in the synthesis or presentation of results. | Page 18 - 19 |
| Synthesis methods | 13a | Describe the processes used to decide which studies were eligible for each synthesis (e.g. tabulating the study intervention characteristics and comparing against the planned groups for each synthesis (item #5)). | Page 18 - 19 |
| | 13b | Describe any methods required to prepare the data for presentation or synthesis, such as handling of missing summary statistics, or data conversions. | Page 18 - 19 |
| | 13c | Describe any methods used to tabulate or visually display results of individual studies and syntheses. | Page 18 - 19 |
| | 13d | Describe any methods used to synthesize results and provide a rationale for the choice(s). If meta-analysis was performed, describe the model(s), method(s) to identify the presence and extent of statistical heterogeneity, and software package(s) used. | Page 18 - 19 |
| | 13e | Describe any methods used to explore possible causes of heterogeneity among study results (e.g. subgroup analysis, meta-regression). | Page 18 - 19 |
| | 13f | Describe any sensitivity analyses conducted to assess robustness of the synthesized results. | Page 18 - 19 |
| Reporting bias assessment | 14 | Describe any methods used to assess risk of bias due to missing results in a synthesis (arising from reporting biases). | Page 18 |
| Certainty assessment | 15 | Describe any methods used to assess certainty (or confidence) in the body of evidence for an outcome. | Page 18 |

PRISMA

### PRISMA 2020 Checklist

| Section and Topic | Item # | Checklist item | Location where item is reported |
|---|---|---|---|
| **RESULTS** | | | |
| Study selection | 16a | Describe the results of the search and selection process, from the number of records identified in the search to the number of studies included in the review, ideally using a flow diagram. | Supplementary Material |
| | 16b | Cite studies that might appear to meet the inclusion criteria, but which were excluded, and explain why they were excluded. | Supplementary Material |
| Study characteristics | 17 | Cite each included study and present its characteristics. | Supplementary Material |
| Risk of bias in studies | 18 | Present assessments of risk of bias for each included study. | Supplementary Material |
| Results of individual studies | 19 | For all outcomes, present, for each study: (a) summary statistics for each group (where appropriate) and (b) an effect estimate and its precision (e.g. confidence/credible interval), ideally using structured tables or plots. | Page 6 – 9 |
| Results of syntheses | 20a | For each synthesis, briefly summarise the characteristics and risk of bias among contributing studies. | Page 6 – 9 |
| | 20b | Present results of all statistical syntheses conducted. If meta-analysis was done, present for each the summary estimate and its precision (e.g. confidence/credible interval) and measures of statistical heterogeneity. If comparing groups, describe the direction of the effect. | Page 6 – 9 |
| | 20c | Present results of all investigations of possible causes of heterogeneity among study results. | Page 6 – 9 |
| | 20d | Present results of all sensitivity analyses conducted to assess the robustness of the synthesized results. | Page 6 – 9 |
| Reporting biases | 21 | Present assessments of risk of bias due to missing results (arising from reporting biases) for each synthesis assessed. | Page 6 – 9 |
| Certainty of evidence | 22 | Present assessments of certainty (or confidence) in the body of evidence for each outcome assessed. | Page 18 |
| **DISCUSSION** | | | |
| Discussion | 23a | Provide a general interpretation of the results in the context of other evidence. | Page 10 – 16 |
| | 23b | Discuss any limitations of the evidence included in the review. | Page 15 – 16 |
| | 23c | Discuss any limitations of the review processes used. | Page 15 – 16 |
| | 23d | Discuss implications of the results for practice, policy, and future research. | Page 16 |
| **OTHER INFORMATION** | | | |
| Registration and protocol | 24a | Provide registration information for the review, including register name and registration number, or state that the review was not registered. | Page 16 |
| | 24b | Indicate where the review protocol can be accessed, or state that a protocol was not prepared. | Page 16 |
| | 24c | Describe and explain any amendments to information provided at registration or in the protocol. | NA |
| Support | 25 | Describe sources of financial or non-financial support for the review, and the role of the funders or sponsors in the review. | Page 20 |
| Competing interests | 26 | Declare any competing interests of review authors. | Page 20 |
| Availability of data, code and other materials | 27 | Report which of the following are publicly available and where they can be found: template data collection forms; data extracted from included studies; data used for all analyses; analytic code; any other materials used in the review. | Page 20 |

*From:* Page MJ, McKenzie JE, Bossuyt PM, Boutron I, Hoffmann TC, Mulrow CD, et al. The PRISMA 2020 statement: an updated guideline for reporting systematic reviews. BMJ 2021;372:n71. doi: 10.1136/bmj.n71.
This work is licensed under CC BY 4.0. To view a copy of this license, visit https://creativecommons.org/licenses/by/4.0/

**Supplementary Table S1. Database Search Strategies**

The following electronic database search strategies were used to identify studies investigating the effects of hypoxia on cognition. Searches were conducted on 6 January 2025 and were limited to studies involving humans published in English.

**PubMed**

**Search date: 6 January 2025**

("Hypoxia"[Title/Abstract]
OR "Altitude"[Title/Abstract]
OR "Hypobaric"[Title/Abstract])
AND
("Cognition"[Title/Abstract]
OR "Attention"[Title/Abstract]
OR "Memory"[Title/Abstract]
OR "Executive Function"[Title/Abstract]
OR "Processing Speed"[Title/Abstract]
OR "Psychomotor Speed"[Title/Abstract])
Filters: Humans; English

**Scopus**

**Search date: 6 January 2025**

TITLE-ABS-KEY
(
(hypoxia
OR altitude
OR hypobaric)
AND
(cognition
OR attention
OR memory
OR "executive function"
OR "processing speed"
OR "psychomotor speed")
)
AND (LIMIT-TO(LANGUAGE,"English"))

**Medline**

**Search date: 6 January 2025**

TS=

(
(hypoxia
OR altitude
OR hypobaric)
AND
(cognition
OR attention
OR memory
OR "executive function"
OR "processing speed"
OR "psychomotor speed")
)
Refined by:
Languages = English

**Cochrane Library**

**Search date: 6 January 2025**

(
hypoxia
OR altitude
OR hypobaric
)
AND
(
cognition
OR attention
OR memory
OR "executive function"
OR "processing speed"
OR "psychomotor speed"
)
Limiters:
Human
English Language

#### Excluded References


(CDC), C. f. D. C. a. P. (2002). Respiratory illness in workers exposed to metalworking fluid contaminated with nontuberculous mycobacteria--Ohio, 2001. *MMWR Morb Mortal Wkly Rep*, *51*(16), 349-352.

Abbasi, A. A., & Harrop, N. S. (2005). An unusual swelling in the neck. *Emerg Med J*, *22*(9), 674-675. https://doi.org/10.1136/emj.2004.014613

Abbate, C., Arosio, B., Galimberti, D., Nicolini, P., Chiara, L. R., Rossi, P. D., Ferri, E., Gussago, C., Deriz, M., Fenoglio, C., Serpente, M., Scarpini, E., & Mari, D. (2014). Phenotypic variability associated with the C9ORF72 hexanucleotide repeat expansion: a sporadic case of frontotemporal lobar degeneration with prodromal hyposmia and predominant semantic deficits. *J Alzheimers Dis*, *40*(4), 849-855. https://doi.org/10.3233/jad-132075

Adams, L., Chronos, N., Lane, R., & Guz, A. (1985). The measurement of breathlessness induced in normal subjects: validity of two scaling techniques. *Clin Sci (Lond)*, *69*(1), 7-16. https://doi.org/10.1042/cs0690007

Adejuyigbe, E. A., Agyeman, I., Anand, P., Anyabolu, H. C., Arya, S., Assenga, E. N., Badhal, S., Brobby, N. W., Chellani, H. K., Chopra, N., Debata, P. K., Dube, Q., Dua, T., Gadama, L., Gera, R., Hammond, C. K., Jain, S., Kantumbiza, F., Kawaza, K., . . . Yiadom, A. B. (2023). Evaluation of the impact of continuous Kangaroo Mother Care (KMC) initiated immediately after birth compared to KMC initiated after stabilization in newborns with birth weight 1.0 to < 1.8 kg on neurodevelopmental outcomes: Protocol for a follow-up study. *Trials*, *24*(1), 265. https://doi.org/10.1186/s13063-023-07192-5

Aguirre, J. A., Etzensperger, F., Brada, M., Guzzella, S., Saporito, A., Blumenthal, S., Bühler, P., & Borgeat, A. (2019). The beach chair position for shoulder surgery in intravenous general anesthesia and controlled hypotension: Impact on cerebral oxygenation, cerebral blood flow and neurobehavioral outcome. *J Clin Anesth*, *53*, 40-48. https://doi.org/10.1016/j.jclinane.2018.09.035

Ahmed, H. A., Ishrat, T., Pillai, B., Fouda, A. Y., Sayed, M. A., Eldahshan, W., Waller, J. L., Ergul, A., & Fagan, S. C. (2018). RAS modulation prevents progressive cognitive impairment after experimental stroke: a randomized, blinded preclinical trial. *J Neuroinflammation*, *15*(1), 229. https://doi.org/10.1186/s12974-018-1262-x

Ahmed, M., & Solela, G. (2024). Thrombus in transit associated with fatal pulmonary thromboembolism in an elderly Ethiopian man following a surgical procedure: A case report. *Clin Case Rep*, *12*(8), e9293. https://doi.org/10.1002/ccr3.9293

Ai, J., & Baker, A. (2006). Long-term potentiation of evoked presynaptic response at CA3-CA1 synapses by transient oxygen-glucose deprivation in rat brain slices. *Exp Brain Res*, *169*(1), 126-129. https://doi.org/10.1007/s00221-005-0314-5

Akça, O., & Sessler, D. I. (2002). Use of cerebral oximetry to detect and manage cerebral desaturation with a rapidly expanding neck hematoma. *Acta Anaesthesiol Scand*, *46*(5), 607-608. https://doi.org/10.1034/j.1399-6576.2002.460521.x

Aker, K., Thomas, N., Adde, L., Koshy, B., Martinez-Biarge, M., Nakken, I., Padankatti, C. S., & Støen, R. (2022). Prediction of outcome from MRI and general movements assessment after hypoxic-ischaemic encephalopathy in low-income and middle-income countries: data from a randomised controlled trial. *Arch Dis Child Fetal Neonatal Ed*, *107*(1), 32-38. https://doi.org/10.1136/archdischild-2020-321309

Akoh, C. C., Schick, C., Otero, J., & Karam, M. (2014). Fat embolism syndrome after femur fracture fixation: a case report. *Iowa Orthop J*, *34*, 55-62.

Alali, A. S., Temkin, N., Vavilala, M. S., Lele, A. V., Barber, J., Dikmen, S., & Chesnut, R. M. (2020). Matching early arterial oxygenation to long-term outcome in severe traumatic brain injury: target values. *J Neurosurg*, *132*(2), 537-544. https://doi.org/10.3171/2018.10.Jns18964

Alberti, A., Valenti, S., Gallo, F., Petolillo, M., & Del Monte, D. (1994). Acute buflomedil intoxication: a life-threatening condition. *Intensive Care Med*, *20*(3), 219-221. https://doi.org/10.1007/bf01704705

Alchanatis, M., Zias, N., Deligiorgis, N., Liappas, I., Chroneou, A., Soldatos, C., & Roussos, C. (2008). Comparison of cognitive performance among different age groups in patients with obstructive sleep apnea. *Sleep Breath*, *12*(1), 17-24. https://doi.org/10.1007/s11325-007-0133-y

Alexander, M. P. (1997). Specific semantic memory loss after hypoxic-ischemic injury. *Neurology*, *48*(1), 165-173. https://doi.org/10.1212/wnl.48.1.165

Ali, K., Warusevitane, A., Lally, F., Sim, J., Sills, S., Pountain, S., Nevatte, T., Allen, M., & Roffe, C. (2014). The stroke oxygen pilot study: a randomized controlled trial of the effects of routine oxygen supplementation early after acute stroke--effect on key outcomes at six months. *PLoS One*, *8*(6), e59274. https://doi.org/10.1371/journal.pone.0059274

Alimović, S., Jurić, N., & Bošnjak, V. M. (2014). Functional vision in children with perinatal brain damage. *J Matern Fetal Neonatal Med*, *27*(14), 1491-1494. https://doi.org/10.3109/14767058.2013.863863

Allado, E., Chenuel, B., Vauthier, J. C., Hily, O., Richard, S., & Poussel, M. (2024). Transient Central Facial Palsy at High Altitude: A Case Report. *High Alt Med Biol*, *25*(1), 100-102. https://doi.org/10.1089/ham.2020.0184

Allen, J. S., Tranel, D., Bruss, J., & Damasio, H. (2006). Correlations between regional brain volumes and memory performance in anoxia. *J Clin Exp Neuropsychol*, *28*(4), 457-476. https://doi.org/10.1080/13803390590949287

Alvarez, F. J., Alvarez, A. A., Rodríguez, J. J., Lafuente, H., Canduela, M. J., Hind, W., Blanco-Bruned, J. L., Alonso-Alconada, D., & Hilario, E. (2023). Effects of Cannabidiol, Hypothermia, and Their Combination in Newborn Rats with Hypoxic-Ischemic Encephalopathy. *eNeuro*, *10*(5). https://doi.org/10.1523/eneuro.0417-22.2023

Amicuzi, I., Cappelli, F., Stortini, M., Cherubini, S., & Pierro, M. M. (2005). Follow-up of neuropsychological function recovery in a 9-year-old girl with anoxic encephalopathy: a window on the brain re-organization processes. *Brain Inj*, *19*(5), 371-388. https://doi.org/10.1080/02699050400004286

Anderson, J. E., & Robichaud, R. C. (1975). Retrograde amnesia induced by hypoxia and electroconvulsive shock in two rat strains. *Physiol Behav*, *14*(1), 81-84. https://doi.org/10.1016/0031-9384(75)90145-6

André, C., Rehel, S., Kuhn, E., Landeau, B., Moulinet, I., Touron, E., Ourry, V., Le Du, G., Mézenge, F., Tomadesso, C., de Flores, R., Bejanin, A., Sherif, S., Delcroix, N., Manrique, A., Abbas, A., Marchant, N. L., Lutz, A., Klimecki, O. M., . . . Rauchs, G. (2020). Association of Sleep-Disordered Breathing With Alzheimer Disease Biomarkers in Community-Dwelling Older Adults: A Secondary Analysis of a Randomized Clinical Trial. *JAMA Neurol*, *77*(6), 716-724. https://doi.org/10.1001/jamaneurol.2020.0311

Andrewes, D. (1989). Management of disruptive behaviour in the brain-damaged patient using selective reinforcement. *J Behav Ther Exp Psychiatry*, *20*(3), 261-264. https://doi.org/10.1016/0005-7916(89)90032-3

Annink, K. V., de Vries, L. S., Groenendaal, F., Eijsermans, R., Mocking, M., van Schooneveld, M. M. J., Dudink, J., van Straaten, H. L. M., Benders, M., Lequin, M., & van der Aa, N. E. (2021). Mammillary body atrophy and other MRI correlates of school-age outcome following neonatal hypoxic-ischemic encephalopathy. *Sci Rep*, *11*(1), 5017. https://doi.org/10.1038/s41598-021-83982-8

Annoni, J. M., Vuagnat, H., Frischknecht, R., & Uebelhart, D. (1998). Percutaneous endoscopic gastrostomy in neurological rehabilitation: a report of six cases. *Disabil Rehabil*, *20*(8), 308-314. https://doi.org/10.3109/09638289809166086

Anobile, G., Tomaiuolo, F., Campana, S., & Cicchini, G. M. (2020). Three-systems for visual

numerosity: A single case study. *Neuropsychologia*, *136*, 107259. https://doi.org/10.1016/j.neuropsychologia.2019.107259
Antonelli Incalzi, R., Marra, C., Giordano, A., Calcagni, M. L., Cappa, A., Basso, S., Pagliari, G., & Fuso, L. (2003). Cognitive impairment in chronic obstructive pulmonary disease--a neuropsychological and spect study. *J Neurol*, *250*(3), 325-332. https://doi.org/10.1007/s00415-003-1005-4
Arai, Y., Shitoto, K., Muta, T., & Kurosawa, H. (1999). Pulmonary thromboembolism after spinal instrumentation surgery. *J Orthop Sci*, *4*(5), 380-383. https://doi.org/10.1007/s007760050120
Arbelaez, A., Castillo, M., & Mukherji, S. K. (1999). Diffusion-weighted MR imaging of global cerebral anoxia. *AJNR Am J Neuroradiol*, *20*(6), 999-1007.
Arciniegas, D. B., Frey, K. L., Anderson, C. A., Brousseau, K. M., & Harris, S. N. (2004). Amantadine for neurobehavioural deficits following delayed post-hypoxic encephalopathy. *Brain Inj*, *18*(12), 1309-1318. https://doi.org/10.1080/02699050410001720130
Arias-Cavieres, A., & Garcia, A. J., 3rd. (2023). A Consequence of Immature Breathing induces Persistent Changes in Hippocampal Synaptic Plasticity and Behavior: A Role of Pro-Oxidant State and NMDA Receptor Imbalance. *bioRxiv*. https://doi.org/10.1101/2023.03.21.533692
Armentrout, J. J., Holland, D. A., O'Toole, K. J., & Ercoline, W. R. (2006). Fatigue and related human factors in the near crash of a large military aircraft. *Aviat Space Environ Med*, *77*(9), 963-970.
Arpesella, R., Dallocchio, C., Arbasino, C., Imberti, R., Martinotti, R., & Frucht, S. J. (2009). A patient with intractable posthypoxic myoclonus (Lance-Adams syndrome) treated with sodium oxybate. *Anaesth Intensive Care*, *37*(2), 314-318. https://doi.org/10.1177/0310057x0903700214
Arteni, N. S., Salgueiro, J., Torres, I., Achaval, M., & Netto, C. A. (2003). Neonatal cerebral hypoxia-ischemia causes lateralized memory impairments in the adult rat. *Brain Res*, *973*(2), 171-178. https://doi.org/10.1016/s0006-8993(03)02436-3
Ashwal, S., & Cranford, R. (2002). The minimally conscious state in children. *Semin Pediatr Neurol*, *9*(1), 19-34. https://doi.org/10.1053/spen.2002.30334
Atluri, P., Vasireddy, D., & Malayala, S. (2021). COVID-19 Encephalopathy in Adults. *Cureus*, *13*(2), e13052. https://doi.org/10.7759/cureus.13052
Auten, J. D., Kuhne, M. A., Walker, H. M., 2nd, & Porter, H. O. (2010). Neurologic decompression sickness following cabin pressure fluctuations at high altitude. *Aviat Space Environ Med*, *81*(4), 427-430. https://doi.org/10.3357/asem.2406.2010
Badran, M., Puech, C., Barrow, M. B., Runion, A. R., & Gozal, D. (2023). Solriamfetol enhances wakefulness and improves cognition and anxiety in a murine model of OSA. *Sleep Med*, *107*, 89-99. https://doi.org/10.1016/j.sleep.2023.04.007
Baggett, M. R., Kelly, M. P., Korenman, L. M., & Ryan, L. M. (2003). Neuropsychological deficits of a U.S. Army pilot following an anoxic event as a function of cardiac arrest. *Mil Med*, *168*(9), 769-771.
Bahemuka, M. (1983). Involuntary movements after closed head injury: hypoxic hypothesis. *East Afr Med J*, *60*(5), 340-342.
Bahreini, M., Talebi Garekani, M., Sotoodehnia, M., & Rasooli, F. (2021). Comparison of the efficacy of ketamine- propofol versus sodium thiopental-fentanyl in sedation: a randomised clinical trial. *Emerg Med J*, *38*(3), 211-216. https://doi.org/10.1136/emermed-2020-209542
Baker, T. L., Boyce, J., Gairy, P., & Mighty, G. (2011). Interprofessional management of a complex continuing care patient admitted with 18 pressure ulcers: a case report. *Ostomy Wound Manage*, *57*(2), 38-47.

Ballot, D. E., Rakotsoane, D., Cooper, P. A., Ramdin, T. D., Chirwa, T., & Pepper, M. S. (2020). A prospective observational study of developmental outcomes in survivors of neonatal hypoxic ischaemic encephalopathy in South Africa. *S Afr Med J*, *110*(4), 308-312. https://doi.org/10.7196/SAMJ.2020.v110i4.14311

Ballout, A. A., Kolesnik, M., Choi, Y., Ayoub, M. S., Harel, A., & Najjar, S. (2022). Case report: Bilateral globus pallidus lesions and delayed progressive leukoencephalopathy in COVID-19: Effects of hypoxia alone or combination of hypoxia and inflammation? *Front Neurol*, *13*, 1084831. https://doi.org/10.3389/fneur.2022.1084831

Banaei, P., Tadibi, V., Amiri, E., & Machado, D. (2023). Concomitant dual-site tDCS and dark chocolate improve cognitive and endurance performance following cognitive effort under hypoxia: a randomized controlled trial. *Sci Rep*, *13*(1), 16473. https://doi.org/10.1038/s41598-023-43568-y

Banderet, L. E., & Lieberman, H. R. (1989). Treatment with tyrosine, a neurotransmitter precursor, reduces environmental stress in humans. *Brain Res Bull*, *22*(4), 759-762. https://doi.org/10.1016/0361-9230(89)90096-8

Banta, G. R., & Kosnosky, D. P. (1978). Case report of an obsessive-compulsive personality: a precurser to accident proneness. *Aviat Space Environ Med*, *49*(6), 827-828.

Baron, J., & Auckley, D. (2004). Gunshot wound to the head: an unusual complication of sleep apnea and bilevel positive airway pressure. *Sleep Breath*, *8*(3), 161-164. https://doi.org/10.1007/s11325-004-0161-9

Bauer, J., Grunwald, T., Huppertz, H. J., König, K., Kohnen, O., Shala, J., & Jokeit, H. (2020). Social cognition in an adult epilepsy patient with developmental amnesia. *Neurocase*, *26*(4), 231-240. https://doi.org/10.1080/13554794.2020.1791904

Baumgartner, R. W., Keller, S., Regard, M., & Bärtsch, P. (2003). Flunarizine in prevention of headache, ataxia, and memory deficits during decompression to 4559 m. *High Alt Med Biol*, *4*(3), 333-339. https://doi.org/10.1089/152702903769192287

Bayer, U., Likar, R., Pinter, G., Stettner, H., Demschar, S., Trummer, B., Neuwersch, S., Glazachev, O., & Burtscher, M. (2019). Effects of intermittent hypoxia-hyperoxia on mobility and perceived health in geriatric patients performing a multimodal training intervention: a randomized controlled trial. *BMC Geriatr*, *19*(1), 167. https://doi.org/10.1186/s12877-019-1184-1

Beatty, W. W., Salmon, D. P., Bernstein, N., & Butters, N. (1987). Remote memory in a patient with amnesia due to hypoxia. *Psychol Med*, *17*(3), 657-665. https://doi.org/10.1017/s0033291700025897

Beatty, W. W., Salmon, D. P., Bernstein, N., Martone, M., Lyon, L., & Butters, N. (1987). Procedural learning in a patient with amnesia due to hypoxia. *Brain Cogn*, *6*(4), 386-402. https://doi.org/10.1016/0278-2626(87)90135-7

Beaumont, M., Batéjat, D., Coste, O., Van Beers, P., Colas, A., Clère, J. M., & Piérard, C. (2004). Effects of zolpidem and zaleplon on sleep, respiratory patterns and performance at a simulated altitude of 4,000 m. *Neuropsychobiology*, *49*(3), 154-162. https://doi.org/10.1159/000076723

Beaumont, M., Batéjat, D., Piérard, C., Van Beers, P., Philippe, M., Léger, D., Savourey, G., & Jouanin, J. C. (2007). Zaleplon and zolpidem objectively alleviate sleep disturbances in mountaineers at a 3,613 meter altitude. *Sleep*, *30*(11), 1527-1533. https://doi.org/10.1093/sleep/30.11.1527

Bekinschtein, T. A., Golombek, D. A., Simonetta, S. H., Coleman, M. R., & Manes, F. F. (2009). Circadian rhythms in the vegetative state. *Brain Inj*, *23*(11), 915-919. https://doi.org/10.1080/02699050903283197

Bekker, A., Shah, R., Quartermain, D., Li, Y. S., & Blanck, T. (2006). Isoflurane preserves spatial working memory in adult mice after moderate hypoxia. *Anesth Analg*, *102*(4), 1134-1138. https://doi.org/10.1213/01.ane.0000198637.36539.c1

Ben Mohamed, D., Zouari, R., Ketata, J., Nabli, F., Blel, S., & Ben Sassi, S. (2023). Myoclonus status revealing COVID 19 infection. *Seizure*, *104*, 12-14. https://doi.org/10.1016/j.seizure.2022.11.010

Benetoli, A., Paganelli, R. A., Giordani, F., Lima, K. C., Fávero Filho, L. A., & Milani, H. (2004). Effect of tacrolimus (FK506) on ischemia-induced brain damage and memory dysfunction in rats. *Pharmacol Biochem Behav*, *77*(3), 607-615. https://doi.org/10.1016/j.pbb.2003.12.022

Berman, D. J., Knibbs, N., Friedman, L., & Rocco, M. (2018). Postpartum hemoptysis as presenting sign of longstanding vasculitis. *Int J Obstet Anesth*, *36*, 122-125. https://doi.org/10.1016/j.ijoa.2018.07.002

Bhaiyat, A. M., Sasson, E., Wang, Z., Khairy, S., Ginzarly, M., Qureshi, U., Fikree, M., & Efrati, S. (2022). Hyperbaric oxygen treatment for long coronavirus disease-19: a case report. *J Med Case Rep*, *16*(1), 80. https://doi.org/10.1186/s13256-022-03287-w

Binns-Loveman, K. M., Kaplowitz, M. R., & Fike, C. D. (2005). Sildenafil and an early stage of chronic hypoxia-induced pulmonary hypertension in newborn piglets. *Pediatr Pulmonol*, *40*(1), 72-80. https://doi.org/10.1002/ppul.20229

Bird, C. M., Shallice, T., & Cipolotti, L. (2007). Fractionation of memory in medial temporal lobe amnesia. *Neuropsychologia*, *45*(6), 1160-1171. https://doi.org/10.1016/j.neuropsychologia.2006.10.011

Blackwell, S. C., Hallak, M., Hotra, J. W., Refuerzo, J., Sokol, R. J., & Sorokin, Y. (2004). Prolonged in utero meconium exposure impairs spatial learning in the adult rat. Central Prize Award. *Am J Obstet Gynecol*, *190*(6), 1551-1555; discussion 1555-1556. https://doi.org/10.1016/j.ajog.2004.03.048

Bojar, R. M., Rastegar, H., Payne, D. D., Harkness, S. H., England, M. R., Stetz, J. J., Weiner, B., & Cleveland, R. J. (1987). Methemoglobinemia from intravenous nitroglycerin: a word of caution. *Ann Thorac Surg*, *43*(3), 332-334. https://doi.org/10.1016/s0003-4975(10)60627-3

Bolouri, M. R., & Small, G. A. (2004). Neuroimaging of hypoxia and cocaine-induced hippocampal stroke. *J Neuroimaging*, *14*(3), 290-291. https://doi.org/10.1177/1051228404265751

Bonifacio, S. L., McDonald, S. A., Chock, V. Y., Wusthoff, C. J., Hintz, S. R., Laptook, A. R., Shankara, S., & Van Meurs, K. P. (2019). Differences in patient characteristics and care practices between two trials of therapeutic hypothermia. *Pediatr Res*, *85*(7), 1008-1015. https://doi.org/10.1038/s41390-019-0371-2

Bornschein, S., Hausteiner, C., Römmelt, H., Nowak, D., Förstl, H., & Zilker, T. (2008). Double-blind placebo-controlled provocation study in patients with subjective Multiple Chemical Sensitivity (MCS) and matched control subjects. *Clin Toxicol (Phila)*, *46*(5), 443-449. https://doi.org/10.1080/15563650701742438

Bosco, G., Ionadi, A., Data, P. G., & Mortola, J. P. (2004). Voluntary breath-holding in the morning and in the evening. *Clin Sci (Lond)*, *106*(4), 347-352. https://doi.org/10.1042/cs20030260

Bosco, M. C., Puppo, M., Santangelo, C., Anfosso, L., Pfeffer, U., Fardin, P., Battaglia, F., & Varesio, L. (2006). Hypoxia modifies the transcriptome of primary human monocytes: modulation of novel immune-related genes and identification of CC-chemokine ligand 20 as a new hypoxia-inducible gene. *J Immunol*, *177*(3), 1941-1955. https://doi.org/10.4049/jimmunol.177.3.1941

Boutelier, A., Ollivier, V., Mazighi, M., Kyheng, M., Labreuche, J., Brikci-Nigassa, N., Solo Nomenjanahary, M., Delvoye, F., Maier, B., Paquet, C., Ho-Tin-Noe, B., & Desilles, J. P. (2024). Acute astrocytic reaction is associated with 3-month functional outcome after stroke treated with endovascular therapy. *Eur Stroke J*, *9*(4), 952-958. https://doi.org/10.1177/23969873241256813

Bouzat, P., Séchaud, G., Banco, P., Davranche, K., Casini, L., Baillieul, S., Manhes, P., Botrè, F., Mazzarino, M., De la Torre, X., Robach, P., & Verges, S. (2018). The effect of zolpidem on cognitive function and postural control at high altitude. *Sleep*, *41*(10). https://doi.org/10.1093/sleep/zsy153

Bowden, V. K., & Loft, S. (2016). Using memory for prior aircraft events to detect conflicts under conditions of proactive air traffic control and with concurrent task requirements. *J Exp Psychol Appl*, *22*(2), 211-224. https://doi.org/10.1037/xap0000085

Bradke, B. S., & Everman, B. R. (2020). Mild Hypoxia of a Skydiver Making Repeated, Medium-Altitude Aircraft Exits. *Aerosp Med Hum Perform*, *91*(2), 110-115. https://doi.org/10.3357/amhp.5497.2020

Bradley, K. T., Jamie, L. K. J. L., Richards, M. F., Beckstrand, D. P., & Wolf, E. G. (2020). Chronic decompression illness cognitive dysfunction improved with hyperbaric oxygen: a case report. *Undersea Hyperb Med*, *47*(1), 131-137. https://doi.org/10.22462/01.03.2020.14

Braga, V. A., Burmeister, M. A., Sharma, R. V., & Davisson, R. L. (2008). Cardiovascular responses to peripheral chemoreflex activation and comparison of different methods to evaluate baroreflex gain in conscious mice using telemetry. *Am J Physiol Regul Integr Comp Physiol*, *295*(4), R1168-1174. https://doi.org/10.1152/ajpregu.90375.2008

Brefel-Courbon, C., Payoux, P., Ory, F., Sommet, A., Slaoui, T., Raboyeau, G., Lemesle, B., Puel, M., Montastruc, J. L., Demonet, J. F., & Cardebat, D. (2007). Clinical and imaging evidence of zolpidem effect in hypoxic encephalopathy. *Ann Neurol*, *62*(1), 102-105. https://doi.org/10.1002/ana.21110

Brenn, B. R., Brislin, R. P., & Rose, J. B. (1998). Epidural analgesia in children with cerebral palsy. *Can J Anaesth*, *45*(12), 1156-1161. https://doi.org/10.1007/bf03012456

Breuking, S. H., Jansen, C., de Haan, T. R., & Bakker, P. (2024). How cold is too cold during maternal sepsis? Navigating between maternal hypothermia and fetal bradycardia. *Eur J Obstet Gynecol Reprod Biol*, *302*, 394-396. https://doi.org/10.1016/j.ejogrb.2024.09.015

Briggs, E. R. (2009). Taste disturbances related to medication use. *Consult Pharm*, *24*(7), 538-543. https://doi.org/10.4140/tcp.n.2009.538

Briscoe, J., Gathercole, S. E., & Marlow, N. (2001). Everyday memory and cognitive ability in children born very prematurely. *J Child Psychol Psychiatry*, *42*(6), 749-754. https://doi.org/10.1111/1469-7610.00771

Brockmann, K., Stolpe, S., Fels, C., Khan, N., Kulozik, A. E., & Pekrun, A. (2005). Moyamoya syndrome associated with hemolytic anemia due to Hb Alesha. *J Pediatr Hematol Oncol*, *27*(8), 436-440. https://doi.org/10.1097/01.mph.0000175409.21342.ea

Broman, M., Rose, A. L., Hotson, G., & Casey, C. M. (1997). Severe anterograde amnesia with onset in childhood as a result of anoxic encephalopathy. *Brain*, *120 ( Pt 3)*, 417-433. https://doi.org/10.1093/brain/120.3.417

Brown, T. M. (2013). A case of Shoshin Beriberi: lessons old and new for the psychiatrist. *Psychosomatics*, *54*(2), 175-180. https://doi.org/10.1016/j.psym.2012.01.010

Bucuk, M., Tomic, Z., Tuskan-Mohar, L., Bonifacic, D., Bralic, M., & Jurjevic, A. (2008). Recurrent transient global amnesia at high altitude. *High Alt Med Biol*, *9*(3), 239-240. https://doi.org/10.1089/ham.2008.0002

Bush, P. G., Mayhew, T. M., Abramovich, D. R., Aggett, P. J., Burke, M. D., & Page, K. R. (2000). A quantitative study on the effects of maternal smoking on placental morphology and cadmium concentration. *Placenta*, *21*(2-3), 247-256. https://doi.org/10.1053/plac.1999.0470

Busija, D. W. (1984). Sympathetic nerves reduce cerebral blood flow during hypoxia in awake rabbits. *Am J Physiol*, *247*(3 Pt 2), H446-451. https://doi.org/10.1152/ajpheart.1984.247.3.H446

Calder, K., Kokorowski, P., Tran, T., & Henderson, S. (2003). Emergency department

presentation of pediatric stroke. *Pediatr Emerg Care*, *19*(5), 320-328. https://doi.org/10.1097/01.pec.0000092577.40174.61
Cannon, T. D., van Erp, T. G., & Glahn, D. C. (2002). Elucidating continuities and discontinuities between schizotypy and schizophrenia in the nervous system. *Schizophr Res*, *54*(1-2), 151-156. https://doi.org/10.1016/s0920-9964(01)00362-0
Caputa, M., Rogalska, J., Wentowska, K., & Nowakowska, A. (2005). Perinatal asphyxia, hyperthermia and hyperferremia as factors inducing behavioural disturbances in adulthood: a rat model. *Behav Brain Res*, *163*(2), 246-256. https://doi.org/10.1016/j.bbr.2005.05.015
Car, H., Wiśniewska, R. J., & Wiśniewski, K. (2004). 2R,4R-APDC influence on hypoxia-induced impairment of learning and memory processes in passive avoidance test. *Pol J Pharmacol*, *56*(5), 527-537.
Carbonnel, S., Charnallet, A., David, D., & Pellat, J. (1997). One or several semantic system(s)? Maybe none: evidence from a case study of modality and category-specific "semantic" impairment. *Cortex*, *33*(3), 391-417. https://doi.org/10.1016/s0010-9452(08)70227-2
Cardoso Vale, T., Echenique, L., Barsottini, O. G. P., & Pedroso, J. L. (2020). Paroxysmal Autonomic Instability with Dystonia after Severe Traumatic Brain Injury. *Tremor Other Hyperkinet Mov (N Y)*, *10*, 12. https://doi.org/10.5334/tohm.81
Carver, R. P., & Winsmann, F. R. (1968). Effect of high elevation upon physical proficiency, cognitive functioning and subjective symptomatology. *Percept Mot Skills*, *26*(1), 223-230. https://doi.org/10.2466/pms.1968.26.1.223
Casati, A., Fanelli, G., Pietropaoli, P., Proietti, R., Tufano, R., Danelli, G., Fierro, G., De Cosmo, G., & Servillo, G. (2005). Continuous monitoring of cerebral oxygen saturation in elderly patients undergoing major abdominal surgery minimizes brain exposure to potential hypoxia. *Anesth Analg*, *101*(3), 740-747. https://doi.org/10.1213/01.ane.0000166974.96219.cd
Cecere, R., Romei, V., Bertini, C., & Làdavas, E. (2014). Crossmodal enhancement of visual orientation discrimination by looming sounds requires functional activation of primary visual areas: a case study. *Neuropsychologia*, *56*, 350-358. https://doi.org/10.1016/j.neuropsychologia.2014.02.008
Chachkhiani, D., Chimakurthy, A. K., Verdecie, O., Goyne, C. T., & Mader, E. C., Jr. (2021). Delayed Toxic-Hypoxic Leukoencephalopathy As Sequela of Opioid Overdose and Cerebral Hypoxia-Ischemia. *Cureus*, *13*(12), e20271. https://doi.org/10.7759/cureus.20271
Chail, A., Baby, S., Sharma, R., & Dubey, A. (2019). Impact of eclectic cognitive retraining in a case of high-altitude cerebral edema. *Ind Psychiatry J*, *28*(2), 318-320. https://doi.org/10.4103/ipj.ipj_108_20
Chalak, L. F., Nguyen, K. A., Prempunpong, C., Heyne, R., Thayyil, S., Shankaran, S., Laptook, A. R., Rollins, N., Pappas, A., Koclas, L., Shah, B., Montaldo, P., Techasaensiri, B., Sánchez, P. J., & Sant'Anna, G. (2018). Prospective research in infants with mild encephalopathy identified in the first six hours of life: neurodevelopmental outcomes at 18-22 months. *Pediatr Res*, *84*(6), 861-868. https://doi.org/10.1038/s41390-018-0174-x
Chan, C. C., Wu, H. C., Wu, C. H., & Hsu, C. Y. (1995). Hepatopulmonary syndrome in liver cirrhosis: report of a case. *J Formos Med Assoc*, *94*(4), 185-188.
Chanana, V., Hackett, M., Deveci, N., Aycan, N., Ozaydin, B., Cagatay, N. S., Hanalioglu, D., Kintner, D. B., Corcoran, K., Yapici, S., Camci, F., Eickhoff, J., Frick, K. M., Ferrazano, P., Levine, J. E., & Cengiz, P. (2023). TrkB-mediated sustained neuroprotection is sex-specific and ERα dependent in adult mice following neonatal hypoxia ischemia. *Res Sq*. https://doi.org/10.21203/rs.3.rs-3325405/v1
Chang, C. H., Lian, H. W., & Sung, Y. F. (2022). Cystic Encephalomalacia in a Young Woman After Cardiac Arrest Due to Diabetic Ketoacidosis and Thyroid Storm. *Cureus*, *14*(3),

e23707. https://doi.org/10.7759/cureus.23707
Chapman, K. R., & Rebuck, A. S. (1991). Dysphagia as a manifestation of occult hypoxemia. The role of oximetry during meal times. *Chest*, *99*(4), 1030-1032. https://doi.org/10.1378/chest.99.4.1030
Charnallet, A., Carbonnel, S., David, D., & Moreaud, O. (2008). Associative visual agnosia: a case study. *Behav Neurol*, *19*(1-2), 41-44. https://doi.org/10.1155/2008/241753
Chen, D. Y., Di, X., Amaya, N., Sun, H., Pal, S., & Biswal, B. B. (2024). Brain activation during the N-back working memory task in individuals with spinal cord injury: a functional near-infrared spectroscopy study. *bioRxiv*. https://doi.org/10.1101/2024.02.09.579655
Chen, L., Bai, S., Su, W., Song, X., Zhang, P., Li, L., & Ji, J. J. (2011). Transient oxygen-glucose deprivation causes immediate changes in redox activity in mouse brain tissue. *Brain Res*, *1390*, 99-107. https://doi.org/10.1016/j.brainres.2011.03.022
Chen, L., Wang, L., Zhuo, Q., Zhang, Q., Chen, F., Li, L., & Lin, L. (2018). Effect of Shenmai injection on cognitive function after cardiopulmonary bypass in cardiac surgical patients: a randomized controlled trial. *BMC Anesthesiol*, *18*(1), 142. https://doi.org/10.1186/s12871-018-0604-7
Chen, W. F., Hsu, J. H., Lin, C. S., Jong, Y. J., Yang, C. H., Huang, L. T., & Yang, S. N. (2011). Granulocyte-colony stimulating factor alleviates perinatal hypoxia-induced decreases in hippocampal synaptic efficacy and neurogenesis in the neonatal rat brain. *Pediatr Res*, *70*(6), 589-595. https://doi.org/10.1203/PDR.0b013e3182324424
Cheng, M. I., Hong, L., Chen, B., Chin, S., Luthers, C. R., Bustillos, C., Sheikh, S. Z., & Su, M. A. (2023). Hypoxia-sensing by the Histone Demethylase UTX ( KDM6A ) Controls Colitogenic CD4 (+) T cell Fate and Mucosal Inflammation. *bioRxiv*. https://doi.org/10.1101/2023.07.27.550746
Chevret, S., Verlhac, S., Ducros-Miralles, E., Dalle, J. H., de Latour, R. P., de Montalembert, M., Benkerrou, M., Pondarré, C., Thuret, I., Guitton, C., Lesprit, E., Etienne-Julan, M., Elana, G., Vannier, J. P., Lutz, P., Neven, B., Galambrun, C., Paillard, C., Runel, C., . . . Bernaudin, F. (2017). Design of the DREPAGREFFE trial: A prospective controlled multicenter study evaluating the benefit of genoidentical hematopoietic stem cell transplantation over chronic transfusion in sickle cell anemia children detected to be at risk of stroke by transcranial Doppler (NCT 01340404). *Contemp Clin Trials*, *62*, 91-104. https://doi.org/10.1016/j.cct.2017.08.008
Chia, K. X., Polakhare, S., & Bruno, S. D. (2020). Possible affective cognitive cerebellar syndrome in a young patient with COVID-19 CNS vasculopathy and stroke. *BMJ Case Rep*, *13*(10). https://doi.org/10.1136/bcr-2020-237926
Chiu, T. F., Yang, C. C., Ger, J., Deng, J. F., & Bullard, M. J. (1994). Adult respiratory distress syndrome and late death following imipramine overdose: a case report. *Zhonghua Yi Xue Za Zhi (Taipei)*, *54*(6), 436-441.
Churchill, S., Weaver, L. K., Deru, K., Russo, A. A., Handrahan, D., Orrison, W. W., Jr., Foley, J. F., & Elwell, H. A. (2013). A prospective trial of hyperbaric oxygen for chronic sequelae after brain injury (HYBOBI). *Undersea Hyperb Med*, *40*(2), 165-193.
Cioffi, A., Cecannecchia, C., Bosco, M. A., Gurgoglione, G., Baldari, B., & De Simone, S. (2023). Lethal nitrous oxide (N(2)O) intoxication during surgery: the contribution of immunohistochemistry in identifying the cause of death: a case report. *J Med Case Rep*, *17*(1), 424. https://doi.org/10.1186/s13256-023-04159-7
Clincke, G. H., & Wauquier, A. (1984). Pharmacological protection against hypoxia-induced effects on medium-term memory in a two-way avoidance paradigm. *Behav Brain Res*, *14*(2), 139-142. https://doi.org/10.1016/0166-4328(84)90181-5
Cohen, R. D., Galko, B. M., Contreras, M., Kenny, F. T., & Rebuck, A. S. (1986). Neuropsychological effects of short-term discontinuation of oxygen therapy. Observations in patients with chronic hypoxemia who are receiving long-term oxygen

therapy. *Arch Intern Med*, *146*(8), 1557-1559.
Cohen-Zion, M., Stepnowsky, C., Johnson, S., Marler, M., Dimsdale, J. E., & Ancoli-Israel, S. (2004). Cognitive changes and sleep disordered breathing in elderly: differences in race. *J Psychosom Res*, *56*(5), 549-553. https://doi.org/10.1016/j.jpsychores.2004.02.002
Coleman, K., Phillips, J., Sciarini, M., Stubbs, B., Jackson, O., & Kernagis, D. (2021). A Metabolic Intervention for Improving Human Cognitive Performance During Hypoxia. *Aerosp Med Hum Perform*, *92*(7), 556-562. https://doi.org/10.3357/amhp.5767.2021
Coppa, G. F., Gouge, T. H., & Hofstetter, S. R. (1981). Air embolism: a lethal but preventable complication of subclavian vein catheterization. *JPEN J Parenter Enteral Nutr*, *5*(2), 166-168. https://doi.org/10.1177/0148607181005002166
Corn, M., Pham, T., & Kemp, W. (2023). Adverse Fetal Outcomes and Histopathology of Placentas Affected by COVID-19: A Report of Four Cases. *Cureus*, *15*(8), e44402. https://doi.org/10.7759/cureus.44402
Costeff, H., Cohen, B. E., Weller, L., & Kleckner, H. (1981). Pathogenic factors in idiopathic mental retardation. *Dev Med Child Neurol*, *23*(4), 484-493. https://doi.org/10.1111/j.1469-8749.1981.tb02022.x
Cotroneo, A. M., Castagna, A., Putignano, S., Lacava, R., Fantò, F., Monteleone, F., Rocca, F., Malara, A., & Gareri, P. (2013). Effectiveness and safety of citicoline in mild vascular cognitive impairment: the IDEALE study. *Clin Interv Aging*, *8*, 131-137. https://doi.org/10.2147/cia.S38420
Cotten, C. M., Murtha, A. P., Goldberg, R. N., Grotegut, C. A., Smith, P. B., Goldstein, R. F., Fisher, K. A., Gustafson, K. E., Waters-Pick, B., Swamy, G. K., Rattray, B., Tan, S., & Kurtzberg, J. (2014). Feasibility of autologous cord blood cells for infants with hypoxic-ischemic encephalopathy. *J Pediatr*, *164*(5), 973-979.e971. https://doi.org/10.1016/j.jpeds.2013.11.036
Cruikshank, B. M., Eliason, M., & Merrifield, B. (1988). Long-term sequelae of cold water near-drowning. *J Pediatr Psychol*, *13*(3), 379-388. https://doi.org/10.1093/jpepsy/13.3.379
Culley, D. J., Baxter, M., Yukhananov, R., & Crosby, G. (2003). The memory effects of general anesthesia persist for weeks in young and aged rats. *Anesth Analg*, *96*(4), 1004-1009. https://doi.org/10.1213/01.Ane.0000052712.67573.12
Cunningham, D. R., Cunningham, C. A., & Vise, L. K. (1985). The effects of chronic hypoxemia on central auditory processing in patients with chronic obstructive pulmonary disease. *Ear Hear*, *6*(6), 297-303. https://doi.org/10.1097/00003446-198511000-00004
Custodio, C. M., & Basford, J. R. (2004). Delayed postanoxic encephalopathy: a case report and literature review. *Arch Phys Med Rehabil*, *85*(3), 502-505. https://doi.org/10.1016/s0003-9993(03)00471-4
Daher, A., Balfanz, P., Aetou, M., Hartmann, B., Müller-Wieland, D., Müller, T., Marx, N., Dreher, M., & Cornelissen, C. G. (2021). Clinical course of COVID-19 patients needing supplemental oxygen outside the intensive care unit. *Sci Rep*, *11*(1), 2256. https://doi.org/10.1038/s41598-021-81444-9
Dalecki, M., Bock, O., & Guardiera, S. (2010). Simulated flight path control of fighter pilots and novice subjects at +3 Gz in a human centrifuge. *Aviat Space Environ Med*, *81*(5), 484-488. https://doi.org/10.3357/asem.2665.2010
Daly, M. W., Custer, G., & McLeay, P. D. (2008). Cardiac arrest with pulseless electrical activity associated with methylphenidate in an adolescent with a normal baseline echocardiogram. *Pharmacotherapy*, *28*(11), 1408-1412. https://doi.org/10.1592/phco.28.11.1408
Damasceno, B. P. (1991). Decerebrate rigidity with preserved cognition and gait: a possible role of anoxic-ischemic brain damage. *Int J Neurosci*, *58*(3-4), 283-287. https://doi.org/10.3109/00207459108985444

Dandrea, K. E., & Cotten, J. F. (2021). A Comparison of Breathing Stimulants for Reversal of Synthetic Opioid-Induced Respiratory Depression in Conscious Rats. *J Pharmacol Exp Ther*, *378*(2), 146-156. https://doi.org/10.1124/jpet.121.000675

Dang, S., Yan, H., Yamamoto, S., Wang, X., & Zeng, L. (2004). Poor nutritional status of younger Tibetan children living at high altitudes. *Eur J Clin Nutr*, *58*(6), 938-946. https://doi.org/10.1038/sj.ejcn.1601915

Dattilo, G., Tulino, V., Tulino, D., Lamari, A., Falanga, G., Marte, F., & Patanè, S. (2011). Perinatal asphyxia and cardiac abnormalities. *Int J Cardiol*, *147*(2), e39-40. https://doi.org/10.1016/j.ijcard.2009.01.032

de Aquino, M. M., & Cecatti, J. G. (2003). Misoprostol versus oxytocin for labor induction in term and post-term pregnancy: randomized controlled trial. *Sao Paulo Med J*, *121*(3), 102-106. https://doi.org/10.1590/s1516-31802003000300003

De Koninck, B. P., Brazeau, D., Deshaies, A. A., Briand, M. M., Maschke, C., Williams, V., Arbour, C., Williamson, D., Duclos, C., Bernard, F., Blain-Moraes, S., & De Beaumont, L. (2024). Modulation of brain activity in brain-injured patients with a disorder of consciousness in intensive care with repeated 10-Hz transcranial alternating current stimulation (tACS): a randomised controlled trial protocol. *BMJ Open*, *14*(7), e078281. https://doi.org/10.1136/bmjopen-2023-078281

De Luca, F., Benuzzi, F., Bertossi, E., Braghittoni, D., di Pellegrino, G., & Ciaramelli, E. (2018). Episodic future thinking and future-based decision-making in a case of retrograde amnesia. *Neuropsychologia*, *110*, 92-103. https://doi.org/10.1016/j.neuropsychologia.2017.08.007

De Renzi, E., & Lucchelli, F. (1993). Dense retrograde amnesia, intact learning capability and abnormal forgetting rate: a consolidation deficit? *Cortex*, *29*(3), 449-466. https://doi.org/10.1016/s0010-9452(13)80253-5

de Wilton, A., Kilich, E., Chaudhry, Z., Bell, L. C., Gahir, J., Cadman, J., Lever, R. A., & Logan, S. A. (2020). Delayed healthcare seeking and prolonged illness in healthcare workers during the COVID-19 pandemic: a single-centre observational study. *BMJ Open*, *10*(11), e040216. https://doi.org/10.1136/bmjopen-2020-040216

Debette, S., Kozlowski, O., Steinling, M., & Rousseaux, M. (2002). Levodopa and bromocriptine in hypoxic brain injury. *J Neurol*, *249*(12), 1678-1682. https://doi.org/10.1007/s00415-002-0903-1

Decell, M. K., Gordon, J. B., Silver, K., & Meagher-Villemure, K. (1994). Fulminant hepatic failure associated with status epilepticus in children: three cases and a review of potential mechanisms. *Intensive Care Med*, *20*(5), 375-378. https://doi.org/10.1007/bf01720913

Decker, M. J., Hue, G. E., Caudle, W. M., Miller, G. W., Keating, G. L., & Rye, D. B. (2003). Episodic neonatal hypoxia evokes executive dysfunction and regionally specific alterations in markers of dopamine signaling. *Neuroscience*, *117*(2), 417-425. https://doi.org/10.1016/s0306-4522(02)00805-9

DeKosky, S. T., Kochanek, P. M., Valadka, A. B., Clark, R. S. B., Chou, S. H., Au, A. K., Horvat, C., Jha, R. M., Mannix, R., Wisniewski, S. R., Wintermark, M., Rowell, S. E., Welch, R. D., Lewis, L., House, S., Tanzi, R. E., Smith, D. R., Vittor, A. Y., Denslow, N. D., . . . Hayes, R. L. (2021). Blood Biomarkers for Detection of Brain Injury in COVID-19 Patients. *J Neurotrauma*, *38*(1), 1-43. https://doi.org/10.1089/neu.2020.7332

Della Sala, S., & Spinnler, H. (1986). 'Indifférence amnésique' in a case of global amnesia following acute brain hypoxia. *Eur Neurol*, *25*(2), 98-109. https://doi.org/10.1159/000115994

Demay, F., & Bande, J. (1980). The effect of piracetam on volunteers in a low-pressure tank. *J Int Med Res*, *8*(1), 90-94. https://doi.org/10.1177/030006058000800116

Demeyere, N., Lestou, V., & Humphreys, G. W. (2010). Neuropsychological evidence for a

dissociation in counting and subitizing. *Neurocase*, *16*(3), 219-237. https://doi.org/10.1080/13554790903405719
DeNoble, V. J., DeNoble, K. F., Spencer, K. R., Johnson, L. C., Cook, L., Myers, M. J., & Scribner, R. M. (1990). Comparison of DuP 996, with physostigmine, THA and 3,4-DAP on hypoxia-induced amnesia in rats. *Pharmacol Biochem Behav*, *36*(4), 957-961. https://doi.org/10.1016/0091-3057(90)90106-r
Deonarain, D., & Karki, P. (2022). Surviving a 400 m Fall on Mount Everest. *Wilderness Environ Med*, *33*(4), 460-463. https://doi.org/10.1016/j.wem.2022.06.005
Devereaux, M. W., & Partnow, M. J. (1975). Delayed hypoxic encephalopathy without cognitive dysfunction. *Arch Neurol*, *32*(10), 704-705. https://doi.org/10.1001/archneur.1975.00490520074013
Dhok, S. M., Gudipati, A. R., Kaul, S., & Papalkar, A. S. (2023). Neuroimaging Features of High-Altitude Cerebral Edema: A Case Report. *Neurol India*, *71*(6), 1254-1256. https://doi.org/10.4103/0028-3886.391385
Dimitrova, D. S., & Getova-Spassova, D. P. (2006). Effects of galantamine and donepezil on active and passive avoidance tests in rats with induced hypoxia. *J Pharmacol Sci*, *101*(3), 199-204. https://doi.org/10.1254/jphs.fpe05006x
Dobashi, S., Koyama, K., Endo, J., Kiuchi, M., & Horiuchi, M. (2019). Impact of Dietary Nitrate Supplementation on Executive Function During Hypoxic Exercise. *High Alt Med Biol*, *20*(2), 187-191. https://doi.org/10.1089/ham.2018.0114
Dobrynina, L. A., Alexandrova, E. V., Zabitova, M. R., Kalashnikova, L. A., Krotenkova, M. V., & Akhmetzyanov, B. M. (2021). Anti-NR2 glutamate receptor antibodies as an early biomarker of cerebral small vessel disease. *Clin Biochem*, *96*, 26-32. https://doi.org/10.1016/j.clinbiochem.2021.07.003
Doneddu, A., Roberto, S., Guicciardi, M., Pazzona, R., Manca, A., Monni, A., Fanni, M., Leban, B., Ghiani, G., Spranger, M. D., Mulliri, G., & Crisafulli, A. (2024). Hemodynamics and cerebral oxygenation during acute exercise in moderate normobaric hypoxia and with concurrent cognitive task in young healthy males. *Appl Physiol Nutr Metab*, *49*(11), 1573-1584. https://doi.org/10.1139/apnm-2023-0629
Dopwell, F., Maypole, J., Sinha, B., Currier, H., DeBassio, W., & Augustyn, M. (2014). "More than meets the eye": when the neonatal course may impact several years out. *J Dev Behav Pediatr*, *35*(7), 467-469. https://doi.org/10.1097/dbp.0000000000000085
Downard, C. D., Grant, S. N., Matheson, P. J., Guillaume, A. W., Debski, R., Fallat, M. E., & Garrison, R. N. (2011). Altered intestinal microcirculation is the critical event in the development of necrotizing enterocolitis. *J Pediatr Surg*, *46*(6), 1023-1028. https://doi.org/10.1016/j.jpedsurg.2011.03.023
Drago, F., Grassi, M., Valerio, C., Coppi, G., Lauria, N., Nicotra, G. C., & Raffaele, R. (1991). Behavioral changes induced by the thyrotropin-releasing hormone analogue, RGH 2202. *Peptides*, *12*(6), 1309-1313. https://doi.org/10.1016/0196-9781(91)90212-8
Drummond, S. R., & Dutton, G. N. (2007). Simultanagnosia following perinatal hypoxia: a possible pediatric variant of Balint syndrome. *J aapos*, *11*(5), 497-498. https://doi.org/10.1016/j.jaapos.2007.03.007
Duan, J., Han, X., Bai, L., Zhou, L., & Huang, S. (2017). Assessment of heart rate, acidosis, consciousness, oxygenation, and respiratory rate to predict noninvasive ventilation failure in hypoxemic patients. *Intensive Care Med*, *43*(2), 192-199. https://doi.org/10.1007/s00134-016-4601-3
Dubiel, M., Gudmundsson, S., Thuring-Jönsson, A., Maesel, A., & Marsal, K. (1997). Doppler velocimetry and nonstress test for predicting outcome of pregnancies with decreased fetal movements. *Am J Perinatol*, *14*(3), 139-144. https://doi.org/10.1055/s-2007-994114
Dubsky, P., Sevelda, P., Jakesz, R., Hausmaninger, H., Samonigg, H., Seifert, M., Denison, U., Mlineritsch, B., Steger, G., Kwasny, W., Stöger, H., Bartsch, R., Stierer, M., Taucher, S.,

Fridrik, M., Schippinger, W., Greil, R., Pötter, R., & Gnant, M. (2008). Anemia is a significant prognostic factor in local relapse-free survival of premenopausal primary breast cancer patients receiving adjuvant cyclophosphamide/methotrexate/5-fluorouracil chemotherapy. *Clin Cancer Res*, *14*(7), 2082-2087. https://doi.org/10.1158/1078-0432.Ccr-07-2068
Duderstadt, Y., Schreiber, S., Burtscher, J., Schega, L., Müller, N. G., Brigadski, T., Braun-Dullaeus, R. C., Leßmann, V., & Müller, P. (2024). Controlled Hypoxia Acutely Prevents Physical Inactivity-Induced Peripheral BDNF Decline. *Int J Mol Sci*, *25*(14). https://doi.org/10.3390/ijms25147536
Duff, M. C., Hengst, J., Tranel, D., & Cohen, N. J. (2006). Development of shared information in communication despite hippocampal amnesia. *Nat Neurosci*, *9*(1), 140-146. https://doi.org/10.1038/nn1601
Duraski, S. A. (2017). The Importance of Monitoring Patients' Responses to Medications: Increased Arousal after Administration of Zolpidem in Those with Hypoxic Ischemic Encephalopathy-A Case Study. *Rehabil Nurs*, *42*(2), 75-79. https://doi.org/10.1002/rnj.232
Dyson, E., Voisey, S., Hughes, S., Higgins, B., & McQuillan, P. J. (2004). Educational psychology in medical learning: a randomised controlled trial of two aide memoires for the recall of causes of electromechanical dissociation. *Emerg Med J*, *21*(4), 457-460. https://doi.org/10.1136/emj.2003.012377
Eames, P. (1992). Hysteria following brain injury. *J Neurol Neurosurg Psychiatry*, *55*(11), 1046-1053. https://doi.org/10.1136/jnnp.55.11.1046
Ebert, A. D., Walzer, T. A., Huth, C., & Herrmann, M. (2001). Early neurobehavioral disorders after cardiac surgery: a comparative analysis of coronary artery bypass graft surgery and valve replacement. *J Cardiothorac Vasc Anesth*, *15*(1), 15-19. https://doi.org/10.1053/jcan.2001.20211
Edby, K., Larsson, J., Eek, M., von Wendt, L., & Ostergård, B. (1995). Amantadine treatment of a patient with anoxic brain injury. *Childs Nerv Syst*, *11*(10), 607-609. https://doi.org/10.1007/bf00301001
Edelstyn, N. M., Baker, S. R., Ellis, S. J., & Jenkinson, P. (2004). A cognitive neuropsychological and psychophysiological investigation of a patient who exhibited an acute exacerbated behavioural response during innocuous somatosensory stimulation and movement. *Behav Neurol*, *15*(1-2), 15-22. https://doi.org/10.1155/2004/458327
Eggers, S. D., Moster, M. L., & Cranmer, K. (2008). Selective saccadic palsy after cardiac surgery. *Neurology*, *70*(4), 318-320. https://doi.org/10.1212/01.wnl.0000287139.01789.97
Eguchi, H., Hotta, F., Kuwahara, T., Nakayama-Imaohji, H., Kusaka, S., & Shimomura, Y. (2017). Acute keratoconjunctivitis due to contamination of contact lens care solution with histamine-producing Raoultella species: A case report. *Medicine (Baltimore)*, *96*(50), e9310. https://doi.org/10.1097/md.0000000000009310
Eicher, D. J., Wagner, C. L., Katikaneni, L. P., Hulsey, T. C., Bass, W. T., Kaufman, D. A., Horgan, M. J., Languani, S., Bhatia, J. J., Givelichian, L. M., Sankaran, K., & Yager, J. Y. (2005). Moderate hypothermia in neonatal encephalopathy: efficacy outcomes. *Pediatr Neurol*, *32*(1), 11-17. https://doi.org/10.1016/j.pediatrneurol.2004.06.014
Elhakim, M., Abdelhamid, D., Abdelfattach, H., Magdy, H., Elsayed, A., & Elshafei, M. (2010). Effect of epidural dexmedetomidine on intraoperative awareness and post-operative pain after one-lung ventilation. *Acta Anaesthesiol Scand*, *54*(6), 703-709. https://doi.org/10.1111/j.1399-6576.2009.02199.x
Elzinga, L., Marcus, M., Peek, D., Borg, P., Jansen, J., Koster, J., & Enk, D. (2009). Hemodynamic stability ensured by a low dose, low volume, unilateral hypobaric spinal block: modification of a technique. *Acta Anaesthesiol Belg*, *60*(4), 217-220.

Emekli, A. S., Ekizoglu, E., & Yesilot, N. (2020). Diffuse enlargement of cerebral vasculature in an adult patient operated for cyanotic congenital heart disease. *Cardiol Young*, *30*(5), 734-736. https://doi.org/10.1017/s1047951120000669
Emmelkamp, P. M., & Felten, M. (1985). The process of exposure in vivo: cognitive and physiological changes during treatment of acrophobia. *Behav Res Ther*, *23*(2), 219-223. https://doi.org/10.1016/0005-7967(85)90034-8
Engelhardt, S., Huang, S. F., Patkar, S., Gassmann, M., & Ogunshola, O. O. (2015). Differential responses of blood-brain barrier associated cells to hypoxia and ischemia: a comparative study. *Fluids Barriers CNS*, *12*, 4. https://doi.org/10.1186/2045-8118-12-4
Epstein, C. M. (2003). Aliasing in the visual EEG: a potential pitfall of video display technology. *Clin Neurophysiol*, *114*(10), 1974-1976. https://doi.org/10.1016/s1388-2457(03)00168-8
Ercoline, W. R., Self, B. P., & Matthews, R. S. (2002). Effects of three helmet-mounted display symbologies on unusual attitude recognition and recovery. *Aviat Space Environ Med*, *73*(11), 1053-1058.
Ergun-Longmire, B., Nguyen, M. H. N., & Com, G. (2022). Electrical status epilepticus during sleep in a child with Prader-Willi syndrome: a case report. *AME Case Rep*, *6*, 7. https://doi.org/10.21037/acr-21-34
Escapita, A. C., Thomas, J. G., & Johnson, T. L. (2024). A case report of spastic diplegic cerebral palsy in a late preterm child with hypoplastic left heart syndrome. *Transl Pediatr*, *13*(7), 1258-1265. https://doi.org/10.21037/tp-24-57
Esfahani-Bayerl, N., Finke, C., Braun, M., Düzel, E., Heekeren, H. R., Holtkamp, M., Hasper, D., Storm, C., & Ploner, C. J. (2016). Visuo-spatial memory deficits following medial temporal lobe damage: A comparison of three patient groups. *Neuropsychologia*, *81*, 168-179. https://doi.org/10.1016/j.neuropsychologia.2015.12.024
Etemadi, M., Amiri, E., Tadibi, V., Grospretre, S., Valipour Dehnou, V., & Machado, D. (2023). Anodal tDCS over the left DLPFC but not M1 increases muscle activity and improves psychophysiological responses, cognitive function, and endurance performance in normobaric hypoxia: a randomized controlled trial. *BMC Neurosci*, *24*(1), 25. https://doi.org/10.1186/s12868-023-00794-4
Fadden, S., Ververs, P. M., & Wickens, C. D. (2001). Pathway HUDs: are they viable? *Hum Factors*, *43*(2), 173-193. https://doi.org/10.1518/001872001775900841
Fadhlillah, F., & Patil, S. (2018). Pharmacological and mechanical management of calcium channel blocker toxicity. *BMJ Case Rep*, *2018*. https://doi.org/10.1136/bcr-2018-225324
Fagenholz, P. J., Murray, A. F., Gutman, J. A., Findley, J. K., & Harris, N. S. (2007). New-onset anxiety disorders at high altitude. *Wilderness Environ Med*, *18*(4), 312-316. https://doi.org/10.1580/07-weme-br-102r1.1
Fan, A. P., Khalil, A. A., Fiebach, J. B., Zaharchuk, G., Villringer, A., Villringer, K., & Gauthier, C. J. (2020). Elevated brain oxygen extraction fraction measured by MRI susceptibility relates to perfusion status in acute ischemic stroke. *J Cereb Blood Flow Metab*, *40*(3), 539-551. https://doi.org/10.1177/0271678x19827944
Farah, M. J., Levine, D. N., & Calvanio, R. (1988). A case study of mental imagery deficit. *Brain Cogn*, *8*(2), 147-164. https://doi.org/10.1016/0278-2626(88)90046-2
Farkas, E., De Jong, G. I., de Vos, R. A., Jansen Steur, E. N., & Luiten, P. G. (2000). Pathological features of cerebral cortical capillaries are doubled in Alzheimer's disease and Parkinson's disease. *Acta Neuropathol*, *100*(4), 395-402. https://doi.org/10.1007/s004010000195
Felling, R. J., Snyder, M. J., Romanko, M. J., Rothstein, R. P., Ziegler, A. N., Yang, Z., Givogri, M. I., Bongarzone, E. R., & Levison, S. W. (2006). Neural stem/progenitor cells participate in the regenerative response to perinatal hypoxia/ischemia. *J Neurosci*, *26*(16), 4359-4369. https://doi.org/10.1523/jneurosci.1898-05.2006
Fernandes, C. J., Fernandes, C. J., & Chong, D. Y. (2020). Intraoperative Pulmonary Embolism

in an Adolescent Patient with Type III Spinal Muscular Atrophy: A Case Report. *JBJS Case Connect*, *10*(3), e20.00087. https://doi.org/10.2106/jbjs.Cc.20.00087
Fernández-Torre, J. L., Hernández-Hernández, M. A., Mato-Mañas, D., Marco de Lucas, E., Gómez-Ruiz, E., & Martín-Láez, R. (2021). Intracortical focal non-convulsive status epilepticus causing cerebral hypoxia and intracranial hypertension. *Epileptic Disord*, *23*(6), 911-916. https://doi.org/10.1684/epd.2021.1348
Ferrazzi, E., Bulfamante, G., Mezzopane, R., Barbera, A., Ghidini, A., & Pardi, G. (1999). Uterine Doppler velocimetry and placental hypoxic-ischemic lesion in pregnancies with fetal intrauterine growth restriction. *Placenta*, *20*(5-6), 389-394. https://doi.org/10.1053/plac.1999.0395
Field, A. J., & Cottrell, D. J. (2005). Postural hallucinations? An unusual presentation of anaemia. *Arch Dis Child*, *90*(11), 1192-1193. https://doi.org/10.1136/adc.2005.075994
Fields, A. W., & Shelton, A. L. (2006). Individual skill differences and large-scale environmental learning. *J Exp Psychol Learn Mem Cogn*, *32*(3), 506-515. https://doi.org/10.1037/0278-7393.32.3.506
Flemming, B., Seeliger, E., Wronski, T., Steer, K., Arenz, N., & Persson, P. B. (2000). Oxygen and renal hemodynamics in the conscious rat. *J Am Soc Nephrol*, *11*(1), 18-24. https://doi.org/10.1681/asn.V11118
Fossi, S., Amantini, A., Grippo, A., Cossu, C., Boni, N., & Pinto, F. (2004). Anoxic-ischemic alpha coma: prognostic significance of the incomplete variant. *Neurol Sci*, *24*(6), 397-400. https://doi.org/10.1007/s10072-003-0195-y
Fox, M. J., & Snider, G. L. (1979). Respiratory therapy. Current practice in ambulatory patients with chronic airflow obstruction. *Jama*, *241*(9), 937-940. https://doi.org/10.1001/jama.241.9.937
Frisaldi, E., Bottino, P., Fabbri, M., Trucco, M., De Ceglia, A., Esposito, N., Barbiani, D., Camerone, E. M., Costa, F., Destefanis, C., Milano, E., Massazza, G., Zibetti, M., Lopiano, L., & Benedetti, F. (2021). Effectiveness of a dance-physiotherapy combined intervention in Parkinson's disease: a randomized controlled pilot trial. *Neurol Sci*, *42*(12), 5045-5053. https://doi.org/10.1007/s10072-021-05171-9
Fujimoto, S., Suzuki, M., Sakamoto, K., Ibusuki, R., Tamura, K., Shiozawa, A., Ishii, S., Iikura, M., Izumi, S., & Sugiyama, H. (2019). Comparison of End-Tidal, Arterial, Venous, and Transcutaneous P(CO(2)). *Respir Care*, *64*(10), 1208-1214. https://doi.org/10.4187/respcare.06094
Funayama, M., & Takata, T. (2019). Forced person-following: a new type of stimulus-bound behavior. *Neurocase*, *25*(3-4), 75-79. https://doi.org/10.1080/13554794.2019.1638944
Furian, M., Latshang, T. D., Aeschbacher, S. S., Ulrich, S., Sooronbaev, T., Mirrakhimov, E. M., Aldashev, A., & Bloch, K. E. (2015). Cerebral oxygenation in highlanders with and without high-altitude pulmonary hypertension. *Exp Physiol*, *100*(8), 905-914. https://doi.org/10.1113/ep085200
Gao, S., Wang, T., Cao, L., Li, L., & Yang, S. (2023). Clinical effects of remimazolam alone or in combination with dexmedetomidine in patients receiving bronchoscopy and influences on postoperative cognitive function: a randomized-controlled trial. *Int J Clin Pharm*, *45*(1), 137-145. https://doi.org/10.1007/s11096-022-01487-4
Gao, W., She, J., Wang, M., Li, S., Chen, X., & Zhu, R. (2024). Argon gas poisoning leading to persistent memory impairment: A 2-year case report. *Medicine (Baltimore)*, *103*(24), e38545. https://doi.org/10.1097/md.0000000000038545
Gardner, C. L., & Burke, H. B. (2024). Individual heart failure patient variability in nocturnal hypoxia and arrhythmias. *Medicine (Baltimore)*, *103*(41), e40083. https://doi.org/10.1097/md.0000000000040083
Garegrat, R., Montaldo, P., Burgod, C., Pant, S., Mazlan, M., Palanisami, B., Chakkarapani, E., Woolfall, K., Johnson, S., Grant, P. E., Land, S., Mahmoud, M., Brady, T., Cornelius, V.,

Adams, E., Dorling, J., Aladangadi, N., Fleming, P., Pressler, R., . . . Thayyil, S. (2024). Whole-body hypothermia in mild neonatal encephalopathy: protocol for a multicentre phase III randomised controlled trial. *BMC Pediatr*, *24*(1), 460. https://doi.org/10.1186/s12887-024-04935-4

Gattis, M., & Holyoak, K. J. (1996). Mapping conceptual to spatial relations in visual reasoning. *J Exp Psychol Learn Mem Cogn*, *22*(1), 231-239. https://doi.org/10.1037//0278-7393.22.1.231

Gautier, H., & Bonora, M. (1982). Effects of hypoxia and respiratory stimulants in conscious intact and carotid denervated cats. *Bull Eur Physiopathol Respir*, *18*(4), 565-582.

Gazit, V., Ben-Abraham, R., Pick, C. G., Ben-Shlomo, I., & Katz, Y. (2003). Long-term neurobehavioral and histological damage in brain of mice induced by L-cysteine. *Pharmacol Biochem Behav*, *75*(4), 795-799. https://doi.org/10.1016/s0091-3057(03)00147-3

Gelineau-Morel, R., Dlamini, N., Bruss, J., Cohen, A. L., Robertson, A., Alexopoulos, D., Smyser, C. D., & Boes, A. D. (2024). Network localization of pediatric lesion-induced dystonia. *medRxiv*. https://doi.org/10.1101/2024.04.06.24305421

Genah, S., Angeli, A., Supuran, C. T., & Morbidelli, L. (2020). Effect of Carbonic Anhydrase IX inhibitors on human endothelial cell survival. *Pharmacol Res*, *159*, 104964. https://doi.org/10.1016/j.phrs.2020.104964

Gerhart, H. D., Seo, Y., Kim, J. H., Followay, B., Vaughan, J., Quinn, T., Gunstad, J., & Glickman, E. L. (2019). Investigating Effects of Cold Water Hand Immersion on Selective Attention in Normobaric Hypoxia. *Int J Environ Res Public Health*, *16*(16). https://doi.org/10.3390/ijerph16162859

Gessaga, E. C., & Urich, H. (1985). The cerebellum of epileptics. *Clin Neuropathol*, *4*(6), 238-245.

Ghetti, S., Lee, J. K., Sims, C. E., Demaster, D. M., & Glaser, N. S. (2010). Diabetic ketoacidosis and memory dysfunction in children with type 1 diabetes. *J Pediatr*, *156*(1), 109-114. https://doi.org/10.1016/j.jpeds.2009.07.054

Ghika, J., Villemure, J. G., Miklossy, J., Temperli, P., Pralong, E., Christen-Zaech, S., Pollo, C., Maeder, P., Bogousslavsky, J., & Vingerhoets, F. (2002). Postanoxic generalized dystonia improved by bilateral Voa thalamic deep brain stimulation. *Neurology*, *58*(2), 311-313. https://doi.org/10.1212/wnl.58.2.311

Gilmer, B., Kilkenny, J., Tomaszewski, C., & Watts, J. A. (2002). Hyperbaric oxygen does not prevent neurologic sequelae after carbon monoxide poisoning. *Acad Emerg Med*, *9*(1), 1-8. https://doi.org/10.1111/j.1553-2712.2002.tb01159.x

Giombolini, C., Notaristefano, S., Santucci, S., Savino, K., Pasquino, S., Ragni, T., & Ambrosio, G. (2005). Platypnea-orthodeoxia induced by fenestrated atrial septal aneurysm. *Ital Heart J*, *6*(2), 164-167.

Giovanello, K. S., Keane, M. M., & Verfaellie, M. (2006). The contribution of familiarity to associative memory in amnesia. *Neuropsychologia*, *44*(10), 1859-1865. https://doi.org/10.1016/j.neuropsychologia.2006.03.004

Giunta, M., Recchia, E. G., Capuano, P., Toscano, A., Attisani, M., Rinaldi, M., & Brazzi, L. (2023). Management of harlequin syndrome under ECPELLA support: A report of two cases and a proposed approach. *Ann Card Anaesth*, *26*(1), 97-101. https://doi.org/10.4103/aca.aca_176_21

Gleeson, J., Lederman, R., Herrman, H., Koval, P., Eleftheriadis, D., Bendall, S., Cotton, S. M., & Alvarez-Jimenez, M. (2017). Moderated online social therapy for carers of young people recovering from first-episode psychosis: study protocol for a randomised controlled trial. *Trials*, *18*(1), 27. https://doi.org/10.1186/s13063-016-1775-5

Glover, G. H., & Thomason, M. E. (2004). Improved combination of spiral-in/out images for BOLD fMRI. *Magn Reson Med*, *51*(4), 863-868. https://doi.org/10.1002/mrm.20016

Goetz, D. W., Hall, S. E., Harbison, R. W., & Reid, M. J. (1988). Pediatric acquired immunodeficiency syndrome with negative human immunodeficiency virus antibody response by enzyme-linked immunosorbent assay and Western blot. *Pediatrics*, *81*(3), 356-359.

Golan, H., Kashtuzki, I., Hallak, M., Sorokin, Y., & Huleihel, M. (2004). Maternal hypoxia during pregnancy induces fetal neurodevelopmental brain damage: partial protection by magnesium sulfate. *J Neurosci Res*, *78*(3), 430-441. https://doi.org/10.1002/jnr.20269

Goldbart, A., Row, B. W., Kheirandish, L., Schurr, A., Gozal, E., Guo, S. Z., Payne, R. S., Cheng, Z., Brittian, K. R., & Gozal, D. (2003). Intermittent hypoxic exposure during light phase induces changes in cAMP response element binding protein activity in the rat CA1 hippocampal region: water maze performance correlates. *Neuroscience*, *122*(3), 585-590. https://doi.org/10.1016/j.neuroscience.2003.08.054

Goldberg, K. B., & Ellis, D. W. (1997). Anoxic encephalopathy: a neurobehavioural study in rehabilitation. *Brain Inj*, *11*(10), 743-750. https://doi.org/10.1080/026990597123115

Goldstein, J. M., Seidman, L. J., Buka, S. L., Horton, N. J., Donatelli, J. L., Rieder, R. O., & Tsuang, M. T. (2000). Impact of genetic vulnerability and hypoxia on overall intelligence by age 7 in offspring at high risk for schizophrenia compared with affective psychoses. *Schizophr Bull*, *26*(2), 323-334. https://doi.org/10.1093/oxfordjournals.schbul.a033456

Gong, W., Liu, S., Xu, P., Fan, M., & Xue, M. (2015). Simultaneous Quantification of Diazepam and Dexamethasone in Plasma by High-Performance Liquid Chromatography with Tandem Mass Spectrometry and Its Application to a Pharmacokinetic Comparison between Normoxic and Hypoxic Rats. *Molecules*, *20*(4), 6901-6912. https://doi.org/10.3390/molecules20046901

Gotanda, H., Kameyama, Y., Yamaguchi, Y., Ishii, M., Hanaoka, Y., Yamamoto, H., Ogawa, S., Iijima, K., Akishita, M., & Ouchi, Y. (2013). Acute exogenous lipoid pneumonia caused by accidental kerosene ingestion in an elderly patient with dementia: a case report. *Geriatr Gerontol Int*, *13*(1), 222-225. https://doi.org/10.1111/j.1447-0594.2012.00896.x

Goussakov, I., Synowiec, S., Fabres, R. B., Almeida, G. D., Takada, S. H., Aksenov, D., & Drobyshevsky, A. (2024). Abnormal local cortical functional connectivity due to interneuron dysmaturation after neonatal intermittent hypoxia. *bioRxiv*. https://doi.org/10.1101/2024.06.04.596449

Gozal, D., Xue, Y. D., & Simakajornboon, N. (1999). Hypoxia induces c-Fos protein expression in NMDA but not AMPA glutamate receptor labeled neurons within the nucleus tractus solitarii of the conscious rat. *Neurosci Lett*, *262*(2), 93-96. https://doi.org/10.1016/s0304-3940(99)00065-8

Gozal, E., Shah, Z. A., Pequignot, J. M., Pequignot, J., Sachleben, L. R., Czyzyk-Krzeska, M. F., Li, R. C., Guo, S. Z., & Gozal, D. (2005). Tyrosine hydroxylase expression and activity in the rat brain: differential regulation after long-term intermittent or sustained hypoxia. *J Appl Physiol (1985)*, *99*(2), 642-649. https://doi.org/10.1152/japplphysiol.00880.2004

Grand, J., Kjaergaard, J., Nielsen, N., Friberg, H., Cronberg, T., Bro-Jeppesen, J., Karsdal, M. A., Nielsen, H. B., Frydland, M., Henriksen, K., Mattsson, N., Zetterberg, H., & Hassager, C. (2019). Serum tau fragments as predictors of death or poor neurological outcome after out-of-hospital cardiac arrest. *Biomarkers*, *24*(6), 584-591. https://doi.org/10.1080/1354750x.2019.1609580

Gray, W. P., May, K., & Sundström, L. E. (2002). Seizure induced dentate neurogenesis does not diminish with age in rats. *Neurosci Lett*, *330*(3), 235-238. https://doi.org/10.1016/s0304-3940(02)00810-8

Gregory, D. W., Schaffner, W., Alford, R. H., Kaiser, A. B., & McGee, Z. A. (1979). Sporadic cases of Legionnaires' disease: the expanding clinical spectrum. *Ann Intern Med*, *90*(4), 518-521. https://doi.org/10.7326/0003-4819-90-4-518

Gross, J. B., Blouin, R. T., Zandsberg, S., Conard, P. F., & Häussler, J. (1996). Effect of flumazenil on ventilatory drive during sedation with midazolam and alfentanil. *Anesthesiology*, *85*(4), 713-720. https://doi.org/10.1097/00000542-199610000-00005

Groswasser, Z., Cohen, M., & Costeff, H. (1989). Rehabilitation outcome after anoxic brain damage. *Arch Phys Med Rehabil*, *70*(3), 186-188.

Gu, R., Ye, G., Zhou, Y., & Jiang, Z. (2020). Combined mutations of NKX2-1 and surfactant protein C genes for refractory low oxyhemoglobin saturation and interstitial pneumonia: A case report. *Medicine (Baltimore)*, *99*(12), e19650. https://doi.org/10.1097/md.0000000000019650

Guerrini, C., Berlucchi, G., Bricolo, E., & Aglioti, S. M. (2003). Temporal modulation of spatial tactile extinction in right-brain-damaged patients. *J Cogn Neurosci*, *15*(4), 523-536. https://doi.org/10.1162/089892903321662912

Guillamondegui, O. D., Richards, J. E., Ely, E. W., Jackson, J. C., Archer, K. R., Norris, P. R., & Obremskey, W. T. (2011). Does hypoxia affect intensive care unit delirium or long-term cognitive impairment after multiple trauma without intracranial hemorrhage? *J Trauma*, *70*(4), 910-915. https://doi.org/10.1097/TA.0b013e3182114f18

Guillet, R., Edwards, A. D., Thoresen, M., Ferriero, D. M., Gluckman, P. D., Whitelaw, A., & Gunn, A. J. (2012). Seven- to eight-year follow-up of the CoolCap trial of head cooling for neonatal encephalopathy. *Pediatr Res*, *71*(2), 205-209. https://doi.org/10.1038/pr.2011.30

Guo, B., Yang, T., Nan, J., Huang, Q., Wang, C., & Xu, W. (2021). Efficacy and safety of Shenfu injection combined with sodium nitroprusside in the treatment of chronic heart failure in patients with coronary heart disease: A protocol of randomized controlled trial. *Medicine (Baltimore)*, *100*(7), e24414. https://doi.org/10.1097/md.0000000000024414

Hadanny, A., Golan, H., Fishlev, G., Bechor, Y., Volkov, O., Suzin, G., Ben-Jacob, E., & Efrati, S. (2015). Hyperbaric oxygen can induce neuroplasticity and improve cognitive functions of patients suffering from anoxic brain damage. *Restor Neurol Neurosci*, *33*(4), 471-486. https://doi.org/10.3233/rnn-150517

Hagino, N., Kobayashi, S., Tsutsumi, T., Horiuchi, S., Nagai, R., Setalo, G., & Dettrich, E. (2004). Vascular change of hippocampal capillary is associated with vascular change of retinal capillary in aging. *Brain Res Bull*, *62*(6), 537-547. https://doi.org/10.1016/s0361-9230(03)00082-0

Hagiwara, Y., & Inoue, N. (2015). First case of methemoglobinemia caused by a ClO2-based household product. *Pediatr Int*, *57*(6), 1182-1183. https://doi.org/10.1111/ped.12708

Hallam, T. M., Floyd, C. L., Folkerts, M. M., Lee, L. L., Gong, Q. Z., Lyeth, B. G., Muizelaar, J. P., & Berman, R. F. (2004). Comparison of behavioral deficits and acute neuronal degeneration in rat lateral fluid percussion and weight-drop brain injury models. *J Neurotrauma*, *21*(5), 521-539. https://doi.org/10.1089/089771504774129865

Hammond, J. B., Peraza, J., & Pierce, C. A. (2024). A case report of long-term effects of Delayed post-hypoxic leukoencephalopathy (DPHL) following benzodiazepine overdose. *Clin Neuropsychol*, *38*(7), 1756-1772. https://doi.org/10.1080/13854046.2024.2315746

Han, S. J., Lee, T. H., Yang, J. K., Cho, Y. S., Jung, Y., Chung, I. K., Park, S. H., Park, S., & Kim, S. J. (2019). Etomidate Sedation for Advanced Endoscopic Procedures. *Dig Dis Sci*, *64*(1), 144-151. https://doi.org/10.1007/s10620-018-5220-3

Hanif, S., Sinha, S., & Siddiqui, K. A. (2014). Electroencephalography findings in patients with acute post coronary artery bypass graft encephalopathy. *Neurosciences (Riyadh)*, *19*(4), 331-333.

Hayakawa, F., Okumura, A., Kato, T., Kuno, K., & Watanabe, K. (1997). Dysmature EEG pattern in EEGs of preterm infants with cognitive impairment: maturation arrest caused by prolonged mild CNS depression. *Brain Dev*, *19*(2), 122-125. https://doi.org/10.1016/s0387-7604(96)00491-3

Hayashi, R., Matsuzawa, Y., Kubo, K., & Kobayashi, T. (2005). Effects of simulated high altitude on event-related potential (P300) and auditory brain-stem responses. *Clin Neurophysiol*, *116*(6), 1471-1476. https://doi.org/10.1016/j.clinph.2005.02.020
Hayashida, Y., Hirakawa, H., Nakamura, T., & Maeda, M. (1996). Chemoreceptors in autonomic responses to hypoxia in conscious rats. *Adv Exp Med Biol*, *410*, 439-442. https://doi.org/10.1007/978-1-4615-5891-0_67
Hayhow, B., Velakoulis, D., Dewhurst, R., & Gaillard, F. (2015). Neuropsychiatric presentation following acute hypoxic-ischaemic encephalopathy. *Aust N Z J Psychiatry*, *49*(2), 188-189. https://doi.org/10.1177/0004867414555418
He, B., Wang, J., Qian, G., Hu, M., Qu, X., Wei, Z., Li, J., Chen, Y., Chen, H., Zhou, Q., & Wang, G. (2013). Analysis of high-altitude de-acclimatization syndrome after exposure to high altitudes: a cluster-randomized controlled trial. *PLoS One*, *8*(5), e62072. https://doi.org/10.1371/journal.pone.0062072
Heaton, R. K., Grant, I., McSweeny, A. J., Adams, K. M., & Petty, T. L. (1983). Psychologic effects of continuous and nocturnal oxygen therapy in hypoxemic chronic obstructive pulmonary disease. *Arch Intern Med*, *143*(10), 1941-1947.
Hecht, N., Fiss, I., Wolf, S., Barth, M., Vajkoczy, P., & Woitzik, J. (2011). Modified flow- and oxygen-related autoregulation indices for continuous monitoring of cerebral autoregulation. *J Neurosci Methods*, *201*(2), 399-403. https://doi.org/10.1016/j.jneumeth.2011.08.018
Heizati, M., Li, N., Shao, L., Yao, X., Wang, Y., Hong, J., Zhou, L., Zhang, D., Chang, G., & Abulikemu, S. (2017). Does increased serum d-lactate mean subclinical hyperpermeability of intestinal barrier in middle-aged nonobese males with OSA? *Medicine (Baltimore)*, *96*(49), e9144. https://doi.org/10.1097/md.0000000000009144
Helm, C., Labovsky, K., Thakrar, P. D., & Diaz, C. D. (2020). E-cigarette, or Vaping, Product Use-Associated Lung Injury-Lessons Learned: A Case Series. *A A Pract*, *14*(8), e01242. https://doi.org/10.1213/xaa.0000000000001242
Hendricks, D. M., Pollock, N. W., Natoli, M. J., & Vann, R. D. (2000). Mountaineering oxygen mask performance at 4572 m. *Aviat Space Environ Med*, *71*(11), 1142-1147.
Higgins, E. A., Chiles, W. D., McKenzie, J. M., Jennings, A. E., Funkhouser, G. E., & Mullen, S. R. (1979). Effects of altitude and two decongestant-antihistamine preparations on physiological functions and performance. *Aviat Space Environ Med*, *50*(2), 154-158.
Himmelbach, M., & Karnath, H. O. (2003). Goal-directed hand movements are not affected by the biased space representation in spatial neglect. *J Cogn Neurosci*, *15*(7), 972-980. https://doi.org/10.1162/089892903770007362
Ho, A. M., Chung, D. C., To, E. W., & Karmakar, M. K. (2004). Total airway obstruction during local anesthesia in a non-sedated patient with a compromised airway. *Can J Anaesth*, *51*(8), 838-841. https://doi.org/10.1007/bf03018461
Hobgood, C., Hevia, A., & Hinchey, P. (2004). Profiles in patient safety: when an error occurs. *Acad Emerg Med*, *11*(7), 766-770. https://doi.org/10.1197/j.aem.2003.11.023
Hoffmann, U., Smerecnik, M., Leyk, D., & Essfeld, D. (2005). Cardiovascular responses to apnea during dynamic exercise. *Int J Sports Med*, *26*(6), 426-431. https://doi.org/10.1055/s-2004-821113
Höhne, C., Arntz, E., Krebs, M. O., Boemke, W., & Kaczmarczyk, G. (2003). Nifedipine inhibits the hypoxia-induced decrease in plasma renin activity in conscious dogs. *J Physiol Pharmacol*, *54*(2), 137-149.
Holdstock, J. S., Parslow, D. M., Morris, R. G., Fleminger, S., Abrahams, S., Denby, C., Montaldi, D., & Mayes, A. R. (2008). Two case studies illustrating how relatively selective hippocampal lesions in humans can have quite different effects on memory. *Hippocampus*, *18*(7), 679-691. https://doi.org/10.1002/hipo.20427
Holley-Jones, M., Howard, V. B., Siegfried, C., & Trautman, D. (1992). Cardiopulmonary bypass

in the successful resuscitation of a 28-year-old woman with cardiac arrest. *J Emerg Nurs*, *18*(5), 377-379.

Holzgraefe, B., Andersson, C., Kalzén, H., von Bahr, V., Mosskin, M., Larsson, E. M., Palmér, K., Frenckner, B., & Larsson, A. (2017). Does permissive hypoxaemia during extracorporeal membrane oxygenation cause long-term neurological impairment?: A study in patients with H1N1-induced severe respiratory failure. *Eur J Anaesthesiol*, *34*(2), 98-103. https://doi.org/10.1097/eja.0000000000000544

Hood, A. M., Stotesbury, H., Kölbel, M., DeHaan, M., Downes, M., Kawadler, J. M., Sahota, S., Dimitriou, D., Inusa, B., Wilkey, O., Pelidis, M., Trompeter, S., Leigh, A., Younis, J., Drasar, E., Chakravorty, S., Rees, D. C., Height, S., Lawson, S., . . . Kirkham, F. J. (2021). Study of montelukast in children with sickle cell disease (SMILES): a study protocol for a randomised controlled trial. *Trials*, *22*(1), 690. https://doi.org/10.1186/s13063-021-05626-6

Hopkins, R. O., Gale, S. D., Johnson, S. C., Anderson, C. V., Bigler, E. D., Blatter, D. D., & Weaver, L. K. (1995). Severe anoxia with and without concomitant brain atrophy and neuropsychological impairments. *J Int Neuropsychol Soc*, *1*(5), 501-509. https://doi.org/10.1017/s135561770000059x

Hopkins, R. O., Tate, D. F., & Bigler, E. D. (2005). Anoxic versus traumatic brain injury: amount of tissue loss, not etiology, alters cognitive and emotional function. *Neuropsychology*, *19*(2), 233-242. https://doi.org/10.1037/0894-4105.19.2.233

Hopkins, R. O., Waldram, K., & Kesner, R. P. (2004). Sequences assessed by declarative and procedural tests of memory in amnesic patients with hippocampal damage. *Neuropsychologia*, *42*(14), 1877-1886. https://doi.org/10.1016/j.neuropsychologia.2004.05.008

Hopkins-Golightly, T., Raz, S., & Sander, C. J. (2003). Influence of slight to moderate risk for birth hypoxia on acquisition of cognitive and language function in the preterm infant: a cross-sectional comparison with preterm-birth controls. *Neuropsychology*, *17*(1), 3-13.

Horiuchi, A., Hayashi, T., Kikuchi, N., Hayashi, A., Fuseya, C., Shiozawa, T., & Konishi, I. (2012). Hypoxia upregulates ovarian cancer invasiveness via the binding of HIF-1α to a hypoxia-induced, methylation-free hypoxia response element of S100A4 gene. *Int J Cancer*, *131*(8), 1755-1767. https://doi.org/10.1002/ijc.27448

Hornbein, T. F. (1992). Long term effects of high altitude on brain function. *Int J Sports Med*, *13 Suppl 1*, S43-45. https://doi.org/10.1055/s-2007-1024589

Hortigüela, M. M., Martínez-Biarge, M., Conejo, D., Vega-Del-Val, C., & Arnaez, J. (2024). Motor, cognitive and behavioural outcomes after neonatal hypoxic-ischaemic encephalopathy. *An Pediatr (Engl Ed)*, *100*(2), 104-114. https://doi.org/10.1016/j.anpede.2024.01.009

Hövels-Gürich, H. H., Seghaye, M. C., Schnitker, R., Wiesner, M., Huber, W., Minkenberg, R., Kotlarek, F., Messmer, B. J., & Von Bernuth, G. (2002). Long-term neurodevelopmental outcomes in school-aged children after neonatal arterial switch operation. *J Thorac Cardiovasc Surg*, *124*(3), 448-458. https://doi.org/10.1067/mtc.2002.122307

Hu, X., Qiu, J., Grafe, M. R., Rea, H. C., Rassin, D. K., & Perez-Polo, J. R. (2003). Bcl-2 family members make different contributions to cell death in hypoxia and/or hyperoxia in rat cerebral cortex. *Int J Dev Neurosci*, *21*(7), 371-377. https://doi.org/10.1016/s0736-5748(03)00089-3

Huang, S., Wang, R., Guo, B., Ruan, H., Ma, J., Ren, L., & Liu, L. (2020). Fatal Methemoglobinemia Due to Acute Inhalation of Methyl Nitrite in an Industrial Accident. *J Forensic Sci*, *65*(3), 1016-1022. https://doi.org/10.1111/1556-4029.14275

Huchzermeyer, C., Albus, K., Gabriel, H. J., Otáhal, J., Taubenberger, N., Heinemann, U., Kovács, R., & Kann, O. (2008). Gamma oscillations and spontaneous network activity in the hippocampus are highly sensitive to decreases in pO2 and concomitant changes in

mitochondrial redox state. *J Neurosci*, *28*(5), 1153-1162. https://doi.org/10.1523/jneurosci.4105-07.2008
Hunt, R. W., Liley, H. G., Wagh, D., Schembri, R., Lee, K. J., Shearman, A. D., Francis-Pester, S., deWaal, K., Cheong, J. Y. L., Olischar, M., Badawi, N., Wong, F. Y., Osborn, D. A., Rajadurai, V. S., Dargaville, P. A., Headley, B., Wright, I., & Colditz, P. B. (2021). Effect of Treatment of Clinical Seizures vs Electrographic Seizures in Full-Term and Near-Term Neonates: A Randomized Clinical Trial. *JAMA Netw Open*, *4*(12), e2139604. https://doi.org/10.1001/jamanetworkopen.2021.39604
Hunter, G. R., & Young, G. B. (2010). Recovery of awareness after hyperacute hepatic encephalopathy with "flat" EEG, severe brain edema and deep coma. *Neurocrit Care*, *13*(2), 247-251. https://doi.org/10.1007/s12028-010-9391-9
Hynes, S. M., Fish, J., & Manly, T. (2014). Intensive working memory training: a single case experimental design in a patient following hypoxic brain damage. *Brain Inj*, *28*(13-14), 1766-1775. https://doi.org/10.3109/02699052.2014.954622
Ichii, M., Mori, K., Miyaoka, D., Sonoda, M., Tsujimoto, Y., Nakatani, S., Shoji, T., & Emoto, M. (2021). Suppression of thyrotropin secretion during roxadustat treatment for renal anemia in a patient undergoing hemodialysis. *BMC Nephrol*, *22*(1), 104. https://doi.org/10.1186/s12882-021-02304-2
Ikeda, T., Mishima, K., Aoo, N., Egashira, N., Iwasaki, K., Fujiwara, M., & Ikenoue, T. (2004). Combination treatment of neonatal rats with hypoxia-ischemia and endotoxin induces long-lasting memory and learning impairment that is associated with extended cerebral damage. *Am J Obstet Gynecol*, *191*(6), 2132-2141. https://doi.org/10.1016/j.ajog.2004.04.039
Ikeya, Y., Takeda, S., Tunakawa, M., Karakida, H., Toda, K., Yamaguchi, T., & Aburada, M. (2004). Cognitive improving and cerebral protective effects of acylated oligosaccharides in Polygala tenuifolia. *Biol Pharm Bull*, *27*(7), 1081-1085. https://doi.org/10.1248/bpb.27.1081
Incalzi, R. A., Corsonello, A., Trojano, L., Pedone, C., Acanfora, D., Spada, A., Izzo, O., & Rengo, F. (2008). Cognitive training is ineffective in hypoxemic COPD: a six-month randomized controlled trial. *Rejuvenation Res*, *11*(1), 239-250. https://doi.org/10.1089/rej.2007.0607
Ingraham, J. P., Forbes, M. E., Riddle, D. R., & Sonntag, W. E. (2008). Aging reduces hypoxia-induced microvascular growth in the rodent hippocampus. *J Gerontol A Biol Sci Med Sci*, *63*(1), 12-20. https://doi.org/10.1093/gerona/63.1.12
Inuzuka, Y., Edo, N., Araki, Y., Hoshi, T., Maruta, M., Nakamoto, N., & Suzuki, S. (2022). Decompression Illness Treated with the Hart-Kindwall Protocol in a Monoplace Chamber. *Am J Case Rep*, *23*, e935534. https://doi.org/10.12659/ajcr.935534
Iversen, O. H. (1985). What's new in proliferation and differentiation in malignant tumours? *Pathol Res Pract*, *180*(6), 705-710. https://doi.org/10.1016/s0344-0338(85)80053-4
Ivert, A., Lindblad Wollmann, C., & Pettersson, K. (2023). A Case Series on Pregnant Patients with Mild Covid-19 Infection and Signs of Severe Placental Insufficiency. *Case Rep Obstet Gynecol*, *2023*, 2018551. https://doi.org/10.1155/2023/2018551
Iwai, M., Stetler, R. A., Xing, J., Hu, X., Gao, Y., Zhang, W., Chen, J., & Cao, G. (2010). Enhanced oligodendrogenesis and recovery of neurological function by erythropoietin after neonatal hypoxic/ischemic brain injury. *Stroke*, *41*(5), 1032-1037. https://doi.org/10.1161/strokeaha.109.570325
Iwasaki, K., Egashira, N., Hatip-Al-Khatib, I., Akiyoshi, Y., Arai, T., Takagaki, Y., Watanabe, T., Mishima, K., & Fujiwara, M. (2006). Cerebral ischemia combined with beta-amyloid impairs spatial memory in the eight-arm radial maze task in rats. *Brain Res*, *1097*(1), 216-223. https://doi.org/10.1016/j.brainres.2006.04.073
Iwase, M., Izumizaki, M., Kanamaru, M., & Homma, I. (2004). Effects of hyperthermia on

ventilation and metabolism during hypoxia in conscious mice. *Jpn J Physiol*, *54*(1), 53-59. https://doi.org/10.2170/jjphysiol.54.53
Izraeli, S., Avgar, D., Almog, S., Shochat, I., Tochner, Z., Tamir, A., & Ribak, J. (1990). The effect of repeated doses of 30 mg pyridostigmine bromide on pilot performance in an A-4 flight simulator. *Aviat Space Environ Med*, *61*(5), 430-432.
Jana, S., Chakravarty, C., Taraphder, A., & Ramasubban, S. (2014). Successful use of sustained low efficiency dialysis in a case of severe phenobarbital poisoning. *Indian J Crit Care Med*, *18*(8), 530-532. https://doi.org/10.4103/0972-5229.138159
Jang, S. H., Hyun, Y. J., & Lee, H. D. (2016). Recovery of consciousness and an injured ascending reticular activating system in a patient who survived cardiac arrest: A case report. *Medicine (Baltimore)*, *95*(26), e4041. https://doi.org/10.1097/md.0000000000004041
Jang, S. H., & Kwon, H. G. (2016). Neural injury of the Papez circuit following hypoxic-ischemic brain injury: A case report. *Medicine (Baltimore)*, *95*(44), e5173. https://doi.org/10.1097/md.0000000000005173
Jang, S. H., & Kwon, H. G. (2017). Injury of ascending reticular activating system associated with delayed post-hypoxic leukoencephalopathy: a case report. *BMC Neurol*, *17*(1), 139. https://doi.org/10.1186/s12883-017-0917-z
Jang, S. H., & Kwon, H. G. (2024). Akinetic mutism and gait disturbance in a patient with delayed post-hypoxic leukoencephalopathy. *Neurocase*, *30*(1), 29-31. https://doi.org/10.1080/13554794.2024.2353125
Jang, S. H., Yeo, S. S., Cho, M. J., & Chung, W. K. (2024). Correlation between Thalamocortical Tract and Default Mode Network with Consciousness Levels in Hypoxic-Ischemic Brain Injury Patients: A Comparative Study Using the Coma Recovery Scale-Revised. *Med Sci Monit*, *30*, e943802. https://doi.org/10.12659/msm.943802
Jang, Y., Lee, E., Kim, Y., & Park, J. H. (2020). Number Processing Error as a Clinical Manifestation of Hemispatial Neglect Following Hypoxic Brain Injury: a Case Report. *Brain Neurorehabil*, *13*(3), e20. https://doi.org/10.12786/bn.2020.13.e20
Janowsky, D. S., Meacham, M. P., Blaine, J. D., Schoor, M., & Bozzetti, L. P. (1976). Simulated flying performance after marihuana intoxication. *Aviat Space Environ Med*, *47*(2), 124-128.
Janssen Daalen, J. M., Meinders, M. J., Giardina, F., Roes, K. C. B., Stunnenberg, B. C., Mathur, S., Ainslie, P. N., Thijssen, D. H. J., & Bloem, B. R. (2022). Multiple N-of-1 trials to investigate hypoxia therapy in Parkinson's disease: study rationale and protocol. *BMC Neurol*, *22*(1), 262. https://doi.org/10.1186/s12883-022-02770-7
Janssen Daalen, J. M., Meinders, M. J., Mathur, S., van Hees, H. W. H., Ainslie, P. N., Thijssen, D. H. J., & Bloem, B. R. (2024). Randomized controlled trial of intermittent hypoxia in Parkinson's disease: study rationale and protocol. *BMC Neurol*, *24*(1), 212. https://doi.org/10.1186/s12883-024-03702-3
Jebasingh, D., Devavaram Jackson, D., Venkataraman, S., Adeghate, E., & Starling Emerald, B. (2014). The protective effects of Cyperus rotundus on behavior and cognitive function in a rat model of hypoxia injury. *Pharm Biol*, *52*(12), 1558-1569. https://doi.org/10.3109/13880209.2014.908395
Jeffries, O., Patterson, S. D., & Waldron, M. (2019). The effect of severe and moderate hypoxia on exercise at a fixed level of perceived exertion. *Eur J Appl Physiol*, *119*(5), 1213-1224. https://doi.org/10.1007/s00421-019-04111-y
Jeong, J. H., Kwon, J. C., Chin, J., Yoon, S. J., & Na, D. L. (2002). Globus pallidus lesions associated with high mountain climbing. *J Korean Med Sci*, *17*(6), 861-863. https://doi.org/10.3346/jkms.2002.17.6.861
Jerez-Calero, A., Salvatierra-Cuenca, M. T., Benitez-Feliponi, Á., Fernández-Marín, C. E., Narbona-López, E., Uberos-Fernández, J., & Muñoz-Hoyos, A. (2020). Hypothermia

Plus Melatonin in Asphyctic Newborns: A Randomized-Controlled Pilot Study. *Pediatr Crit Care Med*, *21*(7), 647-655. https://doi.org/10.1097/pcc.0000000000002346

Jersey, S. L., Baril, R. T., McCarty, R. D., & Millhouse, C. M. (2010). Severe neurological decompression sickness in a U-2 pilot. *Aviat Space Environ Med*, *81*(1), 64-68. https://doi.org/10.3357/asem.2303.2010

Ji, L., Lyu, C. L., Feng, M., & Qiang, H. (2021). Asymptomatic Pulmonary Embolism After Shoulder Arthroscopy: Case Report and Literature Review. *Orthop Surg*, *13*(3), 1119-1125. https://doi.org/10.1111/os.12982

Jia, H., Chen, Y., Wang, Y., Jia, L., Tian, Y., & Jiang, H. (2023). The neuroprotective effect of electro-acupuncture on cognitive recovery for patients with mild traumatic brain injury: A randomized controlled clinical trial. *Medicine (Baltimore)*, *102*(6), e32885. https://doi.org/10.1097/md.0000000000032885

Jiang, J., Cong, X., Alageel, S., Dornseifer, U., Schilling, A. F., Hadjipanayi, E., Machens, H. G., & Moog, P. (2023). In Vitro Comparison of Lymphangiogenic Potential of Hypoxia Preconditioned Serum (HPS) and Platelet-Rich Plasma (PRP). *Int J Mol Sci*, *24*(3). https://doi.org/10.3390/ijms24031961

Jiang, W. I., Cao, Y., Xue, Y., Ji, Y., Winer, B. Y., Zhang, M., Singhal, N. S., Pierce, J. T., Chen, S., & Ma, D. K. (2024). Suppressing APOE4-induced mortality and cellular damage by targeting VHL. *bioRxiv*. https://doi.org/10.1101/2024.02.28.582664

Jin, H., Ding, Z., Lian, S., Zhao, Y., He, S., Zhou, L., Zhuoga, C., Wang, H., Xu, J., Du, A., Yan, G., & Sun, Y. (2020). Prevalence and Risk Factors of White Matter Lesions in Tibetan Patients Without Acute Stroke. *Stroke*, *51*(1), 149-153. https://doi.org/10.1161/strokeaha.119.027115

Jin, H. K., Yang, R. H., Chen, Y. F., Thornton, R. M., Jackson, R. M., & Oparil, S. (1989). Hemodynamic effects of arginine vasopressin in rats adapted to chronic hypoxia. *J Appl Physiol (1985)*, *66*(1), 151-160. https://doi.org/10.1152/jappl.1989.66.1.151

Jobe, J. B., Shukitt-Hale, B., Banderet, L. E., & Rock, P. B. (1991). Effects of dexamethasone and high terrestrial altitude on cognitive performance and affect. *Aviat Space Environ Med*, *62*(8), 727-732.

Johnson, K., Dhamrah, U., Amin, A., & Masci, J. (2021). Case Report: A Great Mimicker of Neoplasm: Pulmonary Cryptococcosis in an Immunocompetent Host After High-Altitude Descent. *Am J Trop Med Hyg*, *105*(2), 454-457. https://doi.org/10.4269/ajtmh.21-0238

Johnstone, B., & Bouman, D. E. (1992). Anoxic encephalopathy: a case study of an eight-year-old male with no residual cognitive deficits. *Int J Neurosci*, *62*(3-4), 207-213. https://doi.org/10.3109/00207459108999772

Jongruk, P., Soontaravarapas, N., Angkurawaranon, S., Kosarat, S., Khuwuthyakorn, V., Tantiprabha, W., Manopunya, S., Boonchooduang, N., Louthrenoo, O., Likhitweerawong, N., Katanyuwong, K., Sanguansermsri, C., & Wiwattanadittakul, N. (2024). Adjuvant High-Dose Erythropoietin With Delayed Therapeutic Hypothermia in Neonatal Hypoxic-Ischemic Encephalopathy. *Pediatr Neurol*, *161*, 268-276. https://doi.org/10.1016/j.pediatrneurol.2024.10.003

Jonin, P. Y., Besson, G., La Joie, R., Pariente, J., Belliard, S., Barillot, C., & Barbeau, E. J. (2018). Superior explicit memory despite severe developmental amnesia: In-depth case study and neural correlates. *Hippocampus*, *28*(12), 867-885. https://doi.org/10.1002/hipo.23010

Jordan, C. M., Whitman, R. D., & Harbut, M. (1997). Memory deficits and industrial toxicant exposure: a comparative study of hard metal, solvent and asbestos workers. *Int J Neurosci*, *90*(1-2), 113-128. https://doi.org/10.3109/00207459709000631

Joshi, A., Shrestha, P. S., Dangol, S., Shrestha, N. C., Poudyal, P., & Shrestha, A. (2017). Hemiconvulsion-Hemiplegia-Epilepsy Syndrome in a Girl Presented with Complex Partial Seizures. *Kathmandu Univ Med J (KUMJ)*, *15*(59), 256-260.

Kaandorp, J. J., Benders, M. J., Rademaker, C. M., Torrance, H. L., Oudijk, M. A., de Haan, T. R., Bloemenkamp, K. W., Rijken, M., van Pampus, M. G., Bos, A. F., Porath, M. M., Oetomo, S. B., Willekes, C., Gavilanes, A. W., Wouters, M. G., van Elburg, R. M., Huisjes, A. J., Bakker, S. C., van Meir, C. A., . . . Derks, J. B. (2010). Antenatal allopurinol for reduction of birth asphyxia induced brain damage (ALLO-Trial); a randomized double blind placebo controlled multicenter study. *BMC Pregnancy Childbirth*, *10*, 8. https://doi.org/10.1186/1471-2393-10-8

Kadoi, Y., Saito, S., Goto, F., & Fujita, N. (2001). Decrease in jugular venous oxygen saturation during normothermic cardiopulmonary bypass predicts short-term postoperative neurologic dysfunction in elderly patients. *J Am Coll Cardiol*, *38*(5), 1450-1455. https://doi.org/10.1016/s0735-1097(01)01584-4

Kahata, K., Hashino, S., Imamura, M., Mori, A., Kobayashi, S., & Asaka, M. (1997). Inhaled vancomycin-induced allergic reaction in decontamination of respiratory tracts for allogeneic bone marrow transplantation. *Bone Marrow Transplant*, *20*(11), 1001-1003. https://doi.org/10.1038/sj.bmt.1701007

Kam, C. A., Yoong, F. F., & Ganendran, A. (1978). Cortical blindness following hypoxia during cardiac arrest. *Anaesth Intensive Care*, *6*(2), 143-145. https://doi.org/10.1177/0310057x7800600209

Kamble, A., Khairkar, P., Kalantri, S. P., & Babhulkar, S. (2020). Fatal Suicidal Attempt by Deliberate Ingestion of Nicotine-containing Solution in Childhood-onset Depression Mediated through Internet Suicide Guideline: A Case Report. *Indian J Crit Care Med*, *24*(8), 719-721. https://doi.org/10.5005/jp-journals-10071-23524

Kang, P., Zhang, L., Liang, W., Zhu, Z., Liu, Y., Liu, X., & Yang, H. (2012). Medical evacuation management and clinical characteristics of 3,255 inpatients after the 2010 Yushu earthquake in China. *J Trauma Acute Care Surg*, *72*(6), 1626-1633. https://doi.org/10.1097/TA.0b013e3182479e07

Kaplan, C. P. (1999). Anoxic-hypotensive brain injury: neuropsychological performance at 1 month as an indicator of recovery. *Brain Inj*, *13*(4), 305-310. https://doi.org/10.1080/026990599121674

Kapur, N., Thompson, P., Kartsounis, L. D., & Abbott, P. (1999). Retrograde amnesia: clinical and methodological caveats. *Neuropsychologia*, *37*(1), 27-30. https://doi.org/10.1016/s0028-3932(98)00065-7

Kapus, G., Kertész, S., Gigler, G., Simó, A., Vegh, M., Barkóczy, J., Hársing, L. G., Szabó, G., & Lévay, G. (2004). Comparison of the AMPA antagonist action of new 2,3-benzodiazepines in vitro and their neuroprotective effects in vivo. *Pharm Res*, *21*(2), 317-323. https://doi.org/10.1023/b:pham.0000016245.74809.41

Karetzky, M. S. (1975). Asthma mortality: an analysis of one years experience, review of the literature and assessment of current modes of therapy. *Medicine (Baltimore)*, *54*(6), 471-484.

Karr, J. E., White, A. E., Leong, S. E., & Logan, T. K. (2025). The Neurobehavioral Symptom Inventory: Psychometric Properties and Symptom Comparisons in Women With and Without Brain Injuries Due to Intimate Partner Violence. *Assessment*, *32*(1), 102-118. https://doi.org/10.1177/10731911241236687

Kasper, J., Hermanns, M. I., Bantz, C., Maskos, M., Stauber, R., Pohl, C., Unger, R. E., & Kirkpatrick, J. C. (2011). Inflammatory and cytotoxic responses of an alveolar-capillary coculture model to silica nanoparticles: comparison with conventional monocultures. *Part Fibre Toxicol*, *8*(1), 6. https://doi.org/10.1186/1743-8977-8-6

Katsuragi, S., Ikeda, T., Date, I., Shingo, T., Yasuhara, T., Mishima, K., Aoo, N., Harada, K., Egashira, N., Iwasaki, K., Fujiwara, M., & Ikenoue, T. (2005). Implantation of encapsulated glial cell line-derived neurotrophic factor-secreting cells prevents long-lasting learning impairment following neonatal hypoxic-ischemic brain insult in rats.

*Am J Obstet Gynecol*, *192*(4), 1028-1037. https://doi.org/10.1016/j.ajog.2005.01.014
Katyal, N., Narula, N., George, P., Nattanamai, P., Newey, C. R., & Beary, J. M. (2018). Delayed Post-hypoxic Leukoencephalopathy: A Case Series and Review of the Literature. *Cureus*, *10*(4), e2481. https://doi.org/10.7759/cureus.2481
Kaur, S., Firdaus, S., Solano, J., Manjunath, S., & Ahmed, A. (2024). Incidental Finding of a Persistent Left Superior Vena Cava During Permanent Dual-Chamber Pacemaker Implantation: A Case Report. *Cureus*, *16*(11), e72865. https://doi.org/10.7759/cureus.72865
Kayser, B., & Mariani, B. (2022). Exceptional Performance in Competitive Ski Mountaineering: An Inertial Sensor Case Study. *Front Sports Act Living*, *4*, 854614. https://doi.org/10.3389/fspor.2022.854614
Ke, F., Dong, Z. H., Bu, F., Li, C. N., He, Q. T., Liu, Z. C., Lu, J., Yu, K., Wang, D. G., Xu, H. N., & Ye, C. T. (2024). Clostridium difficile infection following colon subtotal resection in a patient with gallstones: A case report and review of literature. *World J Gastrointest Surg*, *16*(9), 3048-3056. https://doi.org/10.4240/wjgs.v16.i9.3048
Kelleher, J. A. (1998). Old warp, new weft: weaving a new life fabric after anoxia. *Brain Inj*, *12*(4), 299-306. https://doi.org/10.1080/026990598122601
Kelly, M. C., & Loan, W. C. (1996). Use of a modified non-rebreathing mask during upper intestinal endoscopy. *Endoscopy*, *28*(8), 689-693. https://doi.org/10.1055/s-2007-1005578
Kelly, S. J., & Richards, J. E. (1998). Heart rate orienting and respiratory sinus arrhythmia development in rats exposed to alcohol or hypoxia. *Neurotoxicol Teratol*, *20*(2), 193-202. https://doi.org/10.1016/s0892-0362(97)00090-1
Kesner, R. P., & Hopkins, R. O. (2001). Short-term memory for duration and distance in humans: role of the hippocampus. *Neuropsychology*, *15*(1), 58-68. https://doi.org/10.1037//0894-4105.15.1.58
Keyghobadi, N., Roland, J., & Strobeck, C. (2005). Genetic differentiation and gene flow among populations of the alpine butterfly, Parnassius smintheus, vary with landscape connectivity. *Mol Ecol*, *14*(7), 1897-1909. https://doi.org/10.1111/j.1365-294X.2005.02563.x
Khan, F. A., McIntyre, C., Khan, A. M., & Maslov, A. (2020). Headache and Methemoglobinemia. *Headache*, *60*(1), 291-297. https://doi.org/10.1111/head.13696
Khot, S., Walker, M., Lacy, J. M., Oakes, P., & Longstreth, W. T., Jr. (2007). An unsuccessful trial of immunomodulatory therapy in delayed posthypoxic demyelination. *Neurocrit Care*, *7*(3), 253-256. https://doi.org/10.1007/s12028-007-0044-6
Kilicdag, H., Daglıoglu, K., Erdogan, S., Guzel, A., Sencar, L., Polat, S., & Zorludemir, S. (2013). The effect of levetiracetam on neuronal apoptosis in neonatal rat model of hypoxic ischemic brain injury. *Early Hum Dev*, *89*(5), 355-360. https://doi.org/10.1016/j.earlhumdev.2012.12.002
Kim, J. H., Kim, S. J., & Kim, H. Y. (2017). Right Hand Weakness and Headache During Ascent to Mount Everest: A Case of Cerebral Venous Infarction. *Neurologist*, *22*(3), 98-100. https://doi.org/10.1097/nrl.0000000000000121
King, J. A., Burgess, N., Hartley, T., Vargha-Khadem, F., & O'Keefe, J. (2002). Human hippocampus and viewpoint dependence in spatial memory. *Hippocampus*, *12*(6), 811-820. https://doi.org/10.1002/hipo.10070
Kinney, H. C., Korein, J., Panigrahy, A., Dikkes, P., & Goode, R. (1994). Neuropathological findings in the brain of Karen Ann Quinlan. The role of the thalamus in the persistent vegetative state. *N Engl J Med*, *330*(21), 1469-1475. https://doi.org/10.1056/nejm199405263302101
Kishiyama, M. M., Yonelinas, A. P., & Lazzara, M. M. (2004). The von Restorff effect in amnesia: the contribution of the hippocampal system to novelty-related memory

enhancements. *J Cogn Neurosci*, *16*(1), 15-23. https://doi.org/10.1162/089892904322755511
Klein, R., Soung, A., Sissoko, C., Nordvig, A., Canoll, P., Mariani, M., Jiang, X., Bricker, T., Goldman, J., Rosoklija, G., Arango, V., Underwood, M., Mann, J. J., Boon, A., Dowrk, A., & Boldrini, M. (2021). COVID-19 induces neuroinflammation and loss of hippocampal neurogenesis. *Res Sq*. https://doi.org/10.21203/rs.3.rs-1031824/v1
Knowlton, R. G., Ackerman, K. A., & Kaminsky, L. A. (1988). Physiological and performance comparisons of running flat and hill routes as applied to orienteering navigation. *J Sports Med Phys Fitness*, *28*(2), 189-193.
Koniaris, L. G., Kross, M. E., O'Malley, N., & Cornwell, E. E. (2000). Traumatic asphyxia complicated by unwitnessed cardiac arrest. *South Med J*, *93*(9), 905-908.
Kopelman, M. D., & Morton, J. (2015). Amnesia in an actor: Learning and re-learning of play passages despite severe autobiographical amnesia. *Cortex*, *67*, 1-14. https://doi.org/10.1016/j.cortex.2015.03.001
Kopelman, M. D., Reed, L. J., Marsden, P., Mayes, A. R., Jaldow, E., Laing, H., & Isaac, C. (2001). Amnesic syndrome and severe ataxia following the recreational use of 3,4-methylene-dioxymethamphetamine (MDMA, 'ecstasy') and other substances. *Neurocase*, *7*(5), 423-432. https://doi.org/10.1076/neur.7.5.423.16247
Koutroumanidis, M., Gratwicke, J., Sharma, S., Whelan, A., Tan, S. V., & Glover, G. (2021). Alpha coma EEG pattern in patients with severe COVID-19 related encephalopathy. *Clin Neurophysiol*, *132*(1), 218-225. https://doi.org/10.1016/j.clinph.2020.09.008
Kozora, E., Filley, C. M., Julian, L. J., & Cullum, C. M. (1999). Cognitive functioning in patients with chronic obstructive pulmonary disease and mild hypoxemia compared with patients with mild Alzheimer disease and normal controls. *Neuropsychiatry Neuropsychol Behav Neurol*, *12*(3), 178-183.
Krasney, J. A., Hajduczok, G., Akiba, C., McDonald, B. W., Pendergast, D. R., & Hong, S. K. (1984). Cardiovascular and renal responses to head-out water immersion in canine model. *Undersea Biomed Res*, *11*(2), 169-183.
Krikorian, R., & Layton, B. S. (1998). Implicit memory in posttraumatic stress disorder with amnesia for the traumatic event. *J Neuropsychiatry Clin Neurosci*, *10*(3), 359-362. https://doi.org/10.1176/jnp.10.3.359
Krishnaswamy, A., & Askari, A. T. (2007). A young woman with severe hypoxemia, electrocardiographic changes, and altered mental status. *Cleve Clin J Med*, *74*(7), 521-528. https://doi.org/10.3949/ccjm.74.7.521
Kulikov, V., Tregub, P., Parshin, D., Smirnova, Y., & Smirnov, K. (2022). Hypercapnic hypoxia improves cognitive and motor functions of children with cerebral palsy. *Neurol Res*, *44*(8), 738-747. https://doi.org/10.1080/01616412.2022.2051130
Kumar, R., Kapoor, R., Singh, J., Das, S., Sharma, A., Yanamandra, U., & Nair, V. (2019). Splenic Infarct on Exposure to Extreme High Altitude in Individuals with Sickle Trait: A Single-Center Experience. *High Alt Med Biol*, *20*(3), 215-220. https://doi.org/10.1089/ham.2018.0120
Kumaran, D., Hassabis, D., Spiers, H. J., Vann, S. D., Vargha-Khadem, F., & Maguire, E. A. (2007). Impaired spatial and non-spatial configural learning in patients with hippocampal pathology. *Neuropsychologia*, *45*(12), 2699-2711. https://doi.org/10.1016/j.neuropsychologia.2007.04.007
Kumral, A., Uysal, N., Tugyan, K., Sonmez, A., Yilmaz, O., Gokmen, N., Kiray, M., Genc, S., Duman, N., Koroglu, T. F., Ozkan, H., & Genc, K. (2004). Erythropoietin improves long-term spatial memory deficits and brain injury following neonatal hypoxia-ischemia in rats. *Behav Brain Res*, *153*(1), 77-86. https://doi.org/10.1016/j.bbr.2003.11.002
Kunisawa, T., Kurosawa, A., Hayashi, D., Takahashi, K., Kishi, M., & Iwasaki, H. (2011). Administration of dexmedetomidine alone during diagnostic cardiac catheterization in

adults with congenital heart disease: two case reports. *J Anesth*, *25*(4), 599-602. https://doi.org/10.1007/s00540-011-1174-8
Kurisu, K., Inada, S., Maeda, I., Ogawa, A., Iwase, S., Akechi, T., Morita, T., Oyamada, S., Yamaguchi, T., Imai, K., Nakahara, R., Kaneishi, K., Nakajima, N., Sumitani, M., & Yoshiuchi, K. (2022). A decision tree prediction model for a short-term outcome of delirium in patients with advanced cancer receiving pharmacological interventions: A secondary analysis of a multicenter and prospective observational study (Phase-R). *Palliat Support Care*, *20*(2), 153-158. https://doi.org/10.1017/s1478951521001565
Kwon, O. Y., Chung, S. P., Ha, Y. R., Yoo, I. S., & Kim, S. W. (2004). Delayed postanoxic encephalopathy after carbon monoxide poisoning. *Emerg Med J*, *21*(2), 250-251. https://doi.org/10.1136/emj.2002.002014
Lackmann, G. M., Töllner, U., & Mader, R. (1993). Serum enzyme activities in full-term asphyxiated and healthy newborns: enzyme kinetics during the first 144 hours of life. *Enzyme Protein*, *47*(3), 160-172. https://doi.org/10.1159/000468672
Lan, W. C., Priestley, M., Mayoral, S. R., Tian, L., Shamloo, M., & Penn, A. A. (2011). Sex-specific cognitive deficits and regional brain volume loss in mice exposed to chronic, sublethal hypoxia. *Pediatr Res*, *70*(1), 15-20. https://doi.org/10.1203/PDR.0b013e31821b98a3
Lang, J., Cui, X., Zhang, J., & Huang, Y. (2022). Dyspnea induced by hemidiaphragmatic paralysis after ultrasound-guided supraclavicular brachial plexus block in a morbidly obese patient. *Medicine (Baltimore)*, *101*(2), e28525. https://doi.org/10.1097/md.0000000000028525
Larsen, J. R., Kobborg, T., Shahim, P., Blennow, K., Rasmussen, L. S., & Zetterberg, H. (2021). Serum-neuroproteins, near-infrared spectroscopy, and cognitive outcome after beach-chair shoulder surgery: Observational cohort study analyses. *Acta Anaesthesiol Scand*, *65*(1), 26-33. https://doi.org/10.1111/aas.13691
Lawley, J. S., Macdonald, J. H., Oliver, S. J., & Mullins, P. G. (2017). Unexpected reductions in regional cerebral perfusion during prolonged hypoxia. *J Physiol*, *595*(3), 935-947. https://doi.org/10.1113/jp272557
Lazzaroni, M., & Bianchi-Porro, G. (1999). Premedication, preparation, and surveillance. *Endoscopy*, *31*(1), 2-8. https://doi.org/10.1055/s-1999-13642
Leach, J., & Almond, S. (1999). Ambient air, oxygen and nitrox effects on cognitive performance at altitude. *Appl Human Sci*, *18*(5), 175-179. https://doi.org/10.2114/jpa.18.175
Leconte, C., Bihel, E., Lepelletier, F. X., Bouët, V., Saulnier, R., Petit, E., Boulouard, M., Bernaudin, M., & Schumann-Bard, P. (2011). Comparison of the effects of erythropoietin and its carbamylated derivative on behaviour and hippocampal neurogenesis in mice. *Neuropharmacology*, *60*(2-3), 354-364. https://doi.org/10.1016/j.neuropharm.2010.09.025
Lee, G. W., Park, H. M., & Kang, M. H. (2019). Global brain ischemia in a dog with concurrent multiorgan dysfunction syndrome after bite wound trauma. *Acta Vet Scand*, *61*(1), 22. https://doi.org/10.1186/s13028-019-0458-0
Lee, H. M., Ruggoo, V., & Graudins, A. (2016). Intrathecal Clonidine Pump Failure Causing Acute Withdrawal Syndrome With 'Stress-Induced' Cardiomyopathy. *J Med Toxicol*, *12*(1), 134-138. https://doi.org/10.1007/s13181-015-0505-9
Lee, J., Kim, J., Kim, S., Kim, C., Yoon, T., & Kim, H. (2007). Removal of the laryngeal tube in children: anaesthetized compared with awake. *Br J Anaesth*, *98*(6), 802-805. https://doi.org/10.1093/bja/aem070
Lee, L. Y., & Morton, R. F. (1984). A comparison of breathing pattern between transient and steady state hypoxia in awake dogs. *Lung*, *162*(1), 15-26. https://doi.org/10.1007/bf02715624
Lefferts, W. K., Babcock, M. C., Tiss, M. J., Ives, S. J., White, C. N., Brutsaert, T. D., &

Heffernan, K. S. (2016). Effect of hypoxia on cerebrovascular and cognitive function during moderate intensity exercise. *Physiol Behav*, *165*, 108-118. https://doi.org/10.1016/j.physbeh.2016.07.003
Lefferts, W. K., Hughes, W. E., White, C. N., Brutsaert, T. D., & Heffernan, K. S. (2016). Effect of acute nitrate supplementation on neurovascular coupling and cognitive performance in hypoxia. *Appl Physiol Nutr Metab*, *41*(2), 133-141. https://doi.org/10.1139/apnm-2015-0400
Lehmann, S., Morand, S., James, C., & Schnider, A. (2007). Electrophysiological correlates of deficient encoding in a case of post-anoxic amnesia. *Neuropsychologia*, *45*(8), 1757-1766. https://doi.org/10.1016/j.neuropsychologia.2006.12.018
Lemoncello, R., Sohlberg, M. M., Fickas, S., & Prideaux, J. (2011). A randomised controlled crossover trial evaluating Television Assisted Prompting (TAP) for adults with acquired brain injury. *Neuropsychol Rehabil*, *21*(6), 825-846. https://doi.org/10.1080/09602011.2011.618661
Lenti, C., & Triulzi, F. (1996). Discordant clinical and neuroradiological features of congenital bilateral perisylvian syndrome in monozygotic female twins. *Ital J Neurol Sci*, *17*(4), 287-290. https://doi.org/10.1007/bf01997788
Leonard, W. R., Katzmarzyk, P. T., Stephen, M. A., & Ross, A. G. (1995). Comparison of the heart rate-monitoring and factorial methods: assessment of energy expenditure in highland and coastal Ecuadoreans. *Am J Clin Nutr*, *61*(5), 1146-1152. https://doi.org/10.1093/ajcn/61.4.1146
Leung, E., Javaid, S., Pelshaw, C. B., & Erlandson, E. (2020). Superimposed Guillain-Barré Syndrome (GBS) in pediatric hypoxic brain injury. *J Pediatr Rehabil Med*, *13*(1), 63-69. https://doi.org/10.3233/prm-180562
Levy-Zaks, A., Pollak, Y., & Ben-Pazi, H. (2014). Cerebral palsy risk factors and their impact on psychopathology. *Neurol Res*, *36*(1), 92-94. https://doi.org/10.1179/1743132813y.0000000290
Lewandowski, K. B. (1982). Strabismus as a possible sign of subclinical muscular dystrophy predisposing to rhabdomyolysis and myoglobinuria: a study of an affected family. *Can Anaesth Soc J*, *29*(4), 372-376. https://doi.org/10.1007/bf03007528
Lewis, J. S., & Jacobs, Z. G. (2020). Subtle case of dapsone-induced methaemoglobinaemia. *BMJ Case Rep*, *13*(8). https://doi.org/10.1136/bcr-2020-235403
Li, B., Ji, S., Peng, A., Yang, N., Zhao, X., Feng, P., Zhang, Y., & Chen, L. (2022). Development of a Gastrointestinal-Myoelectrical-Activity-Based Nomogram Model for Predicting the Risk of Mild Cognitive Impairment. *Biomolecules*, *12*(12). https://doi.org/10.3390/biom12121861
Li, H., Wang, J., Wang, P., Rao, Y., & Chen, L. (2016). Resveratrol Reverses the Synaptic Plasticity Deficits in a Chronic Cerebral Hypoperfusion Rat Model. *J Stroke Cerebrovasc Dis*, *25*(1), 122-128. https://doi.org/10.1016/j.jstrokecerebrovasdis.2015.09.004
Li, J., You, S. J., Xu, Y. N., Yuan, W., Shen, Y., Huang, J. Y., Xiong, K. P., & Liu, C. F. (2019). Cognitive impairment and sleep disturbances after minor ischemic stroke. *Sleep Breath*, *23*(2), 455-462. https://doi.org/10.1007/s11325-018-1709-4
Li, K., Liu, J., Zeng, Z., Kulyar, M. F., Wang, Y., Li, A., Bhutta, Z. A., Aqib, A. I., Shahzad, M., Li, J., & Qi, D. (2020). The Complete Genome of Probiotic Lactobacillus sakei Derived from Plateau Yak Feces. *Genes (Basel)*, *11*(12). https://doi.org/10.3390/genes11121527
Li, M., Sun, Z., Sun, H., Zhao, G., Leng, B., Shen, T., Xue, S., Hou, H., Li, Z., & Zhang, J. (2022). Paroxysmal slow wave events are associated with cognitive impairment in patients with obstructive sleep apnea. *Alzheimers Res Ther*, *14*(1), 200. https://doi.org/10.1186/s13195-022-01153-x
Li, Q., Michaud, M., Stewart, W., Schwartz, M., & Madri, J. A. (2008). Modeling the neurovascular niche: murine strain differences mimic the range of responses to chronic

hypoxia in the premature newborn. *J Neurosci Res*, *86*(6), 1227-1242. https://doi.org/10.1002/jnr.21597

Li, R. C., Row, B. W., Kheirandish, L., Brittian, K. R., Gozal, E., Guo, S. Z., Sachleben, L. R., Jr., & Gozal, D. (2004). Nitric oxide synthase and intermittent hypoxia-induced spatial learning deficits in the rat. *Neurobiol Dis*, *17*(1), 44-53. https://doi.org/10.1016/j.nbd.2004.05.006

Li, S., Wang, Y., Huang, X., Cao, J., & Yang, D. (2015). Diffuse alveolar hemorrhage from systemic lupus erythematosus misdiagnosed as high altitude pulmonary edema. *High Alt Med Biol*, *16*(1), 67-70. https://doi.org/10.1089/ham.2014.1094

Li, Y., Wang, J., Zhang, X., Ye, Q., Yang, Y., Cui, X., Feng, J., & Li, J. (2024). Correlation between serum inflammatory factors and cognitive function in patients with high-altitude polycythemia: A case-control study. *Medicine (Baltimore)*, *103*(17), e37983. https://doi.org/10.1097/md.0000000000037983

Liao, Y. H., Mündel, T., Yang, Y. T., Wei, C. C., & Tsai, S. C. (2019). Effects of periodic carbohydrate ingestion on endurance and cognitive performances during a 40-km cycling time-trial under normobaric hypoxia in well-trained triathletes. *J Sports Sci*, *37*(16), 1805-1815. https://doi.org/10.1080/02640414.2019.1595338

Lien, J., & Dibble, L. (2005). Systems model guided balance rehabilitation in an individual with declarative memory deficits and a total knee arthroplasty: a case report. *J Neurol Phys Ther*, *29*(1), 43-49. https://doi.org/10.1097/01.npt.0000282261.66702.ae

Lin, A., Mao, C., Rao, B., Zhao, H., Wang, Y., Yang, G., Lei, H., Xie, C., Huang, D., Deng, Y., Zhang, X., Wang, X., & Lu, J. (2023). Development and validation of nomogram including high altitude as a risk factor for COPD: A cross-sectional study based on Gansu population. *Front Public Health*, *11*, 1127566. https://doi.org/10.3389/fpubh.2023.1127566

Lin, H., Chang, C. P., Lin, H. J., Lin, M. T., & Tsai, C. C. (2012). Attenuating brain edema, hippocampal oxidative stress, and cognitive dysfunction in rats using hyperbaric oxygen preconditioning during simulated high-altitude exposure. *J Trauma Acute Care Surg*, *72*(5), 1220-1227. https://doi.org/10.1097/TA.0b013e318246ee70

Liscio, M., Adduci, A., Galbiati, S., Poggi, G., Sacchi, D., Strazzer, S., Castelli, E., & Flannery, J. (2008). Cognitive-behavioural stimulation protocol for severely brain-damaged patients in the post-acute stage in developmental age. *Disabil Rehabil*, *30*(4), 275-285. https://doi.org/10.1080/09638280701257023

Litman, R. S., Kottra, J. A., Berkowitz, R. J., & Ward, D. S. (1997). Breathing patterns and levels of consciousness in children during administration of nitrous oxide after oral midazolam premedication. *J Oral Maxillofac Surg*, *55*(12), 1372-1377; discussion 1378-1379. https://doi.org/10.1016/s0278-2391(97)90630-3

Liu, G., Su, Y., Jiang, M., Chen, W., Zhang, Y., Zhang, Y., & Gao, D. (2016). Electroencephalography reactivity for prognostication of post-anoxic coma after cardiopulmonary resuscitation: A comparison of quantitative analysis and visual analysis. *Neurosci Lett*, *626*, 74-78. https://doi.org/10.1016/j.neulet.2016.04.055

Liu, S., Chow, I. H. I., Lu, L., Ren, Y. M., Yang, H. L., Jian, S. Y., Ng, C. H., Ungvari, G. S., Wang, F., & Xiang, Y. T. (2020). Comparison of Sleep Disturbances Between Older Nursing Home Residents in High- and Low-Altitude Areas. *J Geriatr Psychiatry Neurol*, *33*(6), 370-376. https://doi.org/10.1177/0891988719892335

Liu, S., Sun, H., Wang, S., Liao, J., Yang, X., & Cai, Z. (2024). Isolated unilateral brachial plexus injury following carbon monoxide intoxication: a case report and literature review. *Front Neurol*, *15*, 1346353. https://doi.org/10.3389/fneur.2024.1346353

Liu, S., & Zhang, X. (2023). Amlodipine and Atropine for Hypoxia During One-Lung Ventilation: A Case Report. *Heart Surg Forum*, *26*(1), E048-050. https://doi.org/10.1532/hsf.5139

Liu, S. K., Chen, G., Yan, B., Huang, J., & Xu, H. (2019). Adverse Respiratory Events Increase

Post-anesthesia Care Unit Stay in China: A 2-year Retrospective Matched Cohort Study. *Curr Med Sci*, *39*(2), 325-329. https://doi.org/10.1007/s11596-019-2038-y

Liu, Z., Wang, X., Wu, Z., Yin, G., Chu, H., & Zhao, P. (2023). HBOT has a better cognitive outcome than NBH for patients with mild traumatic brain injury: A randomized controlled clinical trial. *Medicine (Baltimore)*, *102*(37), e35215. https://doi.org/10.1097/md.0000000000035215

Liutkeviciene, R., Lesauskaite, V., Asmoniene, V., Zaliūniene, D., & Jasinskas, V. (2010). Factors determining age-related macular degeneration: a current view. *Medicina (Kaunas)*, *46*(2), 89-94.

Lodziński, K., & Reszke, S. (1981). Gastritis necroticans in children. *Z Kinderchir*, *32*(1), 51-55. https://doi.org/10.1055/s-2008-1063233

Löhr, M., Tzouras, G., Molcanyi, M., Ernestus, R. I., Hampl, J. A., Fischer, J. H., Sahin, K., Arendt, T., & Härtig, W. (2008). Degeneration of cholinergic rat basal forebrain neurons after experimental subarachnoid hemorrhage. *Neurosurgery*, *63*(2), 336-344; discussion 344-335. https://doi.org/10.1227/01.Neu.0000320422.54985.6d

Lopez, M. G., Pandharipande, P., Morse, J., Shotwell, M. S., Milne, G. L., Pretorius, M., Shaw, A. D., Roberts, L. J., 2nd, & Billings, F. T. t. (2017). Intraoperative cerebral oxygenation, oxidative injury, and delirium following cardiac surgery. *Free Radic Biol Med*, *103*, 192-198. https://doi.org/10.1016/j.freeradbiomed.2016.12.039

Lorei, N. C., Ashley, R. L., Moberg, L. W., Butler, C. R., Seib, M. B., 2nd, & Hintz Merseal, C. N. (2024). Beetroot in deep waters: Navigating the risks of supplement-induced blackouts in aquatic training. *J Sci Med Sport*, *27*(10), 694-696. https://doi.org/10.1016/j.jsams.2024.06.009

LoVecchio, F., & Fulton, S. E. (2001). Ventricular fibrillation following inhalation of Glade Air Freshener. *Eur J Emerg Med*, *8*(2), 153-154. https://doi.org/10.1097/00063110-200106000-00014

Low, J. A., Galbraith, R. S., Muir, D. W., Killen, H. L., Pater, E. A., & Karchmar, E. J. (1983). Intrapartum fetal hypoxia: a study of long-term morbidity. *Am J Obstet Gynecol*, *145*(2), 129-134. https://doi.org/10.1016/0002-9378(83)90478-7

Low, J. A., Galbraith, R. S., Muir, D. W., Killen, H. L., Pater, E. A., & Karchmar, E. J. (1983). The predictive significance of biologic risk factors for deficits in children of a high-risk population. *Am J Obstet Gynecol*, *145*(8), 1059-1068. https://doi.org/10.1016/0002-9378(83)90867-0

Lu, S., John Portela, I. D., Martino, N., Ramos, R. B., Salinero, A. E., Smith, R. M., Zuloaga, K. L., & Adam, A. P. (2024). A transient brain endothelial translatome response to endotoxin is associated with mild cognitive changes post-shock in young mice. *bioRxiv*. https://doi.org/10.1101/2024.03.03.583191

Lu, Y., Zhuoga, C., Jin, H., Zhu, F., Zhao, Y., Ding, Z., He, S., Du, A., Xu, J., Luo, J., & Sun, Y. (2020). Characteristics of acute ischemic stroke in hospitalized patients in Tibet: a retrospective comparative study. *BMC Neurol*, *20*(1), 380. https://doi.org/10.1186/s12883-020-01957-0

Luewan, S., Sirichotiyakul, S., Charoenkwan, P., Phirom, K., & Tongsong, T. (2022). Hydrops Fetalis Associated with Fetal Hemoglobin H-Pakse Disease. *Fetal Diagn Ther*, *49*(11-12), 528-535. https://doi.org/10.1159/000528510

Lukawski, K., Nieradko, B., & Sieklucka-Dziuba, M. (2005). Effects of cadmium on memory processes in mice exposed to transient cerebral oligemia. *Neurotoxicol Teratol*, *27*(4), 575-584. https://doi.org/10.1016/j.ntt.2005.05.009

Luna, I. E., Kehlet, H., Olsen, R. M., Wede, H. R., Hoevsgaard, S. J., & Aasvang, E. K. (2020). Hypoxemia following hospital discharge after fast-track hip and knee arthroplasty - A prospective observational study subanalysis. *Acta Anaesthesiol Scand*, *64*(10), 1405-1413. https://doi.org/10.1111/aas.13671

Luo, D., Xu, J., Hu, L., Yu, L., Xie, L., & Li, J. (2018). Hyperbaric oxygen therapy to improve cognitive dysfunction and encephalatrophy induced by N(2)O for recreational use: a case report. *Neuropsychiatr Dis Treat*, *14*, 1963-1967. https://doi.org/10.2147/ndt.S170037

Luton, O. W., Stacey, B. S., Mellor, K., James, O. P., Williams, I. M., Warren, N., Egan, R. J., Bailey, D. M., & Lewis, W. G. (2023). Personal protective equipment-induced systemic hypercapnic hypoxaemia: translational implications for impaired cognitive-clinical functional performance. *Br J Surg*, *110*(5), 606-613. https://doi.org/10.1093/bjs/znad067

Maas, J. W., Jr., Indacochea, R. A., Muglia, L. M., Tran, T. T., Vogt, S. K., West, T., Benz, A., Shute, A. A., Holtzman, D. M., Mennerick, S., Olney, J. W., & Muglia, L. J. (2005). Calcium-stimulated adenylyl cyclases modulate ethanol-induced neurodegeneration in the neonatal brain. *J Neurosci*, *25*(9), 2376-2385. https://doi.org/10.1523/jneurosci.4940-04.2005

Mabry, S., Bradshaw, J. L., Gardner, J. J., Wilson, E. N., & Cunningham, R. (2024). Sex-dependent effects of chronic intermittent hypoxia: Implication for obstructive sleep apnea. *Res Sq*. https://doi.org/10.21203/rs.3.rs-3898670/v1

Mabry, S., Bradshaw, J. L., Gardner, J. J., Wilson, E. N., Sunuwar, J., Yeung, H., Shrestha, S., Cunningham, J. T., & Cunningham, R. L. (2024). The impact of chronic intermittent hypoxia on enzymatic activity in memory-associated brain regions of male and female rats. *Res Sq*. https://doi.org/10.21203/rs.3.rs-5449794/v1

Mabry, S., Wilson, E. N., Bradshaw, J. L., Gardner, J. J., Fadeyibi, O., Vera, E., Jr., Osikoya, O., Cushen, S. C., Karamichos, D., Goulopoulou, S., & Cunningham, R. L. (2023). Sex and age differences in social and cognitive function in offspring exposed to late gestational hypoxia. *Res Sq*. https://doi.org/10.21203/rs.3.rs-2507737/v1

Mader, E. C., Jr., Ramos, A. B., Cruz, R. A., & Branch, L. A. (2019). Full Recovery From Cocaine-Induced Toxic Leukoencephalopathy: Emphasizing the Role of Neuroinflammation and Brain Edema. *J Investig Med High Impact Case Rep*, *7*, 2324709619868266. https://doi.org/10.1177/2324709619868266

Magnin, E., Delchev, Y., Chopard, G., Berger, E., Vandel, P., Sechter, D., Rumbach, L., & Haffen, E. (2013). Transient improvement in sensorimotor conversion during post-anoxic encephalopathy with bilateral medial temporal ischemia. *Neurocase*, *19*(6), 576-582. https://doi.org/10.1080/13554794.2012.713488

Maguire, E. A., Frith, C. D., Rudge, P., & Cipolotti, L. (2005). The effect of adult-acquired hippocampal damage on memory retrieval: an fMRI study. *Neuroimage*, *27*(1), 146-152. https://doi.org/10.1016/j.neuroimage.2005.04.006

Maiti, P., Muthuraju, S., Ilavazhagan, G., & Singh, S. B. (2008). Hypobaric hypoxia induces dendritic plasticity in cortical and hippocampal pyramidal neurons in rat brain. *Behav Brain Res*, *189*(2), 233-243. https://doi.org/10.1016/j.bbr.2008.01.007

Maiwald, C. A., Niemarkt, H. J., Poets, C. F., Urschitz, M. S., König, J., Hummler, H., Bassler, D., Engel, C., & Franz, A. R. (2019). Effects of closed-loop automatic control of the inspiratory fraction of oxygen (FiO(2)-C) on outcome of extremely preterm infants - study protocol of a randomized controlled parallel group multicenter trial for safety and efficacy. *BMC Pediatr*, *19*(1), 363. https://doi.org/10.1186/s12887-019-1735-9

Malacrida, S., Katsuyama, Y., Droma, Y., Basnyat, B., Angelini, C., Ota, M., & Danieli, G. A. (2007). Association between human polymorphic DNA markers and hypoxia adaptation in Sherpa detected by a preliminary genome scan. *Ann Hum Genet*, *71*(Pt 5), 630-638. https://doi.org/10.1111/j.1469-1809.2007.00358.x

Malhotra, A., Menahem, S., Shekleton, P., & Gillam, L. (2009). Medical and ethical considerations in twin pregnancies discordant for serious cardiac disease. *J Perinatol*, *29*(10), 662-667. https://doi.org/10.1038/jp.2009.88

Malla, D. K., Singh, S., Basnet, Y., Adhikary, A., Chaudhary, V., Neupane, E., Ghimire, D.,

Baniya, A., O'Neill, V., & Karki, S. (2023). Endotracheal Intubation at 3,600 Meters Above Sea Level for High-Altitude Pulmonary Edema Followed by Helicopter Evacuation in Nepal. *Air Med J*, *42*(1), 58-60. https://doi.org/10.1016/j.amj.2022.11.001
Maller, A. I., Hankins, L. L., Yeakley, J. W., & Butler, I. J. (1998). Rolandic type cerebral palsy in children as a pattern of hypoxic-ischemic injury in the full-term neonate. *J Child Neurol*, *13*(7), 313-321. https://doi.org/10.1177/088307389801300702
Malm, T. M., Iivonen, H., Goldsteins, G., Keksa-Goldsteine, V., Ahtoniemi, T., Kanninen, K., Salminen, A., Auriola, S., Van Groen, T., Tanila, H., & Koistinaho, J. (2007). Pyrrolidine dithiocarbamate activates Akt and improves spatial learning in APP/PS1 mice without affecting beta-amyloid burden. *J Neurosci*, *27*(14), 3712-3721. https://doi.org/10.1523/jneurosci.0059-07.2007
Mancías-Guerra, C., Marroquín-Escamilla, A. R., González-Llano, O., Villarreal-Martínez, L., Jaime-Pérez, J. C., García-Rodríguez, F., Valdés-Burnes, S. L., Rodríguez-Romo, L. N., Barrera-Morales, D. C., Sánchez-Hernández, J. J., Cantú-Rodríguez, O. G., Gutiérrez-Aguirre, C. H., Gómez-De León, A., Elizondo-Riojas, G., Salazar-Riojas, R., & Gómez-Almaguer, D. (2014). Safety and tolerability of intrathecal delivery of autologous bone marrow nucleated cells in children with cerebral palsy: an open-label phase I trial. *Cytotherapy*, *16*(6), 810-820. https://doi.org/10.1016/j.jcyt.2014.01.008
Mancini, M., Filippelli, M., Seghieri, G., Iandelli, I., Innocenti, F., Duranti, R., & Scano, G. (1999). Respiratory muscle function and hypoxic ventilatory control in patients with type I diabetes. *Chest*, *115*(6), 1553-1562. https://doi.org/10.1378/chest.115.6.1553
Manigrasso, M., Protano, C., Martellucci, S., Mattei, V., Vitali, M., & Avino, P. (2019). Evaluation of the Submicron Particles Distribution Between Mountain and Urban Site: Contribution of the Transportation for Defining Environmental and Human Health Issues. *Int J Environ Res Public Health*, *16*(8). https://doi.org/10.3390/ijerph16081339
Mankad, J. P., Paulsen, K., & Shah, M. (2023). Partial Recovery in Toxic Leukoencephalopathy: Is It Really a Slow Improvement or a Warning Sign? *Cureus*, *15*(8), e42966. https://doi.org/10.7759/cureus.42966
Manley, C. N., Deepak, V., Ravikumar, N., Smith, A. K., Knight, A. K., Badell, M. L., Sidell, N., & Rajakumar, A. (2020). Transcription factor AP2A affects sFLT1 expression and decidualization in decidual stromal cells: Implications to preeclampsia pathology. *Pregnancy Hypertens*, *21*, 152-158. https://doi.org/10.1016/j.preghy.2020.06.001
Manning, L. (2002). Focal retrograde amnesia documented with matching anterograde and retrograde procedures. *Neuropsychologia*, *40*(1), 28-38. https://doi.org/10.1016/s0028-3932(01)00076-8
Marciante, A. B., Tadjalli, A., Burrowes, K. A., Oberto, J. R., Luca, E. K., Seven, Y. B., Nikodemova, M., Watters, J. J., Baker, T. L., & Mitchell, G. S. (2024). Microglia regulate motor neuron plasticity via reciprocal fractalkine/adenosine signaling. *bioRxiv*. https://doi.org/10.1101/2024.05.07.592939
Marques, A., Bourgois, N., Vidal, T., Ferrier, A., Mathais, S., Merlin, C., Valla, C., De Schlichting, E., Jean, B., Deffond, D., & Durif, F. (2018). Subacute corticobasal syndrome following internal carotid endarterectomy. *Rev Neurol (Paris)*, *174*(3), 157-161. https://doi.org/10.1016/j.neurol.2017.06.017
Martín, E. D., González-García, C., Milán, M., Fariñas, I., & Ceña, V. (2004). Stressor-related impairment of synaptic transmission in hippocampal slices from alpha-synuclein knockout mice. *Eur J Neurosci*, *20*(11), 3085-3091. https://doi.org/10.1111/j.1460-9568.2004.03801.x
Matchock, R. L., Susman, E. J., & Brown, F. M. (2004). Seasonal rhythms of menarche in the United States: correlates to menarcheal age, birth age, and birth month. *Womens Health Issues*, *14*(6), 184-192. https://doi.org/10.1016/j.whi.2004.07.007
Mateer, J. R., Olson, D. W., Stueven, H. A., & Aufderheide, T. P. (1993). Continuous pulse

oximetry during emergency endotracheal intubation. *Ann Emerg Med*, *22*(4), 675-679. https://doi.org/10.1016/s0196-0644(05)81846-3
Mathews, C. A., Bimson, B., Lowe, T. L., Herrera, L. D., Budman, C. L., Erenberg, G., Naarden, A., Bruun, R. D., Freimer, N. B., & Reus, V. I. (2006). Association between maternal smoking and increased symptom severity in Tourette's syndrome. *Am J Psychiatry*, *163*(6), 1066-1073. https://doi.org/10.1176/ajp.2006.163.6.1066
Matos, A. M. B., Dahy, F. E., de Moura, J. V. L., Marcusso, R. M. N., Gomes, A. B. F., Carvalho, F. M. M., Fernandes, G. B. P., Felix, A. C., Smid, J., Vidal, J. E., Frota, N. A. F., Casseb, J., Easton, A., Solomon, T., Witkin, S. S., Malta Romano, C., & de Oliveira, A. C. P. (2021). Subacute Cognitive Impairment in Individuals With Mild and Moderate COVID-19: A Case Series. *Front Neurol*, *12*, 678924. https://doi.org/10.3389/fneur.2021.678924
Matthey, S. (1996). Modification of perseverative behaviour in an adult with anoxic brain damage. *Brain Inj*, *10*(3), 219-227. https://doi.org/10.1080/026990596124539
Mayes, P. A., Campbell, L., Ricci, M. S., Plastaras, J. P., Dicker, D. T., & El-Deiry, W. S. (2005). Modulation of TRAIL-induced tumor cell apoptosis in a hypoxic environment. *Cancer Biol Ther*, *4*(10), 1068-1074. https://doi.org/10.4161/cbt.4.10.2255
Mayr, N. P., Hapfelmeier, A., Martin, K., Kurz, A., van der Starre, P., Babik, B., Mazzitelli, D., Lange, R., Wiesner, G., & Tassani-Prell, P. (2016). Comparison of sedation and general anaesthesia for transcatheter aortic valve implantation on cerebral oxygen saturation and neurocognitive outcome†. *Br J Anaesth*, *116*(1), 90-99. https://doi.org/10.1093/bja/aev294
Mazière, S., Pépin, J. L., Siyanko, N., Bioteau, C., Launois, S., Tamisier, R., Arnol, N., Lévy, P., Couturier, P., Bosson, J. L., & Gavazzi, G. (2014). Usefulness of oximetry for sleep apnea screening in frail hospitalized elderly. *J Am Med Dir Assoc*, *15*(6), 447.e449-414. https://doi.org/10.1016/j.jamda.2014.03.011
Mazza, S., Pépin, J. L., Naëgelé, B., Rauch, E., Deschaux, C., Ficheux, P., & Lévy, P. (2006). Driving ability in sleep apnoea patients before and after CPAP treatment: evaluation on a road safety platform. *Eur Respir J*, *28*(5), 1020-1028. https://doi.org/10.1183/09031936.06.00112905
McAuliffe, J. J., Joseph, B., Hughes, E., Miles, L., & Vorhees, C. V. (2008). Metallothionein I,II deficient mice do not exhibit significantly worse long-term behavioral outcomes following neonatal hypoxia-ischemia: MT-I,II deficient mice have inherent behavioral impairments. *Brain Res*, *1190*, 175-185. https://doi.org/10.1016/j.brainres.2007.11.038
McAuliffe, J. J., Joseph, B., & Vorhees, C. V. (2007). Isoflurane-delayed preconditioning reduces immediate mortality and improves striatal function in adult mice after neonatal hypoxia-ischemia. *Anesth Analg*, *104*(5), 1066-1077, tables of contents. https://doi.org/10.1213/01.ane.0000260321.62377.74
McCarthy, G. W., & Craig, K. D. (1995). Flying therapy for flying phobia. *Aviat Space Environ Med*, *66*(12), 1179-1184.
McClure, T. S., Phillips, J., Kernagis, D., Coleman, K., Chappe, E., Cutter, G. R., Egan, B., Norell, T., Stubbs, B. J., Bamman, M. M., & Koutnik, A. P. (2024). Ketone monoester attenuates oxygen desaturation during weighted ruck exercise under acute hypoxic exposure but does not impact cognitive performance. *Exp Physiol*, *109*(10), 1768-1781. https://doi.org/10.1113/ep091789
McClure, T. S., Phillips, J., Koutnik, A. P., Coleman, K., Chappe, E., Cutter, G. R., Egan, B., Norell, T., Stubbs, B. J., Bamman, M. M., & Kernagis, D. (2024). Ketone monoester attenuates declines in cognitive performance and oxygen saturation during acute severe hypoxic exposure under resting conditions. *Exp Physiol*, *109*(10), 1672-1682. https://doi.org/10.1113/ep091794
McComb, C., Varshney, N., & Lukowiak, K. (2005). Juvenile Lymnaea ventilate, learn and

remember differently than do adult Lymnaea. *J Exp Biol*, *208*(Pt 8), 1459-1467. https://doi.org/10.1242/jeb.01544
McCullough, M. L. (1976). The behavioral effects of prenatal hypoxia in the rat. *Dev Psychobiol*, *9*(4), 335-342. https://doi.org/10.1002/dev.420090406
McKhann, G. M., Grega, M. A., Borowicz, L. M., Jr., Bailey, M. M., Barry, S. J., Zeger, S. L., Baumgartner, W. A., & Selnes, O. A. (2005). Is there cognitive decline 1 year after CABG? Comparison with surgical and nonsurgical controls. *Neurology*, *65*(7), 991-999. https://doi.org/10.1212/01.wnl.0000175220.78475.99
McLean, A., Jr., Cardenas, D. D., Burgess, D., & Gamzu, E. (1991). Placebo-controlled study of pramiracetam in young males with memory and cognitive problems resulting from head injury and anoxia. *Brain Inj*, *5*(4), 375-380. https://doi.org/10.3109/02699059109008110
Medalia, A. A., Merriam, A. E., & Ehrenreich, J. H. (1991). The neuropsychological sequelae of attempted hanging. *J Neurol Neurosurg Psychiatry*, *54*(6), 546-548. https://doi.org/10.1136/jnnp.54.6.546
Mehdinezhad, H., Ghadir, S., Maleh, P. A., Akhondzadeh, A., & Baziboroun, M. (2024). Cryptococcal meningoencephalitis and pneumonia in a HIV positive patient: A case report. *Clin Case Rep*, *12*(8), e9196. https://doi.org/10.1002/ccr3.9196
Meixensberger, J., Renner, C., Simanowski, R., Schmidtke, A., Dings, J., & Roosen, K. (2004). Influence of cerebral oxygenation following severe head injury on neuropsychological testing. *Neurol Res*, *26*(4), 414-417. https://doi.org/10.1179/016164104225014094
Mercogliano, C., & Poddar, K. (2021). Long-Term Comorbid Neuropsychiatric Sequelae of Hypoxia at Birth. *Cureus*, *13*(1), e12687. https://doi.org/10.7759/cureus.12687
Michael, F. A., Peveling-Oberhag, J., Herrmann, E., Zeuzem, S., Bojunga, J., & Friedrich-Rust, M. (2021). Evaluation of the Integrated Pulmonary Index® during non-anesthesiologist sedation for percutaneous endoscopic gastrostomy. *J Clin Monit Comput*, *35*(5), 1085-1092. https://doi.org/10.1007/s10877-020-00563-2
Mikati, M. A., Zeinieh, M. P., Kurdi, R. M., Harb, S. A., El Hokayem, J. A., Daderian, R. H., Shamseddine, A., Obeid, M., Bitar, F. F., & El Sabban, M. (2005). Long-term effects of acute and of chronic hypoxia on behavior and on hippocampal histology in the developing brain. *Brain Res Dev Brain Res*, *157*(1), 98-102. https://doi.org/10.1016/j.devbrainres.2005.03.007
Millan, P. D., Kleiman, A. M., Friedman, J. F., Dunn, L. K., Gui, J. L., Bechtel, A. J., Collins, S. R., Huffmyer, J. L., Dwivedi, P., Wolpaw, J. T., Nemergut, E. C., Tsang, S., & Forkin, K. T. (2024). The impact of hindsight bias on the diagnosis of perioperative events by anesthesia providers: A multicenter randomized crossover study. *J Clin Anesth*, *97*, 111549. https://doi.org/10.1016/j.jclinane.2024.111549
Mindus, P., Cronholm, B., & Levander, S. E. (1975). Does piracetam counteract the ECT-induced memory dysfunctions in depressed patients? *Acta Psychiatr Scand*, *51*(5), 319-326. https://doi.org/10.1111/j.1600-0447.1975.tb00011.x
Miner, J. R., Bachman, A., Kosman, L., Teng, B., Heegaard, W., & Biros, M. H. (2005). Assessment of the onset and persistence of amnesia during procedural sedation with propofol. *Acad Emerg Med*, *12*(6), 491-496. https://doi.org/10.1197/j.aem.2005.01.011
Mishima, K., Ikeda, T., Yoshikawa, T., Aoo, N., Egashira, N., Xia, Y. X., Ikenoue, T., Iwasaki, K., & Fujiwara, M. (2004). Effects of hypothermia and hyperthermia on attentional and spatial learning deficits following neonatal hypoxia-ischemic insult in rats. *Behav Brain Res*, *151*(1-2), 209-217. https://doi.org/10.1016/j.bbr.2003.08.018
Mısırlıoğlu, M., Yıldızdaş, D., Aslan, N., Horoz, Ö., & Özden, Ö. (2019). Tracheal Rupture: A Rare Complication of Endotracheal Intubation. *Turk Arch Otorhinolaryngol*, *57*(3), 154-156. https://doi.org/10.5152/tao.2019.4225
Miskowiak, K. W., Damgaard, V., Schandorff, J. M., Macoveanu, J., Knudsen, G. M., Johansen, A., Plaven-Sigray, P., Svarer, C., Fussing, C. B., Cramer, K., Jørgensen, M. B., Kessing,

L. V., & Ehrenreich, H. (2024). Effects of cognitive training under hypoxia on cognitive proficiency and neuroplasticity in remitted patients with mood disorders and healthy individuals: ALTIBRAIN study protocol for a randomized controlled trial. *Trials*, *25*(1), 648. https://doi.org/10.1186/s13063-024-08463-5

Mitaki, S., Onoda, K., Ishihara, M., Nabika, Y., & Yamaguchi, S. (2012). Dysfunction of default-mode network in encephalopathy with a reversible corpus callosum lesion. *J Neurol Sci*, *317*(1-2), 154-156. https://doi.org/10.1016/j.jns.2012.02.020

Mittelbronn, M., Capper, D., Bader, B., Schittenhelm, J., Haybaeck, J., Weber, P., Meyermann, R., Kretzschmar, H. A., & Wietholter, H. (2008). Severe hypoxia and multiple infarctions resembling Creutzfeldt-Jakob disease. *Folia Neuropathol*, *46*(2), 149-153.

Miura, S., Ohyagi, Y., Ohno, M., Inoue, I., Ochi, H., Murai, H., Furuya, H., Yamada, T., & Kira, J. (2002). A patient with delayed posthypoxic demyelination: a case report of hyperbaric oxygen treatment. *Clin Neurol Neurosurg*, *104*(4), 311-314. https://doi.org/10.1016/s0303-8467(02)00019-7

Miyamoto, N., Pham, L. D., Hayakawa, K., Matsuzaki, T., Seo, J. H., Magnain, C., Ayata, C., Kim, K. W., Boas, D., Lo, E. H., & Arai, K. (2013). Age-related decline in oligodendrogenesis retards white matter repair in mice. *Stroke*, *44*(9), 2573-2578. https://doi.org/10.1161/strokeaha.113.001530

Mizuno, K., & Sumiyoshi, R. (1998). Air contamination of a closed anesthesia circuit. *Acta Anaesthesiol Scand*, *42*(1), 128-130. https://doi.org/10.1111/j.1399-6576.1998.tb05093.x

Molesworth, B. R., Bennett, L., & Kehoe, E. J. (2011). Promoting learning, memory, and transfer in a time-constrained, high hazard environment. *Accid Anal Prev*, *43*(3), 932-938. https://doi.org/10.1016/j.aap.2010.11.016

Mølgaard Nielsen, F., Lass Klitgaard, T., Crescioli, E., Rosborg Aagaard, S., Andreasen, A. S., Musaeus Poulsen, L., Siegemund, M., Craveiro Brøchner, A., Bestle, M. H., Andi Iversen, S., Brand, B. A., Laake, J. H., Grøfte, T., Hildebrandt, T., Lange, T., Perner, A., Lilleholt Schjørring, O., & Steen Rasmussen, B. (2021). Handling oxygenation targets in ICU patients with COVID-19-Protocol and statistical analysis plan in the HOT-COVID trial. *Acta Anaesthesiol Scand*, *65*(10), 1497-1504. https://doi.org/10.1111/aas.13956

Moller, J. T., Svennild, I., Johannessen, N. W., Jensen, P. F., Espersen, K., Gravenstein, J. S., Cooper, J. B., Djernes, M., & Johansen, S. H. (1993). Perioperative monitoring with pulse oximetry and late postoperative cognitive dysfunction. *Br J Anaesth*, *71*(3), 340-347. https://doi.org/10.1093/bja/71.3.340

Molloy, S., Soh, C., & Williams, T. L. (2006). Reversible delayed posthypoxic leukoencephalopathy. *AJNR Am J Neuroradiol*, *27*(8), 1763-1765.

Monroy-Gómez, J., & Torres-Fernández, O. (2020). Effects of the severe acute respiratory syndrome coronavirus (SARS-CoV) and the Middle East respiratory syndrome coronavirus (MERS-CoV) on the nervous system. What can we expect from SARS -CoV-2? *Biomedica*, *40*(Supl. 2), 173-179. https://doi.org/10.7705/biomedica.5682

Montes de Oca, E., Ball, G. E., & Spence, J. R. (2007). Diversity of Carabidae (Insecta, Coleoptera) in epiphytic Bromeliaceae in central Veracruz, Mexico. *Environ Entomol*, *36*(3), 560-568. https://doi.org/10.1603/0046-225x(2007)36[560:docici]2.0.co;2

Moon, S., Kim, S., Mankhong, S., Choi, S. H., Vandijck, M., Kostanjevecki, V., Jeong, J. H., Yoon, S. J., Park, K. W., Kim, E. J., Yoon, B., Kim, H. J., Jang, J. W., Hong, J. Y., Park, D. H., Shaw, L. M., & Kang, J. H. (2021). Alzheimer's cerebrospinal biomarkers from Lumipulse fully automated immunoassay: concordance with amyloid-beta PET and manual immunoassay in Koreans : CSF AD biomarkers measured by Lumipulse in Koreans. *Alzheimers Res Ther*, *13*(1), 22. https://doi.org/10.1186/s13195-020-00767-3

Morita, Y., Chin-Yee, I., Yu, P., Sibbald, W. J., & Martin, C. M. (2003). Critical oxygen delivery in conscious septic rats under stagnant or anemic hypoxia. *Am J Respir Crit Care Med*, *167*(6), 868-872. https://doi.org/10.1164/rccm.200205-490OC

Moulaert, V. R., Verbunt, J. A., van Heugten, C. M., Bakx, W. G., Gorgels, A. P., Bekkers, S. C., de Krom, M. C., & Wade, D. T. (2007). Activity and Life After Survival of a Cardiac Arrest (ALASCA) and the effectiveness of an early intervention service: design of a randomised controlled trial. *BMC Cardiovasc Disord*, *7*, 26. https://doi.org/10.1186/1471-2261-7-26
Mousa, W. F., Mowafi, H. A., Al-Metwalli, R. R., Al-Ghamdi, A. A., & Al-Gameel, H. Z. (2015). Preoperative mannitol infusion improves perioperative cerebral oxygen saturation and enhances postoperative recovery after laparoscopic cholecystectomy. *Saudi Med J*, *36*(10), 1199-1204. https://doi.org/10.15537/smj.2015.10.12105
Mujica-Parodi, L. R., Renelique, R., & Taylor, M. K. (2009). Higher body fat percentage is associated with increased cortisol reactivity and impaired cognitive resilience in response to acute emotional stress. *Int J Obes (Lond)*, *33*(1), 157-165. https://doi.org/10.1038/ijo.2008.218
Mundy, M. E., Downing, P. E., Dwyer, D. M., Honey, R. C., & Graham, K. S. (2013). A critical role for the hippocampus and perirhinal cortex in perceptual learning of scenes and faces: complementary findings from amnesia and FMRI. *J Neurosci*, *33*(25), 10490-10502. https://doi.org/10.1523/jneurosci.2958-12.2013
Muñoz, A. A., & Cavieres, L. A. (2006). A multi-species assessment of post-dispersal seed predation in the central Chilean Andes. *Ann Bot*, *98*(1), 193-201. https://doi.org/10.1093/aob/mcl087
Munoz, M., Chadwick, M., Perez-Hernandez, E., Vargha-Khadem, F., & Mishkin, M. (2011). Novelty preference in patients with developmental amnesia. *Hippocampus*, *21*(12), 1268-1276. https://doi.org/10.1002/hipo.20836
Murdoch, M., Chang, M., & McVicar, J. (1995). Central pontine myelinolysis after liver transplantation: a case report. *Transpl Int*, *8*(5), 399-402. https://doi.org/10.1007/bf00337174
Murray, R. M., Sham, P., Van Os, J., Zanelli, J., Cannon, M., & McDonald, C. (2004). A developmental model for similarities and dissimilarities between schizophrenia and bipolar disorder. *Schizophr Res*, *71*(2-3), 405-416. https://doi.org/10.1016/j.schres.2004.03.002
Mustafin, R. N., Shilova, I. V., Suslov, N. I., Kuvacheva, N. V., & Amelchenko, V. P. (2011). Nootropic activity of extracts from wild and cultivated Alfredia cernua. *Bull Exp Biol Med*, *150*(3), 333-335. https://doi.org/10.1007/s10517-011-1135-0
Mutch, W. A., Warrian, R. K., Eschun, G. M., Girling, L. G., Doiron, L., Cheang, M. S., & Lefevre, G. R. (2000). Biologically variable pulsation improves jugular venous oxygen saturation during rewarming. *Ann Thorac Surg*, *69*(2), 491-497. https://doi.org/10.1016/s0003-4975(99)01077-2
Myers, C. E., Deluca, J., Hopkins, R. O., & Gluck, M. A. (2006). Conditional discrimination and reversal in amnesia subsequent to hypoxic brain injury or anterior communicating artery aneurysm rupture. *Neuropsychologia*, *44*(1), 130-139. https://doi.org/10.1016/j.neuropsychologia.2005.03.026
Myers, C. E., Hopkins, R. O., DeLuca, J., Moore, N. B., Wolansky, L. J., Sumner, J. M., & Gluck, M. A. (2008). Learning and generalization deficits in patients with memory impairments due to anterior communicating artery aneurysm rupture or hypoxic brain injury. *Neuropsychology*, *22*(5), 681-686. https://doi.org/10.1037/0894-4105.22.5.681
Nadlewska, A., & Wiśniewska, R. J. (2009). Effect of MPEP on rat's behavioral activity in experimental episodes of hypoxia. *Adv Med Sci*, *54*(2), 277-282. https://doi.org/10.2478/v10039-009-0041-4
Nagy, E., Farkas, N., & Hollódy, K. (2020). Does Co-occurred Cerebral Palsy Change the Prognosis of West Syndrome? *Neuropediatrics*, *51*(1), 30-36. https://doi.org/10.1055/s-0039-1698450
Naismith, S., Winter, V., Gotsopoulos, H., Hickie, I., & Cistulli, P. (2004). Neurobehavioral

functioning in obstructive sleep apnea: differential effects of sleep quality, hypoxemia and subjective sleepiness. *J Clin Exp Neuropsychol*, *26*(1), 43-54. https://doi.org/10.1076/jcen.26.1.43.23929

Nascimento, F. A., Guebert, M., Rizelio, V., & da Rocha, S. F. (2016). Kleptomania following hypoxic-ischaemic damage to bilateral caudate nuclei. *BMJ Case Rep*, *2016*. https://doi.org/10.1136/bcr-2015-213710

Nassi, N., Piumelli, R., Lombardi, E., Landini, L., Donzelli, G., & de Martino, M. (2008). Comparison between pulse oximetry and transthoracic impedance alarm traces during home monitoring. *Arch Dis Child*, *93*(2), 126-132. https://doi.org/10.1136/adc.2007.118513

Navarrete-Opazo, A., Alcayaga, J., Testa, D., & Quinteros, A. L. (2016). Intermittent Hypoxia Does not Elicit Memory Impairment in Spinal Cord Injury Patients. *Arch Clin Neuropsychol*, *31*(4), 332-342. https://doi.org/10.1093/arclin/acw012

Nedergaard, H. K., Jensen, H. I., Stylsvig, M., Lauridsen, J. T., & Toft, P. (2016). Non-sedation versus sedation with a daily wake-up trial in critically ill patients recieving mechanical ventilation - effects on long-term cognitive function: Study protocol for a randomized controlled trial, a substudy of the NONSEDA trial. *Trials*, *17*(1), 269. https://doi.org/10.1186/s13063-016-1390-5

Nekoui, A., Tresierra del, C. E., Abdolmohammadi, S., Charbonneau, S., & Blaise, G. (2016). Recovery of brain function after cardiac arrest, case report and review. *Acta Anaesthesiol Belg*, *67*(1), 43-47.

Neppe, V. M. (2007). Differential cerebral cortical responsiveness examination in minimally conscious versus persistent vegetative states: a new role for neuropsychiatry and behavioral neurology. *J Neuropsychiatry Clin Neurosci*, *19*(4), 478-479. https://doi.org/10.1176/jnp.2007.19.4.478

Neubauer, J. A., & Edelman, N. H. (1984). Nonuniform brain blood flow response to hypoxia in unanesthetized cats. *J Appl Physiol Respir Environ Exerc Physiol*, *57*(6), 1803-1808. https://doi.org/10.1152/jappl.1984.57.6.1803

Neubauer, J. C., Dixon, J. P., & Herndon, C. M. (1988). Fatal pulmonary decompression sickness: a case report. *Aviat Space Environ Med*, *59*(12), 1181-1184.

Neubauer, R. A., Gottlieb, S. F., & Miale, A., Jr. (1992). Identification of hypometabolic areas in the brain using brain imaging and hyperbaric oxygen. *Clin Nucl Med*, *17*(6), 477-481. https://doi.org/10.1097/00003072-199206000-00010

Newman, S., Pugsley, W., Klinger, L., Harrison, M., Aveling, W., & Treasure, T. (1989). Neuropsychological consequences of circulatory arrest with hypothermia--a case report. *J Clin Exp Neuropsychol*, *11*(4), 529-538. https://doi.org/10.1080/01688638908400911

Ng, P. (2015). Splenic injury as a complication of colonoscopy: more common than we think? *BMJ Case Rep*, *2015*. https://doi.org/10.1136/bcr-2015-209707

Nickol, A. H., Leverment, J., Richards, P., Seal, P., Harris, G. A., Cleland, J., Dubowitz, G., Collier, D. J., Milledge, J., Stradling, J. R., & Morrell, M. J. (2006). Temazepam at high altitude reduces periodic breathing without impairing next-day performance: a randomized cross-over double-blind study. *J Sleep Res*, *15*(4), 445-454. https://doi.org/10.1111/j.1365-2869.2006.00558.x

Norvilaitė, K., Ramašauskaitė, D., Bartkevičienė, D., Žaliūnas, B., & Kurmanavičius, J. (2021). Doppler Ultrasonography of the Fetal Tibial Artery in High-Risk Pregnancy and Its Value in Predicting and Monitoring Fetal Hypoxia in IUGR Fetuses. *Medicina (Kaunas)*, *57*(10). https://doi.org/10.3390/medicina57101036

Nuño, N., Mäusezahl, D., Hattendorf, J., Verastegui, H., Ortiz, M., & Hartinger, S. M. (2022). Effectiveness of a home-environmental intervention package and an early child development intervention on child health and development in high-altitude rural communities in the Peruvian Andes: a cluster-randomised controlled trial. *Infect Dis*

*Poverty*, *11*(1), 66. https://doi.org/10.1186/s40249-022-00985-x
Nyffeler, T., Pflugshaupt, T., Hofer, H., Baas, U., Gutbrod, K., von Wartburg, R., Hess, C. W., & Müri, R. M. (2005). Oculomotor behaviour in simultanagnosia: a longitudinal case study. *Neuropsychologia*, *43*(11), 1591-1597. https://doi.org/10.1016/j.neuropsychologia.2005.01.011
Nymark, T. B., Hovland, A., Bjørnstad, H., & Nielsen, E. W. (2008). A young man with acute dilated cardiomyopathy associated with methylphenidate. *Vasc Health Risk Manag*, *4*(2), 477-479. https://doi.org/10.2147/vhrm.s2410
Ochozková, A., Mihalčíková, L., Yamamotová, A., & Šlamberová, R. (2021). Can prenatal methamphetamine exposure be considered a good animal model for ADHD? *Physiol Res*, *70*(S3), S431-s440. https://doi.org/10.33549/physiolres.934815
Ogasawara, K., Yamadate, K., Kobayashi, M., Endo, H., Fukuda, T., Yoshida, K., Terasaki, K., Inoue, T., & Ogawa, A. (2005). Effects of the free radical scavenger, edaravone, on the development of postoperative cognitive impairment in patients undergoing carotid endarterectomy. *Surg Neurol*, *64*(4), 309-313; discussion 313-304. https://doi.org/10.1016/j.surneu.2005.01.008
Ogura, T., Kobayashi, H., Suzuki, M., Sato, T., & Tomita, T. (1998). Posthyperventilation hypoxemia after methacholine inhalation. *Jpn J Physiol*, *48*(1), 39-47. https://doi.org/10.2170/jjphysiol.48.39
Ohrui, N., Takeuchi, A., Tong, A., Iwata, M., Nakamura, A., & Ohashi, K. (2005). Allergic rhinitis and ear pain in flight. *Ann Allergy Asthma Immunol*, *95*(4), 350-353. https://doi.org/10.1016/s1081-1206(10)61153-2
Okamoto, T., Hashimoto, K., Aoki, S., & Ohashi, M. (2007). Cerebral blood flow in patients with diffuse axonal injury--examination of the easy Z-score imaging system utility. *Eur J Neurol*, *14*(5), 540-547. https://doi.org/10.1111/j.1468-1331.2007.01742.x
Oliver, S. J., Macdonald, J. H., Harper Smith, A. D., Lawley, J. S., Gallagher, C. A., Di Felice, U., & Walsh, N. P. (2013). High altitude impairs in vivo immunity in humans. *High Alt Med Biol*, *14*(2), 144-149. https://doi.org/10.1089/ham.2012.1070
Ono, S., Yamafuji, T., Chaki, H., Todo, Y., Maekawa, M., Kitamura, K., Kimura, T., Nakada, Y., Mozumi, K., & Narita, H. (1995). A new cognition-enhancing agent, (R)-(-)-1-(benzo[b]thiophen-5-yl)- 2-[2-(N,N-diethylamino)ethoxy]ethanol hydrochloride. Effects on memory impairment in rats generated by cerebral embolization and basal forebrain lesions. *Biol Pharm Bull*, *18*(12), 1779-1783. https://doi.org/10.1248/bpb.18.1779
Ono, Y., Morifusa, M., Ikeda, S., Kunishige, C., & Tohma, Y. (2017). A case of non-cardiogenic pulmonary edema provoked by intravenous acetazolamide. *Acute Med Surg*, *4*(3), 349-352. https://doi.org/10.1002/ams2.279
O'Reilly, S. M., Grubb, N. R., & O'Carroll, R. E. (2003). In-hospital cardiac arrest leads to chronic memory impairment. *Resuscitation*, *58*(1), 73-79. https://doi.org/10.1016/s0300-9572(03)00114-x
Orešič, M., Hyötyläinen, T., Herukka, S. K., Sysi-Aho, M., Mattila, I., Seppänan-Laakso, T., Julkunen, V., Gopalacharyulu, P. V., Hallikainen, M., Koikkalainen, J., Kivipelto, M., Helisalmi, S., Lötjönen, J., & Soininen, H. (2011). Metabolome in progression to Alzheimer's disease. *Transl Psychiatry*, *1*(12), e57. https://doi.org/10.1038/tp.2011.55
Ortapamuk, H., & Naldoken, S. (2006). Brain perfusion abnormalities in chronic obstructive pulmonary disease: comparison with cognitive impairment. *Ann Nucl Med*, *20*(2), 99-106. https://doi.org/10.1007/bf02985621
O'Shaughnessy, J. A. (2002). Effects of epoetin alfa on cognitive function, mood, asthenia, and quality of life in women with breast cancer undergoing adjuvant chemotherapy. *Clin Breast Cancer*, *3 Suppl 3*, S116-120. https://doi.org/10.3816/cbc.2002.s.022
Otsuka, K., Norboo, T., Otsuka, Y., Higuchi, H., Hayajiri, M., Narushima, C., Sato, Y., Tsugoshi,

T., Murakami, S., Wada, T., Ishine, M., Okumiya, K., Matsubayashi, K., Yano, S., Chogyal, T., Angchuk, D., Ichihara, K., Cornélissen, G., & Halberg, F. (2005). Chronoecological health watch of arterial stiffness and neuro-cardio-pulmonary function in elderly community at high altitude (3524 m), compared with Japanese town. *Biomed Pharmacother*, *59 Suppl 1*(Suppl 1), S58-67. https://doi.org/10.1016/s0753-3322(05)80012-5

Owens, R. G., & Ghadiali, E. J. (1991). Judo as a possible cause of anoxic brain damage. A case report. *J Sports Med Phys Fitness*, *31*(4), 627-628.

Pannu, S. R., Haddad, T., Exline, M., Christman, J. W., Horowitz, J. C., Peters, J., Brock, G., Diaz, P., & Crouser, E. D. (2022). Rationale and design of a randomized controlled clinical trial; Titration of Oxygen Levels (TOOL) during mechanical ventilation. *Contemp Clin Trials*, *119*, 106811. https://doi.org/10.1016/j.cct.2022.106811

Papanek, P. E., & Raff, H. (1994). Chronic physiological increases in cortisol inhibit the vasopressin response to hypertonicity in conscious dogs. *Am J Physiol*, *267*(5 Pt 2), R1342-1349. https://doi.org/10.1152/ajpregu.1994.267.5.R1342

Pappas, A., Shankaran, S., McDonald, S. A., Vohr, B. R., Hintz, S. R., Ehrenkranz, R. A., Tyson, J. E., Yolton, K., Das, A., Bara, R., Hammond, J., & Higgins, R. D. (2015). Cognitive outcomes after neonatal encephalopathy. *Pediatrics*, *135*(3), e624-634. https://doi.org/10.1542/peds.2014-1566

Parisi, S., Gunasekara, R., Canale, C., Hasta, F., & Azizi, H. (2021). The Role of the Anterior Cingulate Cortex and Insular Cortex in Suicidal Memory and Intent. *Cureus*, *13*(7), e16335. https://doi.org/10.7759/cureus.16335

Park, C. S., Stojiljkovic, L., Milicic, B., Lin, B. F., & Dror, I. E. (2014). Training induces cognitive bias: the case of a simulation-based emergency airway curriculum. *Simul Healthc*, *9*(2), 85-93. https://doi.org/10.1097/SIH.0b013e3182a90304

Park, S. Y., Kim, S. M., Sung, J. J., Lee, K. M., Park, K. S., Kim, S. Y., Nam, H. W., & Lee, K. W. (2013). Nocturnal hypoxia in ALS is related to cognitive dysfunction and can occur as clusters of desaturations. *PLoS One*, *8*(9), e75324. https://doi.org/10.1371/journal.pone.0075324

Parkin, A. J., Miller, J., & Vincent, R. (1987). Multiple neuropsychological deficits due to anoxic encephalopathy: a case study. *Cortex*, *23*(4), 655-665. https://doi.org/10.1016/s0010-9452(87)80055-2

Patanè, S., Marte, F., Currò, A., & Cimino, C. (2010). Recurrent acute pulmonary embolism and paroxysmal atrial fibrillation associated with subclinical hyperthyroidism. *Int J Cardiol*, *142*(2), e25-26. https://doi.org/10.1016/j.ijcard.2008.11.179

Patanè, S., Marte, F., La Rosa, F. C., Di Bella, G., La Rocca, R., & Villari, S. A. (2010). Abnormal troponin I levels in acute pulmonary embolism without abnormal concentrations of D-dimer at admission. *Int J Cardiol*, *138*(1), 104-105. https://doi.org/10.1016/j.ijcard.2008.05.059

Patrician, A., Tymko, M. M., Caldwell, H. G., Howe, C. A., Coombs, G. B., Stone, R., Hamilton, A., Hoiland, R. L., & Ainslie, P. N. (2019). The Effect of an Expiratory Resistance Mask with Dead Space on Sleep, Acute Mountain Sickness, Cognition, and Ventilatory Acclimatization in Normobaric Hypoxia. *High Alt Med Biol*, *20*(1), 61-70. https://doi.org/10.1089/ham.2018.0074

Pavel, A. M., Rennie, J. M., de Vries, L. S., Mathieson, S. R., Livingstone, V., Finder, M., Foran, A., Shah, D. K., Pressler, R. M., Weeke, L. C., Dempsey, E. M., Murray, D. M., & Boylan, G. B. (2024). Temporal evolution of electrographic seizures in newborn infants with hypoxic-ischaemic encephalopathy requiring therapeutic hypothermia: a secondary analysis of the ANSeR studies. *Lancet Child Adolesc Health*, *8*(3), 214-224. https://doi.org/10.1016/s2352-4642(23)00296-1

Payne, L. B., Abdelazim, H., Hoque, M., Barnes, A., Mironovova, Z., Willi, C. E., Darden, J.,

Jenkins-Houk, C., Sedovy, M. W., Johnstone, S. R., & Chappell, J. C. (2023). A Soluble Platelet-Derived Growth Factor Receptor-β Originates via Pre-mRNA Splicing in the Healthy Brain and is Differentially Regulated during Hypoxia and Aging. *bioRxiv*. https://doi.org/10.1101/2023.02.03.527005

Pelidis, M. A., Kato, G. J., Resar, L. M., Dover, G. J., Nichols, D. G., Walker, L. K., & Casella, J. F. (1997). Successful treatment of life-threatening acute chest syndrome of sickle cell disease with venovenous extracorporeal membrane oxygenation. *J Pediatr Hematol Oncol*, *19*(5), 459-461. https://doi.org/10.1097/00043426-199709000-00010

Pentore, R., Venneri, A., & Nichelli, P. (1996). Accidental choke-cherry poisoning: early symptoms and neurological sequelae of an unusual case of cyanide intoxication. *Ital J Neurol Sci*, *17*(3), 233-235. https://doi.org/10.1007/bf01995689

Per, H., Kurtoğlu, S., Yağmur, F., Gümüş, H., Kumandaş, S., & Poyrazoğlu, M. H. (2007). Calcium carbide poisoning via food in childhood. *J Emerg Med*, *32*(2), 179-180. https://doi.org/10.1016/j.jemermed.2006.05.049

Perelle, I. B., & Ehrman, L. (1982). What is a lefthander? *Experientia*, *38*(10), 1256-1258. https://doi.org/10.1007/bf01959773

Périard, J. D., De Pauw, K., Zanow, F., & Racinais, S. (2018). Cerebrocortical activity during self-paced exercise in temperate, hot and hypoxic conditions. *Acta Physiol (Oxf)*, *222*(1). https://doi.org/10.1111/apha.12916

Perna, R., & Cooper, D. (2012). Perinatal cyanosis: long-term cognitive sequelae and behavioral consequences. *Appl Neuropsychol Child*, *1*(1), 48-52. https://doi.org/10.1080/09084282.2011.643946

Perrin, D., Mamet, J., Scarna, H., Roux, J. C., Bérod, A., & Dalmaz, Y. (2004). Long-term prenatal hypoxia alters maturation of brain catecholaminergic systems and motor behavior in rats. *Synapse*, *54*(2), 92-101. https://doi.org/10.1002/syn.20065

Perry, J. C., D'Almeida, V., Lima, M. M., Godoi, F. R., Vital, M. A., Oliveira, M. G., & Tufik, S. (2008). Intermittent hypoxia and sleep restriction: motor, cognitive and neurochemical alterations in rats. *Behav Brain Res*, *189*(2), 373-380. https://doi.org/10.1016/j.bbr.2008.01.014

Pettersen, J. A., Keith, J., Gao, F., Spence, J. D., & Black, S. E. (2017). CADASIL accelerated by acute hypotension: Arterial and venous contribution to leukoaraiosis. *Neurology*, *88*(11), 1077-1080. https://doi.org/10.1212/wnl.0000000000003717

Pfaff, J. A. R., Machegger, L., Trinka, E., & Mutzenbach, J. S. (2022). Unilateral delayed post-hypoxic leukoencephalopathy: a case report. *J Med Case Rep*, *16*(1), 480. https://doi.org/10.1186/s13256-022-03701-3

Pfeiffer, G., Pfeifer, R., & Isenmann, S. (2014). Cerebral hypoxia, missing cortical somatosensory evoked potentials and recovery of consciousness. *BMC Neurol*, *14*, 82. https://doi.org/10.1186/1471-2377-14-82

Pistoia, F., Sacco, S., Palmirotta, R., Onorati, P., Carolei, A., & Sarà, M. (2008). Mismatch of neurophysiological findings in partial recovery of consciousness: a case report. *Brain Inj*, *22*(7-8), 633-637. https://doi.org/10.1080/02699050802189693

Pitman, J. T., & Harris, N. S. (2012). Possible association with amphetamine usage and development of high altitude pulmonary edema. *Wilderness Environ Med*, *23*(4), 374-376. https://doi.org/10.1016/j.wem.2012.06.006

Pitt, B. R., & Lister, G. (1983). Pulmonary metabolic function in the awake lamb: effect of development and hypoxia. *J Appl Physiol Respir Environ Exerc Physiol*, *55*(2), 383-391. https://doi.org/10.1152/jappl.1983.55.2.383

Poli, S., Mbroh, J., Baron, J. C., Singhal, A. B., Strbian, D., Molina, C., Lemmens, R., Turc, G., Mikulik, R., Michel, P., Tatlisumak, T., Audebert, H. J., Dichgans, M., Veltkamp, R., Hüsing, J., Graessner, H., Fiehler, J., Montaner, J., Adeyemi, A. K., . . . Tuennerhoff, J. (2024). Penumbral Rescue by normobaric O = O administration in patients with ischemic

stroke and target mismatch proFile (PROOF): Study protocol of a phase IIb trial. *Int J Stroke*, *19*(1), 120-126. https://doi.org/10.1177/17474930231185275
Poll-The, B. T., Wanders, R. J., Ruiter, J. P., Ofman, R., Majoie, C. B., Barth, P. G., & Duran, M. (2004). Spastic diplegia and periventricular white matter abnormalities in 2-methyl-3-hydroxybutyryl-CoA dehydrogenase deficiency, a defect of isoleucine metabolism: differential diagnosis with hypoxic-ischemic brain diseases. *Mol Genet Metab*, *81*(4), 295-299. https://doi.org/10.1016/j.ymgme.2003.11.013
Pontolillo, M., Falasca, K., Vecchiet, J., & Ucciferri, C. (2021). It is Not Always COVID-19: Case Report about an Undiagnosed HIV Man with Dyspnea. *Curr HIV Res*, *19*(6), 548-551. https://doi.org/10.2174/1570162x19666210901134104
Poppert Cordts, K. M., Hall, T. A., Hartman, M. E., Luther, M., Wagner, A., Piantino, J., Guilliams, K. P., Guerriero, R. M., Jara, J., & Williams, C. N. (2020). Sleep Measure Validation in a Pediatric Neurocritical Care Acquired Brain Injury Population. *Neurocrit Care*, *33*(1), 196-206. https://doi.org/10.1007/s12028-019-00883-5
Porter, A. L. (1974). Effects of non-hydrogen-bonding anesthetics on memory in the chick. *Behav Biol*, *10*(3), 365-375. https://doi.org/10.1016/s0091-6773(74)91954-3
Pourié, G., Blaise, S., Trabalon, M., Nédélec, E., Guéant, J. L., & Daval, J. L. (2006). Mild, non-lesioning transient hypoxia in the newborn rat induces delayed brain neurogenesis associated with improved memory scores. *Neuroscience*, *140*(4), 1369-1379. https://doi.org/10.1016/j.neuroscience.2006.02.083
Powell, K. B., & Voeller, K. K. (2004). Prefrontal executive function syndromes in children. *J Child Neurol*, *19*(10), 785-797. https://doi.org/10.1177/08830738040190100801
Presti, A. L., Kishkurno, S. V., Slinko, S. K., Randis, T. M., Ratner, V. I., Polin, R. A., & Ten, V. S. (2006). Reoxygenation with 100% oxygen versus room air: late neuroanatomical and neurofunctional outcome in neonatal mice with hypoxic-ischemic brain injury. *Pediatr Res*, *60*(1), 55-59. https://doi.org/10.1203/01.pdr.0000223766.98760.88
Pretto, J. J., & McDonald, C. F. (2008). Acute oxygen therapy does not improve cognitive and driving performance in hypoxaemic COPD. *Respirology*, *13*(7), 1039-1044. https://doi.org/10.1111/j.1440-1843.2008.01392.x
Putzier, D. J., Padrick, K., Westfall, U. E., & Tanner, C. A. (1985). Diagnostic reasoning in critical care nursing. *Heart Lung*, *14*(5), 430-437.
Quamme, J. R., Yonelinas, A. P., Widaman, K. F., Kroll, N. E., & Sauvé, M. J. (2004). Recall and recognition in mild hypoxia: using covariance structural modeling to test competing theories of explicit memory. *Neuropsychologia*, *42*(5), 672-691. https://doi.org/10.1016/j.neuropsychologia.2003.09.008
Quigley, I., & Zafren, K. (2016). Subtle Cognitive Dysfunction in Resolving High Altitude Cerebral Edema Revealed by a Clock Drawing Test. *Wilderness Environ Med*, *27*(2), 256-258. https://doi.org/10.1016/j.wem.2015.12.006
Rabiei, M., Soori, T., Abiri, A., Farsi, Z., Shizarpour, A., & Pirjani, R. (2021). Maternal and fetal effects of COVID-19 virus on a complicated triplet pregnancy: a case report. *J Med Case Rep*, *15*(1), 87. https://doi.org/10.1186/s13256-020-02643-y
Raboud, M., Humm, A. M., Vivekanantham, H., & Suter, P. (2023). Transient central hypoxemia due to intermittent high-degree atrioventricular block in a heart-transplanted patient diagnosed during routine electroencephalography: a case report. *J Med Case Rep*, *17*(1), 3. https://doi.org/10.1186/s13256-022-03574-6
Raff, H., & Roarty, T. P. (1988). Renin, ACTH, and aldosterone during acute hypercapnia and hypoxia in conscious rats. *Am J Physiol*, *254*(3 Pt 2), R431-435. https://doi.org/10.1152/ajpregu.1988.254.3.R431
Rahmani, M., Belaidi, H., Benabdeljlil, M., Bouchhab, W., El Jazouli, N., El Brini, A., Aidi, S., Ouazzani, R. M., & El Alaoui Faris, M. (2013). Bilateral brachial plexus injury following acute carbon monoxide poisoning. *BMC Pharmacol Toxicol*, *14*, 61.

https://doi.org/10.1186/2050-6511-14-61
Rajan, J., Udupa, S., & Bharat, S. (2010). Hypoxia: can neuropsychological rehabilitation attenuate neuropsychological dysfunction. *Indian J Psychol Med*, *32*(1), 65-68. https://doi.org/10.4103/0253-7176.70544
Rajs, J. (1979). Histological diagnosis of myocardial injury. Comparison of hematoxylin-basic fuchsin-picric acid (HBFP)-stained sections obtained during autopsy with isolated viable rat cardiac myocytes exposed to anoxia. *Acta Pathol Microbiol Scand A*, *87a*(4), 289-297.
Rajs, J., Jones, D. P., & Jakobsson, S. W. (1978). Comparison of anoxic changes in isolated rat cardiac myocytes in suspension and in histological sections. *Acta Pathol Microbiol Scand A*, *86a*(5), 401-408. https://doi.org/10.1111/j.1699-0463.1978.tb02064.x
Raman, L., Hamilton, K. L., Gewirtz, J. C., & Rao, R. (2008). Effects of chronic hypoxia in developing rats on dendritic morphology of the CA1 subarea of the hippocampus and on fear-potentiated startle. *Brain Res*, *1190*, 167-174. https://doi.org/10.1016/j.brainres.2007.11.039
Raman, L., Tkac, I., Ennis, K., Georgieff, M. K., Gruetter, R., & Rao, R. (2005). In vivo effect of chronic hypoxia on the neurochemical profile of the developing rat hippocampus. *Brain Res Dev Brain Res*, *156*(2), 202-209. https://doi.org/10.1016/j.devbrainres.2005.02.013
Rambaud, C., & Guilleminault, C. (2012). Death, nasomaxillary complex, and sleep in young children. *Eur J Pediatr*, *171*(9), 1349-1358. https://doi.org/10.1007/s00431-012-1727-3
Ranasinghe, S., Or, G., Wang, E. Y., Ievins, A., McLean, M. A., Niell, C. M., Chau, V., Wong, P. K., Glass, H. C., Sullivan, J., & McQuillen, P. S. (2015). Reduced Cortical Activity Impairs Development and Plasticity after Neonatal Hypoxia Ischemia. *J Neurosci*, *35*(34), 11946-11959. https://doi.org/10.1523/jneurosci.2682-14.2015
Rankin, K. P., Kochamba, G. S., Boone, K. B., Petitti, D. B., & Buckwalter, J. G. (2003). Presurgical cognitive deficits in patients receiving coronary artery bypass graft surgery. *J Int Neuropsychol Soc*, *9*(6), 913-924. https://doi.org/10.1017/s1355617703960115
Rao, R., Comstock, B. A., Wu, T. W., Mietzsch, U., Mayock, D. E., Gonzalez, F. F., Wood, T. R., Heagerty, P. J., Juul, S. E., & Wu, Y. W. (2024). Time to Reaching Target Cooling Temperature and 2-year Outcomes in Infants with Hypoxic-Ischemic Encephalopathy. *J Pediatr*, *266*, 113853. https://doi.org/10.1016/j.jpeds.2023.113853
Rasmussen, P., Foged, E. M., Krogh-Madsen, R., Nielsen, J., Nielsen, T. R., Olsen, N. V., Petersen, N. C., Sørensen, T. A., Secher, N. H., & Lundby, C. (2010). Effects of erythropoietin administration on cerebral metabolism and exercise capacity in men. *J Appl Physiol (1985)*, *109*(2), 476-483. https://doi.org/10.1152/japplphysiol.00234.2010
Raveh, L., Weissman, B. A., Cohen, G., Alkalay, D., Rabinovitz, I., Sonego, H., & Brandeis, R. (2002). Caramiphen and scopolamine prevent soman-induced brain damage and cognitive dysfunction. *Neurotoxicology*, *23*(1), 7-17. https://doi.org/10.1016/s0161-813x(02)00005-0
Reményi, Á., Grósz, A., Szabó, S. A., Tótka, Z., Molnár, D., & Helfferich, F. (2018). Comparative study of the effect of bilastine and cetirizine on cognitive functions at ground level and at an altitude of 4,000 m simulated in hypobaric chamber: a randomized, double-blind, placebo-controlled, cross-over study. *Expert Opin Drug Saf*, *17*(9), 859-868. https://doi.org/10.1080/14740338.2018.1502268
Rether, C., Conen, A., Grossenbacher, M., & Albrich, W. C. (2011). A rare cause of pulmonary infiltrates one should be aware of: a case of daptomycin-induced acute eosinophilic pneumonia. *Infection*, *39*(6), 583-585. https://doi.org/10.1007/s15010-011-0148-y
Reynolds, R. D., Lickteig, J. A., Deuster, P. A., Howard, M. P., Conway, J. M., Pietersma, A., deStoppelaar, J., & Deurenberg, P. (1999). Energy metabolism increases and regional body fat decreases while regional muscle mass is spared in humans climbing Mt. Everest. *J Nutr*, *129*(7), 1307-1314. https://doi.org/10.1093/jn/129.7.1307

Rice, G. M., Vacchiano, C. A., Moore, J. L., Jr., & Anderson, D. W. (2003). Incidence of decompression sickness in hypoxia training with and without 30-min O2 prebreathe. *Aviat Space Environ Med*, *74*(1), 56-61.
Rice, N. J., Edwards, M. G., Schindler, I., Punt, T. D., McIntosh, R. D., Humphreys, G. W., Lestou, V., & Milner, A. D. (2008). Delay abolishes the obstacle avoidance deficit in unilateral optic ataxia. *Neuropsychologia*, *46*(5), 1549-1557. https://doi.org/10.1016/j.neuropsychologia.2008.01.012
Richalet, J. P., Larmignat, P., & Poignard, P. (2020). Transient Cerebral Ischemia at High Altitude and Hyper-Responsiveness to Hypoxia. *High Alt Med Biol*, *21*(1), 105-108. https://doi.org/10.1089/ham.2019.0100
Richardson, A. E., & Bereen, F. J. (1977). Effect of piracetam on level of consciousness after neurosurgery. *Lancet*, *2*(8048), 1110-1111. https://doi.org/10.1016/s0140-6736(77)90550-5
Riddle, A., Srivastava, T., Wang, K., Tellez, E., O'Neill, H., Gong, X., O'Niel, A., Bell, J. A., Raber, J., Lattal, M., Maylie, J., & Back, S. A. (2024). Mild neonatal hypoxia disrupts adult hippocampal learning and memory and is associated with CK2-mediated dysregulation of synaptic calcium-activated potassium channel KCNN2. *bioRxiv*. https://doi.org/10.1101/2024.07.10.602558
Riedel, W. J., & Jolles, J. (1996). Cognition enhancers in age-related cognitive decline. *Drugs Aging*, *8*(4), 245-274. https://doi.org/10.2165/00002512-199608040-00003
Rizzi, M., Airoldi, A., Cristiano, A., Frassanito, F., Macaluso, C., Vanni, S., & Legnani, D. (2016). Oxygen therapy in COPD patients with isolated nocturnal hypoxemia; comparison of quality of life and sleep between bronchitis and emphysema phenotype: A prospective observational study. *Eur J Intern Med*, *34*, 78-84. https://doi.org/10.1016/j.ejim.2016.08.035
Robertson, C. L., Clark, R. S., Dixon, C. E., Alexander, H. L., Graham, S. H., Wisniewski, S. R., Marion, D. W., Safar, P. J., & Kochanek, P. M. (2000). No long-term benefit from hypothermia after severe traumatic brain injury with secondary insult in rats. *Crit Care Med*, *28*(9), 3218-3223. https://doi.org/10.1097/00003246-200009000-00017
Robertsson Grossmann, K., Eriksson Westblad, M., Blennow, M., & Lindström, K. (2023). Outcome at early school age and adolescence after hypothermia-treated hypoxic-ischaemic encephalopathy: an observational, population-based study. *Arch Dis Child Fetal Neonatal Ed*, *108*(3), 295-301. https://doi.org/10.1136/archdischild-2022-324418
Robinson, S., Petelenz, K., Li, Q., Cohen, M. L., Dechant, A., Tabrizi, N., Bucek, M., Lust, D., & Miller, R. H. (2005). Developmental changes induced by graded prenatal systemic hypoxic-ischemic insults in rats. *Neurobiol Dis*, *18*(3), 568-581. https://doi.org/10.1016/j.nbd.2004.10.024
Robl, J., Vutthikraivit, W., Horwitz, P., & Panaich, S. (2022). Percutaneous closure of patent foramen ovale for treatment of hypoxemia: A case series and physiology review. *Catheter Cardiovasc Interv*, *100*(3), 471-475. https://doi.org/10.1002/ccd.30317
Rodricks, C. L., Rose, I. A., Camm, E. J., Jenkin, G., Miller, S. L., & Gibbs, M. E. (2004). The effect of prenatal hypoxia and malnutrition on memory consolidation in the chick. *Brain Res Dev Brain Res*, *148*(1), 113-119. https://doi.org/10.1016/j.devbrainres.2003.10.008
Rodríguez-Fanjul, J., Durán Fernández-Feijóo, C., Lopez-Abad, M., Lopez Ramos, M. G., Balada Caballé, R., Alcántara-Horillo, S., & Camprubí Camprubí, M. (2017). Neuroprotection with hypothermia and allopurinol in an animal model of hypoxic-ischemic injury: Is it a gender question? *PLoS One*, *12*(9), e0184643. https://doi.org/10.1371/journal.pone.0184643
Rodríguez-Hernández, A., Torné, R., Blanco Ibáñez de Opacua, A., Brugada-Bellsolà, F., Remollo, S., Domínguez, C. J., & Rimbau, J. M. (2020). Amateur Endurance Athletes: At

Higher Risk of Suffering Dural Arteriovenous Fistulas? Report of 3 Cases. *World Neurosurg*, *140*, 32-36. https://doi.org/10.1016/j.wneu.2020.05.035

Roehrs, T., Merrion, M., Pedrosi, B., Stepanski, E., Zorick, F., & Roth, T. (1995). Neuropsychological function in obstructive sleep apnea syndrome (OSAS) compared to chronic obstructive pulmonary disease (COPD). *Sleep*, *18*(5), 382-388. https://doi.org/10.1093/sleep/18.5.382

Rogers, A. J., Denk, L. D., & Wax, P. M. (2004). Catastrophic brain injury after nicotine insecticide ingestion. *J Emerg Med*, *26*(2), 169-172. https://doi.org/10.1016/j.jemermed.2003.05.006

Rogers, E. E., Bonifacio, S. L., Glass, H. C., Juul, S. E., Chang, T., Mayock, D. E., Durand, D. J., Song, D., Barkovich, A. J., Ballard, R. A., & Wu, Y. W. (2014). Erythropoietin and hypothermia for hypoxic-ischemic encephalopathy. *Pediatr Neurol*, *51*(5), 657-662. https://doi.org/10.1016/j.pediatrneurol.2014.08.010

Rose, A., Wilson, B. A., Manolov, R., & Florschutz, G. (2016). Seeing red: Relearning to read in a case of Balint's Syndrome. *NeuroRehabilitation*, *39*(1), 111-117. https://doi.org/10.3233/nre-161342

Rosen, D., Wilfond, B., & Lantos, J. D. (2013). Obstructive sleep apnea in a 17-year-old with profound cognitive impairment. *Pediatrics*, *131*(3), 581-585. https://doi.org/10.1542/peds.2012-1489

Rosén, H., Karlsson, J. E., & Rosengren, L. (2004). CSF levels of neurofilament is a valuable predictor of long-term outcome after cardiac arrest. *J Neurol Sci*, *221*(1-2), 19-24. https://doi.org/10.1016/j.jns.2004.03.003

Rosenbaum, R. S., Gao, F., Honjo, K., Raybaud, C., Olsen, R. K., Palombo, D. J., Levine, B., & Black, S. E. (2014). Congenital absence of the mammillary bodies: a novel finding in a well-studied case of developmental amnesia. *Neuropsychologia*, *65*, 82-87. https://doi.org/10.1016/j.neuropsychologia.2014.09.047

Rosenegger, D., Roth, S., & Lukowiak, K. (2004). Learning and memory in Lymnaea are negatively altered by acute low-level concentrations of hydrogen sulphide. *J Exp Biol*, *207*(Pt 15), 2621-2630. https://doi.org/10.1242/jeb.01073

Ross, S. J., & Hodges, J. R. (1997). Preservation of famous person knowledge in a patient with severe post anoxic amnesia. *Cortex*, *33*(4), 733-742. https://doi.org/10.1016/s0010-9452(08)70730-5

Rossetti, G. M., d'Avossa, G., Rogan, M., Macdonald, J. H., Oliver, S. J., & Mullins, P. G. (2021). Reversal of neurovascular coupling in the default mode network: Evidence from hypoxia. *J Cereb Blood Flow Metab*, *41*(4), 805-818. https://doi.org/10.1177/0271678x20930827

Ruan, Y. W., Zou, B., Fan, Y., Li, Y., Lin, N., Zeng, Y. S., Gao, T. M., Yao, Z., & Xu, Z. C. (2006). Dendritic plasticity of CA1 pyramidal neurons after transient global ischemia. *Neuroscience*, *140*(1), 191-201. https://doi.org/10.1016/j.neuroscience.2006.01.039

Rubio, J., Caldas, M., Dávila, S., Gasco, M., & Gonzales, G. F. (2006). Effect of three different cultivars of Lepidium meyenii (Maca) on learning and depression in ovariectomized mice. *BMC Complement Altern Med*, *6*, 23. https://doi.org/10.1186/1472-6882-6-23

Rudolf, J., Ghaemi, M., Ghaemi, M., Haupt, W. F., Szelies, B., & Heiss, W. D. (1999). Cerebral glucose metabolism in acute and persistent vegetative state. *J Neurosurg Anesthesiol*, *11*(1), 17-24. https://doi.org/10.1097/00008506-199901000-00004

Ruiz Á, J., Rondón Sepúlveda, M. A., Panqueva Centanaro, O. P., Waich, A., Ruiz, J., Uriza Carrasco, L. F., Ospina García, J. C., Hill, C. M., Restrepo-Gualteros, S. M., Mendoza, L. O., & Hidalgo Martínez, P. (2022). Sleep problems in low income, urban pediatric populations living at different altitudes in Colombia. *Sleep Med*, *100*, 64-70. https://doi.org/10.1016/j.sleep.2022.07.017

Sahu, S., Lata, I., & Gupta, D. (2010). Management of pregnant female with meningioma for

craniotomy. *J Neurosci Rural Pract*, *1*(1), 35-37. https://doi.org/10.4103/0976-3147.63101
Sajkov, D., Marshall, R., Walker, P., Mykytyn, I., McEvoy, R. D., Wale, J., Flavell, H., Thornton, A. T., & Antic, R. (1998). Sleep apnoea related hypoxia is associated with cognitive disturbances in patients with tetraplegia. *Spinal Cord*, *36*(4), 231-239. https://doi.org/10.1038/sj.sc.3100563
Saleh, N. Y., Aboelghar, H. M., Salem, S. S., Ibrahem, R. A., Khalil, F. O., Abdelgawad, A. S., & Mahmoud, A. A. (2021). The severity and atypical presentations of COVID-19 infection in pediatrics. *BMC Pediatr*, *21*(1), 144. https://doi.org/10.1186/s12887-021-02614-2
Saletu, B., & Grünberger, J. (1984). The hypoxia model in human psychopharmacology: neurophysiological and psychometric studies with aniracetam i.v. *Hum Neurobiol*, *3*(3), 171-181.
Saletu, B., Grünberger, J., Anderer, P., Linzmayer, L., & König, P. (1996). On the cerebro-protective effects of caroverine, a calcium-channel blocker and antiglutamatergic drug: double-blind, placebo-controlled, EEG mapping and psychometric studies under hypoxia. *Br J Clin Pharmacol*, *41*(2), 89-99. https://doi.org/10.1111/j.1365-2125.1996.tb00165.x
Saletu, B., Schulz, H., Herrmann, W. M., Anderer, P., Shrotriya, R. C., & Vanbrabant, E. (1994). BMS-181168 for protection of the human brain against hypoxia: double-blind, placebo-controlled EEG mapping studies. *Pharmacopsychiatry*, *27*(5), 189-197. https://doi.org/10.1055/s-2007-1014303
Saletu, B., Semlitsch, H. V., Anderer, P., Resch, F., Presslich, O., & Schuster, P. (1989). Psychophysiological research in psychiatry and neuropsychopharmacology. II. The investigation of antihypoxidotic/nootropic drugs (tenilsetam and co-dergocrine-mesylate) in elderlies with the Viennese Psychophysiological Test-System (VPTS). *Methods Find Exp Clin Pharmacol*, *11*(1), 43-55.
Sallusti, R., Ferraù, S., Lozano Valdes, A., Gonzales, C., Jónsson, M., & Gullo, A. (2001). Altitude decompression sickness. Case presentation. *Minerva Anestesiol*, *67*(10), 737-743.
Salverda, H. H., Cramer, S. J. E., Witlox, R., Gale, T. J., Dargaville, P. A., Pauws, S. C., & Te Pas, A. B. (2022). Comparison of two devices for automated oxygen control in preterm infants: a randomised crossover trial. *Arch Dis Child Fetal Neonatal Ed*, *107*(1), 20-25. https://doi.org/10.1136/archdischild-2020-321387
Salverda, H. H., Dekker, J., Lopriore, E., Dargaville, P. A., Pauws, S. C., & Te Pas, A. B. (2023). Comparison of two automated oxygen controllers in oxygen targeting in preterm infants during admission: an observational study. *Arch Dis Child Fetal Neonatal Ed*, *108*(4), 394-399. https://doi.org/10.1136/archdischild-2022-324819
Samuelson, H., Nekludov, M., & Levander, M. (2008). Neuropsychological outcome following near-drowning in ice water: two adult case studies. *J Int Neuropsychol Soc*, *14*(4), 660-666. https://doi.org/10.1017/s1355617708080855
Sangha, S., Scheibenstock, A., Martens, K., Varshney, N., Cooke, R., & Lukowiak, K. (2005). Impairing forgetting by preventing new learning and memory. *Behav Neurosci*, *119*(3), 787-796. https://doi.org/10.1037/0735-7044.119.3.787
Sanz, F., Restrepo, M. I., Fernández, E., Mortensen, E. M., Aguar, M. C., Cervera, A., Chiner, E., & Blanquer, J. (2011). Hypoxemia adds to the CURB-65 pneumonia severity score in hospitalized patients with mild pneumonia. *Respir Care*, *56*(5), 612-618. https://doi.org/10.4187/respcare.00853
Sara, S. J. (1974). Delayed development of amnestic behavior after hypoxia. *Physiol Behav*, *13*(5), 693-696. https://doi.org/10.1016/0031-9384(74)90242-x
Sarkar, S., Donn, S. M., Bapuraj, J. R., Bhagat, I., & Barks, J. D. (2012). Distribution and severity of hypoxic-ischaemic lesions on brain MRI following therapeutic cooling:

selective head versus whole body cooling. *Arch Dis Child Fetal Neonatal Ed*, *97*(5), F335-339. https://doi.org/10.1136/fetalneonatal-2011-300964
Sawa, Y., Ichikawa, H., Kagisaki, K., Ohata, T., & Matsuda, H. (1998). Interleukin-6 derived from hypoxic myocytes promotes neutrophil-mediated reperfusion injury in myocardium. *J Thorac Cardiovasc Surg*, *116*(3), 511-517. https://doi.org/10.1016/s0022-5223(98)70018-2
Sayre, R. M., Kollias, N., Ley, R. D., & Baqer, A. H. (1994). Changing the risk spectrum of injury and the performance of sunscreen products throughout the day. *Photodermatol Photoimmunol Photomed*, *10*(4), 148-153.
Schaffler, K., Wauschkuhn, C. H., & Häuser, B. (1991). Study to evaluate the encephalotropic potency of a hemodialysate. Controlled study using electro-retinography and visual evoked potentials under hypoxic conditions in human volunteers (preliminary communication). *Arzneimittelforschung*, *41*(7), 699-704.
Schjørring, O. L., Perner, A., Wetterslev, J., Lange, T., Keus, F., Laake, J. H., Okkonen, M., Siegemund, M., Morgan, M., Thormar, K. M., & Rasmussen, B. S. (2019). Handling Oxygenation Targets in the Intensive Care Unit (HOT-ICU)-Protocol for a randomised clinical trial comparing a lower vs a higher oxygenation target in adults with acute hypoxaemic respiratory failure. *Acta Anaesthesiol Scand*, *63*(7), 956-965. https://doi.org/10.1111/aas.13356
Schmidt, A., & Sempsrott, J. (2015). Drowning In The Adult Population: Emergency Department Resuscitation And Treatment. *Emerg Med Pract*, *17*(5), 1-18; quiz 18-19, 22.
Schmidtke, K., & Vollmer, H. (1997). Retrograde amnesia: a study of its relation to anterograde amnesia and semantic memory deficits. *Neuropsychologia*, *35*(4), 505-518. https://doi.org/10.1016/s0028-3932(96)00109-1
Schneider, S., & Strüder, H. K. (2009). Monitoring effects of acute hypoxia on brain cortical activity by using electromagnetic tomography. *Behav Brain Res*, *197*(2), 476-480. https://doi.org/10.1016/j.bbr.2008.10.020
Scholkmann, F., & Denzler, D. (2024). Cerebral Hypoxia During Intermittent Hypoxic-Hyperoxic Training (IHHT): A Case Study Using Cerebral Oximetry Based on Time-Domain Near-Infrared Spectroscopy. *Adv Exp Med Biol*, *1463*, 135-139. https://doi.org/10.1007/978-3-031-67458-7_23
Seibert, P. S., Fee, L., Basom, J., & Zimmerman, C. (2000). Music and the brain: the impact of music on an oboist's fight for recovery. *Brain Inj*, *14*(3), 295-302. https://doi.org/10.1080/026990500120763
Senthilnathan, S., Nallusamy, G., Varadaraj, P., Reddy, K. S. S., & Ravipati, C. (2024). Cortical Laminar Necrosis as a Rare Complication of Streptococcus pneumoniae Meningitis: A Case Report. *Cureus*, *16*(8), e68086. https://doi.org/10.7759/cureus.68086
Seo, J. P., & Jang, S. H. (2013). Recovery of injured cingulum in a patient with brain injury: diffusion tensor tractography study. *NeuroRehabilitation*, *33*(2), 257-261. https://doi.org/10.3233/nre-130953
Shah, A. H., Osten, M., Leventhal, A., Bach, Y., Yoo, D., Mansour, D., Benson, L., Wilson, W. M., & Horlick, E. (2016). Percutaneous Intervention to Treat Platypnea-Orthodeoxia Syndrome: The Toronto Experience. *JACC Cardiovasc Interv*, *9*(18), 1928-1938. https://doi.org/10.1016/j.jcin.2016.07.003
Shah, M. K., Carayannopoulos, A. G., Burke, D. T., & Al-Adawi, S. (2007). A comparison of functional outcomes in hypoxia and traumatic brain injury: a pilot study. *J Neurol Sci*, *260*(1-2), 95-99. https://doi.org/10.1016/j.jns.2007.04.012
Shah, P., Anvekar, A., McMichael, J., & Rao, S. (2015). Outcomes of infants with Apgar score of zero at 10 min: the West Australian experience. *Arch Dis Child Fetal Neonatal Ed*, *100*(6), F492-494. https://doi.org/10.1136/archdischild-2014-307825
Shankaran, S., Laptook, A. R., Pappas, A., McDonald, S. A., Das, A., Tyson, J. E., Poindexter,

B. B., Schibler, K., Bell, E. F., Heyne, R. J., Pedroza, C., Bara, R., Van Meurs, K. P., Huitema, C. M. P., Grisby, C., Devaskar, U., Ehrenkranz, R. A., Harmon, H. M., Chalak, L. F., . . . Higgins, R. D. (2017). Effect of Depth and Duration of Cooling on Death or Disability at Age 18 Months Among Neonates With Hypoxic-Ischemic Encephalopathy: A Randomized Clinical Trial. *Jama*, *318*(1), 57-67. https://doi.org/10.1001/jama.2017.7218

Shankaran, S., McDonald, S. A., Laptook, A. R., Hintz, S. R., Barnes, P. D., Das, A., Pappas, A., & Higgins, R. D. (2015). Neonatal Magnetic Resonance Imaging Pattern of Brain Injury as a Biomarker of Childhood Outcomes following a Trial of Hypothermia for Neonatal Hypoxic-Ischemic Encephalopathy. *J Pediatr*, *167*(5), 987-993.e983. https://doi.org/10.1016/j.jpeds.2015.08.013

Shankaran, S., Pappas, A., McDonald, S. A., Vohr, B. R., Hintz, S. R., Yolton, K., Gustafson, K. E., Leach, T. M., Green, C., Bara, R., Petrie Huitema, C. M., Ehrenkranz, R. A., Tyson, J. E., Das, A., Hammond, J., Peralta-Carcelen, M., Evans, P. W., Heyne, R. J., Wilson-Costello, D. E., . . . Higgins, R. D. (2012). Childhood outcomes after hypothermia for neonatal encephalopathy. *N Engl J Med*, *366*(22), 2085-2092. https://doi.org/10.1056/NEJMoa1112066

Shao, G., Zhang, R., Wang, Z. L., Gao, C. Y., Huo, X., & Lu, G. W. (2006). Hypoxic preconditioning improves spatial cognitive ability in mice. *Neurosignals*, *15*(6), 314-321. https://doi.org/10.1159/000121368

Sharma, N. R., Pokhrel, M., Kc, P., Paudel, S., Lamichhane, P., Rivera Boadla, M. E., & Alvarez, B. (2024). When the Mind Fails: A Mysterious Case of Concurrent Neurosyphilis, Herpes Simplex Virus Encephalitis, and Suspected Autoimmune Encephalitis. *Cureus*, *16*(10), e72415. https://doi.org/10.7759/cureus.72415

Sheldon, K. S., Huey, R. B., Kaspari, M., & Sanders, N. J. (2018). Fifty Years of Mountain Passes: A Perspective on Dan Janzen's Classic Article. *Am Nat*, *191*(5), 553-565. https://doi.org/10.1086/697046

Shi, S. J., Li, H., Liu, M., Liu, Y. M., Zhou, F., Liu, B., Qu, J. X., & Cao, B. (2017). Mortality prediction to hospitalized patients with influenza pneumonia: PO(2) /FiO(2) combined lymphocyte count is the answer. *Clin Respir J*, *11*(3), 352-360. https://doi.org/10.1111/crj.12346

Shibasaki, J., Mukai, T., Tsuda, K., Takeuchi, A., Ioroi, T., Sano, H., Yutaka, N., Takahashi, A., Sobajima, H., Tamura, M., Hosono, S., Nabetani, M., & Iwata, O. (2020). Outcomes related to 10-min Apgar scores of zero in Japan. *Arch Dis Child Fetal Neonatal Ed*, *105*(1), 64-68. https://doi.org/10.1136/archdischild-2019-316793

Shimada, Y., Kobayashi, M., Yoshida, K., Terasaki, K., Fujiwara, S., Kubo, Y., Beppu, T., & Ogasawara, K. (2019). Reduced Hypoxic Tissue and Cognitive Improvement after Revascularization Surgery for Chronic Cerebral Ischemia. *Cerebrovasc Dis*, *47*(1-2), 57-64. https://doi.org/10.1159/000497244

Shimizu, A., & Matsuo, H. (2016). Sound Environments Surrounding Preterm Infants Within an Occupied Closed Incubator. *J Pediatr Nurs*, *31*(2), e149-154. https://doi.org/10.1016/j.pedn.2015.10.011

Shinohara, T., Kojima, H., Nakamura, N., Ogata, A., Betsuyaku, T., Suzuki, A., Maki, Y., & Nagashima, K. (2005). Pathology of pure hippocampal sclerosis in a patient with dementia and Hodgkin's disease: the Ophelia syndrome. *Neuropathology*, *25*(4), 353-360. https://doi.org/10.1111/j.1440-1789.2005.00622.x

Shprecher, D. R., Flanigan, K. M., Smith, A. G., Smith, S. M., Schenkenberg, T., & Steffens, J. (2008). Clinical and diagnostic features of delayed hypoxic leukoencephalopathy. *J Neuropsychiatry Clin Neurosci*, *20*(4), 473-477. https://doi.org/10.1176/jnp.2008.20.4.473

Shu, L., Luo, L., & Zuo, Y. (2023). Attention to pulmonary arteriovenous fistula in a case of transient hypoxemia and cerebral infarction during pregnancy: a case report and

literature review. *BMC Pregnancy Childbirth*, *23*(1), 626. https://doi.org/10.1186/s12884-023-05946-2
Shu, Y., Patel, S. M., Pac-Soo, C., Fidalgo, A. R., Wan, Y., Maze, M., & Ma, D. (2010). Xenon pretreatment attenuates anesthetic-induced apoptosis in the developing brain in comparison with nitrous oxide and hypoxia. *Anesthesiology*, *113*(2), 360-368. https://doi.org/10.1097/ALN.0b013e3181d960d7
Sica, A., Michieletto, P., Pensiero, S., & Barbi, E. (2024). Successful treatment of cortical visual impairment in children using anti-amblyopia treatment despite the absence of amblyopia: a case report. *Ital J Pediatr*, *50*(1), 123. https://doi.org/10.1186/s13052-024-01679-w
Silva, H., Girard, O., Monteiro, J., Gasques, M., Sousa, A., & Nakamura, F. Y. (2025). Competing at Altitude Reduces In-Match Physical Demands of Professional Soccer Players Compared With Sea Level. *Int J Sports Physiol Perform*, *20*(1), 131-141. https://doi.org/10.1123/ijspp.2024-0335
Silva, P. A., Dias, C., Vilarinho, A., Vaz Ferreira, A., Cerejo, A., & Vaz, R. (2022). The Importance of the Temporary Clip Removal Phase on Exposure to Hypoxia: On-Line Measurement of Temporal Lobe Oxygen Levels During Surgery for Middle Cerebral Artery Aneurysms. *Neurosurgery*, *90*(4), 475-484. https://doi.org/10.1227/neu.0000000000001865
Silva-Urra, J. A., Núñez-Espinosa, C. A., Niño-Mendez, O. A., Gaitán-Peñas, H., Altavilla, C., Toro-Salinas, A., Torrella, J. R., Pagès, T., Javierre, C. F., Behn, C., & Viscor, G. (2015). Circadian and Sex Differences After Acute High-Altitude Exposure: Are Early Acclimation Responses Improved by Blue Light? *Wilderness Environ Med*, *26*(4), 459-471. https://doi.org/10.1016/j.wem.2015.06.009
Silver, B. V., Collins, L., & Zidek, K. A. (2003). Risperidone treatment of motor restlessness following anoxic brain injury. *Brain Inj*, *17*(3), 237-244. https://doi.org/10.1080/0269905021000013192
Singh, S. B., Thakur, L., Anand, J. P., Panjwani, U., Yadav, D., & Selvamurthy, W. (2003). Effect of high altitude (HA) on event related brain potentials. *Indian J Physiol Pharmacol*, *47*(1), 52-58.
Sivakumar, K., Mumtaz, Z. A., & Sagar, P. (2022). Application of Vessel Navigator™ fusion imaging software in a complex transcatheter palliation of Tetralogy of Fallot with pulmonary atresia. *Ann Pediatr Cardiol*, *15*(2), 187-191. https://doi.org/10.4103/apc.apc_2_22
Skoch, S. H., Fu, B., Stein, A. L., & Greenstein, S. P. (2021). It Takes a Village: The Importance of Neuropsychological Findings in a Collaborative Approach for a Patient with Congenital Central Hypoventilation Syndrome and Specific Phobia. *Case Rep Psychiatry*, *2021*, 3891481. https://doi.org/10.1155/2021/3891481
Slater, J. P., Guarino, T., Stack, J., Vinod, K., Bustami, R. T., Brown, J. M., 3rd, Rodriguez, A. L., Magovern, C. J., Zaubler, T., Freundlich, K., & Parr, G. V. (2009). Cerebral oxygen desaturation predicts cognitive decline and longer hospital stay after cardiac surgery. *Ann Thorac Surg*, *87*(1), 36-44; discussion 44-35. https://doi.org/10.1016/j.athoracsur.2008.08.070
Smart, J., & Hunter, D. (1984). Alpine travel. Mountain sickness, the unwelcome companion. *Med J Aust*, *141*(12-13), 792-795.
Smith, J., Green, J., Siddiqi, N., Inouye, S. K., Collinson, M., Farrin, A., & Young, J. (2020). Investigation of ward fidelity to a multicomponent delirium prevention intervention during a multicentre, pragmatic, cluster randomised, controlled feasibility trial. *Age Ageing*, *49*(4), 648-655. https://doi.org/10.1093/ageing/afaa042
Speach, D. P., Wong, T. M., Cattarin, J. A., & Livecchi, M. A. (1998). Hypoxic brain injury with motor apraxia following an anaphylactic reaction to hymenoptera venom. *Brain Inj*, *12*(3), 239-244. https://doi.org/10.1080/026990598122719

Spivack, E. (2001). Tetralogy of Fallot: an overview, case report, and discussion of dental implications. *Spec Care Dentist*, *21*(5), 172-175. https://doi.org/10.1111/j.1754-4505.2001.tb00250.x
Squara, P., Denjean, D., Godard, P., Brunet, F., Brusset, A., & Dubois, C. (1994). Enoximone vs nicardipine during the early postoperative course of patients undergoing cardiac surgery. A prospective study of two therapeutic strategies. *Chest*, *106*(1), 52-58. https://doi.org/10.1378/chest.106.1.52
Steen, R. G., Miles, M. A., Helton, K. J., Strawn, S., Wang, W., Xiong, X., & Mulhern, R. K. (2003). Cognitive impairment in children with hemoglobin SS sickle cell disease: relationship to MR imaging findings and hematocrit. *AJNR Am J Neuroradiol*, *24*(3), 382-389.
Steen, R. G., Xiong, X., Mulhern, R. K., Langston, J. W., & Wang, W. C. (1999). Subtle brain abnormalities in children with sickle cell disease: relationship to blood hematocrit. *Ann Neurol*, *45*(3), 279-286. https://doi.org/10.1002/1531-8249(199903)45:3<279::aid-ana2>3.0.co;2-7
Stern, J. M., Sazbon, L., Becker, E., & Costeff, H. (1988). Severe behavioural disturbance in families of patients with prolonged coma. *Brain Inj*, *2*(3), 259-262. https://doi.org/10.3109/02699058809150951
Steverlynck, L., Baert, N., Buylaert, W., & De Paepe, P. (2017). Combined acute inhalation of hydrofluoric acid and nitric acid: a case report and literature review. *Acta Clin Belg*, *72*(4), 278-288. https://doi.org/10.1080/17843286.2016.1229840
Stoller, K. P. (2007). Hyperbaric oxygen and carbon monoxide poisoning: a critical review. *Neurol Res*, *29*(2), 146-155. https://doi.org/10.1179/016164107x181770
Strata, F., Coq, J. O., Byl, N., & Merzenich, M. M. (2004). Effects of sensorimotor restriction and anoxia on gait and motor cortex organization: implications for a rodent model of cerebral palsy. *Neuroscience*, *129*(1), 141-156. https://doi.org/10.1016/j.neuroscience.2004.07.024
Strens, L. H., Mazibrada, G., Duncan, J. S., & Greenwood, R. (2004). Misdiagnosing the vegetative state after severe brain injury: the influence of medication. *Brain Inj*, *18*(2), 213-218. https://doi.org/10.1080/0269905031000149533
Stuss, D. T., Peterkin, I., Guzman, D. A., Guzman, C., & Troyer, A. K. (1997). Chronic obstructive pulmonary disease: effects of hypoxia on neurological and neuropsychological measures. *J Clin Exp Neuropsychol*, *19*(4), 515-524. https://doi.org/10.1080/01688639708403741
Su, R., Han, C., Chen, G., Li, H., Liu, W., Wang, C., Zhang, W., Zhang, Y., Zhang, D., & Ma, H. (2024). Low- and moderate-intensity aerobic exercise improves the physiological acclimatization of lowlanders on the Tibetan plateau. *Eur J Sport Sci*, *24*(6), 834-845. https://doi.org/10.1002/ejsc.12110
Su, R., Wang, C., Liu, W., Han, C., Fan, J., Ma, H., Li, H., & Zhang, D. (2022). Intensity-dependent acute aerobic exercise: Effect on reactive control of attentional functions in acclimatized lowlanders at high altitude. *Physiol Behav*, *250*, 113785. https://doi.org/10.1016/j.physbeh.2022.113785
Sugimoto, R., Kenzaka, T., Fujikawa, M., Kawasaki, S., & Nishisaki, H. (2020). Humidifier Use and Prone Positioning in a Patient with Severe COVID-19 Pneumonia and Endotracheal Tube Impaction Due to Highly Viscous Sputum. *Cureus*, *12*(6), e8626. https://doi.org/10.7759/cureus.8626
Sugimura, M., Hanamoto, H., Boku, A., Morimoto, Y., Taki, K., Kudo, C., & Niwa, H. (2010). Influence of acute hypoxia combined with nitrous oxide on cardiovascular variability in conscious hypertensive rats. *Auton Neurosci*, *156*(1-2), 73-81. https://doi.org/10.1016/j.autneu.2010.04.008
Sugimura, M., Hirose, Y., Hanamoto, H., Okada, K., Boku, A., Morimoto, Y., Taki, K., & Niwa, H.

(2008). Influence of acute progressive hypoxia on cardiovascular variability in conscious spontaneously hypertensive rats. *Auton Neurosci*, *141*(1-2), 94-103. https://doi.org/10.1016/j.autneu.2008.05.008
Sugrue, P. A., Hurley, M. C., Bendok, B. R., Surdell, D. L., Gottardi-Littell, N., Futterer, S. F., Muro, K., & Batjer, H. H. (2009). High-grade dural arteriovenous fistula simulating a bilateral thalamic neoplasm. *Clin Neurol Neurosurg*, *111*(7), 629-632. https://doi.org/10.1016/j.clineuro.2009.05.004
Sulu, B., Yildiz, B. D., Buyukuysal, C., Demir, E., & Gunerhan, Y. (2012). Comparison of meperidine versus hyoscine during colonoscopy in the elderly: a prospective randomized study. *J Laparoendosc Adv Surg Tech A*, *22*(7), 631-634. https://doi.org/10.1089/lap.2012.0117
Sun, S., Loprinzi, P. D., Guan, H., Zou, L., Kong, Z., Hu, Y., Shi, Q., & Nie, J. (2019). The Effects of High-Intensity Interval Exercise and Hypoxia on Cognition in Sedentary Young Adults. *Medicina (Kaunas)*, *55*(2). https://doi.org/10.3390/medicina55020043
Sutin, J., Vyas, R., Feldman, H. A., Ferradal, S., Hsiao, C. H., Zampolli, L., Pierce, L. J., Nelson, C. A., Morton, S. U., Hay, S., El-Dib, M., Soul, J. S., Lin, P. Y., & Grant, P. E. (2023). Association of cerebral metabolic rate following therapeutic hypothermia with 18-month neurodevelopmental outcomes after neonatal hypoxic ischemic encephalopathy. *EBioMedicine*, *94*, 104673. https://doi.org/10.1016/j.ebiom.2023.104673
Szlabowicz, J. W., & Stewart, J. T. (1990). Amitriptyline treatment of agitation associated with anoxic encephalopathy. *Arch Phys Med Rehabil*, *71*(8), 612-613.
Takuma, T., Okada, K., Uchida, Y., Yamagata, A., & Sawae, Y. (2002). Invasive pulmonary aspergillosis resulting in respiratory failure during neutrophil recovery from postchemotherapy neutropenia in three patients with acute leukaemia. *J Intern Med*, *252*(2), 173-177. https://doi.org/10.1046/j.1365-2796.2002.01012.x
Tamang, A. M., Kalra, B., & Parkash, R. (2017). Cold and desiccation stress induced changes in the accumulation and utilization of proline and trehalose in seasonal populations of Drosophila immigrans. *Comp Biochem Physiol A Mol Integr Physiol*, *203*, 304-313. https://doi.org/10.1016/j.cbpa.2016.10.011
Tan, L., Furian, M., Li, T., & Tang, X. (2022). Effect of acetazolamide on obstructive sleep apnoea in highlanders: protocol for a randomised, placebo-controlled, double-blinded crossover trial. *BMJ Open*, *12*(3), e057113. https://doi.org/10.1136/bmjopen-2021-057113
Tan, L., Li, T., Zhang, Y., He, D., Luo, L., Lei, F., Ren, R., He, J., Bloch, K. E., & Tang, X. (2021). Effect of One Night of Nocturnal Oxygen Supplementation on Highland Patients With OSA: A Randomized, Crossover Trial. *Chest*, *160*(2), 690-700. https://doi.org/10.1016/j.chest.2021.02.046
Tang, A. C., & Nakazawa, M. (2005). Neonatal novelty exposure ameliorates anoxia-induced hyperactivity in the open field. *Behav Brain Res*, *163*(1), 1-9. https://doi.org/10.1016/j.bbr.2005.03.025
Targa, A. D. S., Benítez, I. D., Dakterzada, F., Carnes, A., Pujol, M., Jorge, C., Minguez, O., Dalmases, M., Sánchez-de-la-Torre, M., Barbé, F., & Piñol-Ripoll, G. (2021). Sleep profile predicts the cognitive decline of mild-moderate Alzheimer's disease patients. *Sleep*, *44*(10). https://doi.org/10.1093/sleep/zsab117
Tas, M., Kurtulus, M., Gulnerman, E. F. K., Turkyilmaz, C., Percin, F., Ergenekon, E., & Koc, E. (2022). Pitt-Hopkins syndrome accompanying hypoxic ischemic encephalopathy in a newborn. *Int J Dev Neurosci*, *82*(5), 458-462. https://doi.org/10.1002/jdn.10203
Tazopoulou, E., Miljkovitch, R., Truelle, J. L., Schnitzler, A., Onillon, M., Zucco, T., Hawthorne, G., & Montreuil, M. (2016). Rehabilitation following cerebral anoxia: An assessment of 27 patients. *Brain Inj*, *30*(1), 95-103. https://doi.org/10.3109/02699052.2015.1113563
Ten, V. S., Bradley-Moore, M., Gingrich, J. A., Stark, R. I., & Pinsky, D. J. (2003). Brain injury

and neurofunctional deficit in neonatal mice with hypoxic-ischemic encephalopathy. *Behav Brain Res*, *145*(1-2), 209-219. https://doi.org/10.1016/s0166-4328(03)00146-3
Tharmapoopathy, P., Chisholm, P., Barlas, A., Varsami, M., Gupta, N., Ekitzidou, G., Ponnusamy, V., Kappelou, O., Evanson, J., Rosser, G., & Shah, D. K. (2020). In clinical practice, cerebral MRI in newborns is highly predictive of neurodevelopmental outcome after therapeutic hypothermia. *Eur J Paediatr Neurol*, *25*, 127-133. https://doi.org/10.1016/j.ejpn.2019.12.018
Thiebauld, C., Van Mullem, J., Lintermans, J., & Sprumont, P. (1983). Testing in a hypobaric chamber drugs claimed to improve impaired brain function. *Lancet*, *2*(8343), 225-226. https://doi.org/10.1016/s0140-6736(83)90212-x
Thomas, R. J., Rosen, B. R., Stern, C. E., Weiss, J. W., & Kwong, K. K. (2005). Functional imaging of working memory in obstructive sleep-disordered breathing. *J Appl Physiol (1985)*, *98*(6), 2226-2234. https://doi.org/10.1152/japplphysiol.01225.2004
Tiainen, M., Poutiainen, E., Kovala, T., Takkunen, O., Häppölä, O., & Roine, R. O. (2007). Cognitive and neurophysiological outcome of cardiac arrest survivors treated with therapeutic hypothermia. *Stroke*, *38*(8), 2303-2308. https://doi.org/10.1161/strokeaha.107.483867
Tirabassi, J. N., Olewinski, L., & Khodaee, M. (2018). Variation of Traditional Biomarkers of Liver Injury After an Ultramarathon at Altitude. *Sports Health*, *10*(4), 361-365. https://doi.org/10.1177/1941738118764870
Tonsgard, J. H., Harwicke, N., & Levine, S. C. (1987). Kluver-Bucy syndrome in children. *Pediatr Neurol*, *3*(3), 162-165. https://doi.org/10.1016/0887-8994(87)90084-1
Tran, K., & Wu, J. (2019). Case report: neuroimaging analysis of pediatric ADHD-related symptoms secondary to hypoxic brain injury. *Brain Inj*, *33*(10), 1402-1407. https://doi.org/10.1080/02699052.2019.1641744
Tripp, L. D., Chelette, T., Savul, S., & Widman, R. A. (1998). Female exposure to high G: effects of simulated combat sorties on cerebral and arterial O2 saturation. *Aviat Space Environ Med*, *69*(9), 869-874.
Tsukiura, T., Fujii, T., Fukatsu, R., Otsuki, T., Okuda, J., Umetsu, A., Suzuki, K., Tabuchi, M., Yanagawa, I., Nagasaka, T., Kawashima, R., Fukuda, H., Takahashi, S., & Yamadori, A. (2002). Neural basis of the retrieval of people's names: evidence from brain-damaged patients and fMRI. *J Cogn Neurosci*, *14*(6), 922-937. https://doi.org/10.1162/089892902760191144
Tubek, S., Niewinski, P., Reczuch, K., Janczak, D., Rucinski, A., Paleczny, B., Engelman, Z. J., Banasiak, W., Paton, J. F., & Ponikowski, P. (2016). Effects of selective carotid body stimulation with adenosine in conscious humans. *J Physiol*, *594*(21), 6225-6240. https://doi.org/10.1113/jp272109
Tukacs, V., Mittli, D., Györffy, B. A., Hunyady-Gulyás, É., Hlatky, D., Tóth, V., Ravasz, L., Medzihradszky, F. K., Nyitrai, G., Czurkó, A., Juhász, G., Kardos, J., & Kékesi, K. A. (2020). Chronic stepwise cerebral hypoperfusion differentially induces synaptic proteome changes in the frontal cortex, occipital cortex, and hippocampus in rats. *Sci Rep*, *10*(1), 15999. https://doi.org/10.1038/s41598-020-72868-w
Tvaryanas, A. P. (2004). Visual scan patterns during simulated control of an uninhabited aerial vehicle (UAV). *Aviat Space Environ Med*, *75*(6), 531-538.
Tvedt, B., Skyberg, K., Aaserud, O., Hobbesland, A., & Mathiesen, T. (1991). Brain damage caused by hydrogen sulfide: a follow-up study of six patients. *Am J Ind Med*, *20*(1), 91-101. https://doi.org/10.1002/ajim.4700200109
Tzouvelekis, A., Aidinis, V., Harokopos, V., Karameris, A., Zacharis, G., Mikroulis, D., Konstantinou, F., Steiropoulos, P., Sotiriou, I., Froudarakis, M., Pneumatikos, I., Tringidou, R., & Bouros, D. (2009). Down-regulation of the inhibitor of growth family member 4 (ING4) in different forms of pulmonary fibrosis. *Respir Res*, *10*(1), 14.

https://doi.org/10.1186/1465-9921-10-14
Ueno-Pardi, L. M., Souza-Duran, F. L., Matheus, L., Rodrigues, A. G., Barbosa, E. R. F., Cunha, P. J., Carneiro, C. G., Costa, N. A., Ono, C. R., Buchpiguel, C. A., Negrão, C. E., Lorenzi-Filho, G., & Busatto-Filho, G. (2022). Effects of exercise training on brain metabolism and cognitive functioning in sleep apnea. *Sci Rep*, *12*(1), 9453. https://doi.org/10.1038/s41598-022-13115-2
Umeda, Y., Matsuda, H., Sadamori, H., Shinoura, S., Yoshida, R., Sato, D., Utsumi, M., Yagi, T., & Fujiwara, T. (2011). Leukoencephalopathy syndrome after living-donor liver transplantation. *Exp Clin Transplant*, *9*(2), 139-144.
Urushida, Y., Kikuchi, Y., Shimizu, C., Amari, M., Kawarabayashi, T., Nakamura, T., Ikeda, Y., Takatama, M., & Shoji, M. (2021). Improved Neuroimaging Findings and Cognitive Function in a Case of High-altitude Cerebral Edema. *Intern Med*, *60*(8), 1299-1302. https://doi.org/10.2169/internalmedicine.5747-20
Úsuga, M. J., Jaramillo, G. A., Palacio, V., Correa, S. A., & Suárez-Escudero, J. C. (2021). Velamentous cord insertion, ischemic-hypoxic encephalopathy, and neurological rehabilitation: A case report. *Biomedica*, *41*(1), 8-16. https://doi.org/10.7705/biomedica.5436
Usui, C., Inoue, Y., Kimura, M., Kirino, E., Nagaoka, S., Abe, M., Nagata, T., & Arai, H. (2004). Irreversible subcortical dementia following high altitude illness. *High Alt Med Biol*, *5*(1), 77-81. https://doi.org/10.1089/152702904322963717
Vacchiano, C., Moore, J., Rice, G. M., & Crawley, G. (2008). Fexofenadine effects on cognitive performance in aviators at ground level and simulated altitude. *Aviat Space Environ Med*, *79*(8), 754-760. https://doi.org/10.3357/asem.2212.2008
Vakili, K., Pillay, S. S., Lafer, B., Fava, M., Renshaw, P. F., Bonello-Cintron, C. M., & Yurgelun-Todd, D. A. (2000). Hippocampal volume in primary unipolar major depression: a magnetic resonance imaging study. *Biol Psychiatry*, *47*(12), 1087-1090. https://doi.org/10.1016/s0006-3223(99)00296-6
Valk, P. J., Simons, R., Jetten, A. M., Valiente, R., & Labeaga, L. (2016). Cognitive Performance Effects of Bilastine 20 mg During 6 Hours at 8000 ft Cabin Altitude. *Aerosp Med Hum Perform*, *87*(7), 622-627. https://doi.org/10.3357/amhp.4522.2016
Valk, P. J., Simons, R. M., Struyvenberg, P. A., Kruit, H., & van Berge Henegouwen, M. T. (1997). Effects of a single dose of loratadine on flying ability under conditions of simulated cabin pressure. *Am J Rhinol*, *11*(1), 27-33. https://doi.org/10.2500/105065897781446838
Van Leuven, W., Van Dam, D., Moens, L., De Deyn, P. P., & Dewilde, S. (2013). A behavioural study of neuroglobin-overexpressing mice under normoxic and hypoxic conditions. *Biochim Biophys Acta*, *1834*(9), 1764-1771. https://doi.org/10.1016/j.bbapap.2013.04.015
van Zomeren, A. H., ten Duis, H. J., Minderhoud, J. M., & Sipma, M. (1998). Lightning stroke and neuropsychological impairment: cases and questions. *J Neurol Neurosurg Psychiatry*, *64*(6), 763-769. https://doi.org/10.1136/jnnp.64.6.763
Vangeison, G., Carr, D., Federoff, H. J., & Rempe, D. A. (2008). The good, the bad, and the cell type-specific roles of hypoxia inducible factor-1 alpha in neurons and astrocytes. *J Neurosci*, *28*(8), 1988-1993. https://doi.org/10.1523/jneurosci.5323-07.2008
Vaughan, R. W., & Wise, L. (1976). Intraoperative arterial oxygenation in obese patients. *Ann Surg*, *184*(1), 35-42. https://doi.org/10.1097/00000658-197607000-00006
Veneman, T. F., van Dijk, G. W., Boereboom, E., Joore, H., & Savelkoul, T. J. (1998). Prediction of outcome after resuscitation in a case of electrocution. *Intensive Care Med*, *24*(3), 255-257. https://doi.org/10.1007/s001340050560
Verberk, W., Calosi, P., Spicer, J. I., Kehl, S., & Bilton, D. T. (2018). Does plasticity in thermal tolerance trade off with inherent tolerance? The influence of setal tracheal gills on

thermal tolerance and its plasticity in a group of European diving beetles. *J Insect Physiol*, *106*(Pt 3), 163-171. https://doi.org/10.1016/j.jinsphys.2017.12.005
Verfaellie, M., Koseff, P., & Alexander, M. P. (2000). Acquisition of novel semantic information in amnesia: effects of lesion location. *Neuropsychologia*, *38*(4), 484-492. https://doi.org/10.1016/s0028-3932(99)00089-5
Vinetti, G., Micarelli, A., Falla, M., Randi, A., Dal Cappello, T., Gatterer, H., Brugger, H., Strapazzon, G., & Rauch, S. (2023). Surgical masks and filtering facepiece class 2 respirators (FFP2) have no major physiological effects at rest and during moderate exercise at 3000-m altitude: a randomised controlled trial. *J Travel Med*, *30*(5). https://doi.org/10.1093/jtm/taad031
Virto, M., Bustamante, M., Ruiz de Gordoa, J. C., Amores, G., Fernández-Caballero, P. N., Mandaluniz, N., Arranz, J., Nájera, A. I., Albisu, M., Pérez-Elortondo, F. J., Barron, L. J., & de Renobales, M. (2012). Interannual and geographical reproducibility of the nutritional quality of milk fat from commercial grazing flocks. *J Dairy Res*, *79*(4), 485-494. https://doi.org/10.1017/s0022029912000490
Visser, B. J., Korevaar, D. A., & van der Zee, J. (2012). A 24-year-old Ethiopian farmer with burning feet. *Am J Trop Med Hyg*, *87*(4), 583. https://doi.org/10.4269/ajtmh.2012.12-0405
Vo, H. H., Keegan, D., Sveen, W. N., Wilfond, B. S., Campelia, G., & Henderson, C. M. (2024). Candidacy Decisions for Long-term Ventilation. *Pediatrics*, *154*(6). https://doi.org/10.1542/peds.2024-066985
Vögele, A., van Veelen, M. J., Dal Cappello, T., Falla, M., Nicoletto, G., Dejaco, A., Palma, M., Hüfner, K., Brugger, H., & Strapazzon, G. (2021). Effect of Acute Exposure to Altitude on the Quality of Chest Compression-Only Cardiopulmonary Resuscitation in Helicopter Emergency Medical Services Personnel: A Randomized, Controlled, Single-Blind Crossover Trial. *J Am Heart Assoc*, *10*(23), e021090. https://doi.org/10.1161/jaha.121.021090
Vohr, B. R., Stephens, B. E., McDonald, S. A., Ehrenkranz, R. A., Laptook, A. R., Pappas, A., Hintz, S. R., Shankaran, S., Higgins, R. D., & Das, A. (2013). Cerebral palsy and growth failure at 6 to 7 years. *Pediatrics*, *132*(4), e905-914. https://doi.org/10.1542/peds.2012-3915
Volpe, B. T., & Hirst, W. (1983). The characterization of an amnesic syndrome following hypoxic ischemic injury. *Arch Neurol*, *40*(7), 436-440. https://doi.org/10.1001/archneur.1983.04050070066017
Vos, P. J., Folgering, H. T., & van Herwaarden, C. L. (1995). Visual attention in patients with chronic obstructive pulmonary disease. *Biol Psychol*, *41*(3), 295-305. https://doi.org/10.1016/0301-0511(95)05140-6
Vuilleumier, P., Staub, F., & Assal, G. (1997). Sniffing behaviour, or recognizing a lily by smell, but not recognizing a sock on sight. *Cortex*, *33*(3), 571-577. https://doi.org/10.1016/s0010-9452(08)70238-7
Wagner, A. S., Baumann, S. M., Semmlack, S., Frei, A. I., Rüegg, S., Hunziker, S., Marsch, S., & Sutter, R. (2023). Comparing Patients With Isolated Seizures and Status Epilepticus in Intensive Care Units: An Observational Cohort Study. *Neurology*, *100*(17), e1763-e1775. https://doi.org/10.1212/wnl.0000000000206838
Wagner, S., Quente, J., Staedtler, S., Koch, K., Richter-Schmidinger, T., Kornhuber, J., Ihmsen, H., & Schuettler, J. (2018). A high risk of sleep apnea is associated with less postoperative cognitive dysfunction after intravenous anesthesia: results of an observational pilot study. *BMC Anesthesiol*, *18*(1), 139. https://doi.org/10.1186/s12871-018-0602-9
Wagner, S., Sutter, L., Wagenblast, F., Walther, A., & Schiff, J. H. (2021). Short term cognitive function after sevoflurane anesthesia in patients suspect to obstructive sleep apnea

syndrome: an observational study. *BMC Anesthesiol*, *21*(1), 150. https://doi.org/10.1186/s12871-021-01363-0
Walker, M., Warburton, K. M., Rencic, J., & Parsons, A. S. (2019). Lessons in clinical reasoning - pitfalls, myths, and pearls: a case of chest pain and shortness of breath. *Diagnosis (Berl)*, *6*(4), 387-392. https://doi.org/10.1515/dx-2019-0030
Walrath, B., Smith, J. E., Raghunandan, A., Boni, B., & Latham, E. (2013). Differential diagnosis considerations of sickness after rapid pressure changes at altitude. *Aviat Space Environ Med*, *84*(12), 1291-1294. https://doi.org/10.3357/asem.3621.2013
Wang, J., Ke, T., Zhang, X., Chen, Y., Liu, M., Chen, J., & Luo, W. (2013). Effects of acetazolamide on cognitive performance during high-altitude exposure. *Neurotoxicol Teratol*, *35*, 28-33. https://doi.org/10.1016/j.ntt.2012.12.003
Wang, J. S., Chen, W. L., & Weng, T. P. (2011). Hypoxic exercise training reduces senescent T-lymphocyte subsets in blood. *Brain Behav Immun*, *25*(2), 270-278. https://doi.org/10.1016/j.bbi.2010.09.018
Wang, J. S., & Weng, T. P. (2011). Hypoxic exercise training promotes antitumour cytotoxicity of natural killer cells in young men. *Clin Sci (Lond)*, *121*(8), 343-353. https://doi.org/10.1042/cs20110032
Wang, T., Wang, Y., Liang, Y., & Lu, G. (2015). Infantile Hepatic Hemangioendothelioma Associated With Congestive Heart Failure: Two Case Reports With Different Outcomes. *Medicine (Baltimore)*, *94*(52), e2344. https://doi.org/10.1097/md.0000000000002344
Wang, W. C., Yang, H. C., & Chen, Y. J. (2015). Acute multiple focal neuropathies and delayed postanoxic encephalopathy after alcohol intoxication. *Neuropsychiatr Dis Treat*, *11*, 1781-1784. https://doi.org/10.2147/ndt.S87731
Wang, Y., Meng, R., Song, H., Liu, G., Hua, Y., Cui, D., Zheng, L., Feng, W., Liebeskind, D. S., Fisher, M., & Ji, X. (2017). Remote Ischemic Conditioning May Improve Outcomes of Patients With Cerebral Small-Vessel Disease. *Stroke*, *48*(11), 3064-3072. https://doi.org/10.1161/strokeaha.117.017691
Watcha, M. F., Garner, F. T., White, P. F., & Lusk, R. (1994). Laryngeal mask airway vs face mask and Guedel airway during pediatric myringotomy. *Arch Otolaryngol Head Neck Surg*, *120*(8), 877-880. https://doi.org/10.1001/archotol.1994.01880320077017
Weale, R. A. (1982). The age variation of "senile' cataract in various parts of the world. *Br J Ophthalmol*, *66*(1), 31-34. https://doi.org/10.1136/bjo.66.1.31
Weinberger, M., & Abu-Hasan, M. (2009). Perceptions and pathophysiology of dyspnea and exercise intolerance. *Pediatr Clin North Am*, *56*(1), 33-48, ix. https://doi.org/10.1016/j.pcl.2008.10.015
White, A. J. (1984). Cognitive impairment of acute mountain sickness and acetazolamide. *Aviat Space Environ Med*, *55*(7), 598-603.
Wickens, C. D., Liang, C. C., Prevett, T., & Olmos, O. (1996). Electronic maps for terminal area navigation: effects of frame of reference and dimensionality. *Int J Aviat Psychol*, *6*(3), 241-271. https://doi.org/10.1207/s15327108ijap0603_3
Wickens, C. D., & Seidler, K. S. (1997). Information access in a dual-task context: testing a model of optimal strategy selection. *J Exp Psychol Appl*, *3*(3), 196-215. https://doi.org/10.1037//1076-898x.3.3.196
Williams, J. A., Barreiro, C. J., Nwakanma, L. U., Lange, M. S., Kratz, L. E., Blue, M. E., Berrong, J., Patel, N. D., Gott, V. L., Troncoso, J. C., Johnston, M. V., & Baumgartner, W. A. (2006). Valproic acid prevents brain injury in a canine model of hypothermic circulatory arrest: a promising new approach to neuroprotection during cardiac surgery. *Ann Thorac Surg*, *81*(6), 2235-2241; discussion 2241-2232. https://doi.org/10.1016/j.athoracsur.2005.12.060
Williams, T. B., Badariotti, J. I., Corbett, J., Miller-Dicks, M., Neupert, E., McMorris, T., Ando, S., Parker, M. O., Thelwell, R. C., Causer, A. J., Young, J. S., Mayes, H. S., White, D. K., de

Carvalho, F. A., Tipton, M. J., & Costello, J. T. (2024). The effects of sleep deprivation, acute hypoxia, and exercise on cognitive performance: A multi-experiment combined stressors study. *Physiol Behav*, *274*, 114409. https://doi.org/10.1016/j.physbeh.2023.114409

Wilson, D. K., Kaplan, R. M., Timms, R. M., & Dawson, A. (1985). Acute effects of oxygen treatment upon information processing in hypoxemic COPD patients. *Chest*, *88*(2), 239-243. https://doi.org/10.1378/chest.88.2.239

Wilson, F. C., Harpur, J., Watson, T., & Morrow, J. I. (2003). Adult survivors of severe cerebral hypoxia--case series survey and comparative analysis. *NeuroRehabilitation*, *18*(4), 291-298.

Wilson, J. E., Duggan, M. C., Chandrasekhar, R., Brummel, N. E., Dittus, R. S., Ely, E. W., Patel, M. B., & Jackson, J. C. (2019). Deficits in Self-Reported Initiation Are AssociatedWith Subsequent Disability in ICU Survivors. *Psychosomatics*, *60*(4), 376-384. https://doi.org/10.1016/j.psym.2018.09.004

Wong, K. K., Grunstein, R. R., Bartlett, D. J., & Gordon, E. (2006). Brain function in obstructive sleep apnea: results from the Brain Resource International Database. *J Integr Neurosci*, *5*(1), 111-121. https://doi.org/10.1142/s0219635206001033

Wright, K. L., Kirwan, C. B., Gale, S. D., Levan, A. J., & Hopkins, R. O. (2017). Long-term cognitive and neuroanatomical stability in patients with anoxic amnesia: A Case Report. *Brain Inj*, *31*(5), 709-716. https://doi.org/10.1080/02699052.2017.1285051

Wright, M., & Nolan, T. (1994). Impact of cyanotic heart disease on school performance. *Arch Dis Child*, *71*(1), 64-70. https://doi.org/10.1136/adc.71.1.64

Wu, J., Gu, M., Chen, S., Chen, W., Ni, K., Xu, H., & Li, X. (2017). Factors related to pediatric obstructive sleep apnea-hypopnea syndrome in children with attention deficit hyperactivity disorder in different age groups. *Medicine (Baltimore)*, *96*(42), e8281. https://doi.org/10.1097/md.0000000000008281

Wu, Y. W., Comstock, B. A., Gonzalez, F. F., Mayock, D. E., Goodman, A. M., Maitre, N. L., Chang, T., Van Meurs, K. P., Lampland, A. L., Bendel-Stenzel, E., Mathur, A. M., Wu, T. W., Riley, D., Mietzsch, U., Chalak, L., Flibotte, J., Weitkamp, J. H., Ahmad, K. A., Yanowitz, T. D., . . . Juul, S. E. (2022). Trial of Erythropoietin for Hypoxic-Ischemic Encephalopathy in Newborns. *N Engl J Med*, *387*(2), 148-159. https://doi.org/10.1056/NEJMoa2119660

Wu, Y. W., Monsell, S. E., Glass, H. C., Wisnowski, J. L., Mathur, A. M., McKinstry, R. C., Bluml, S., Gonzalez, F. F., Comstock, B. A., Heagerty, P. J., & Juul, S. E. (2023). How well does neonatal neuroimaging correlate with neurodevelopmental outcomes in infants with hypoxic-ischemic encephalopathy? *Pediatr Res*, *94*(3), 1018-1025. https://doi.org/10.1038/s41390-023-02510-8

Xapsos, M. A., Summers, G. P., Shapiro, P., & Burke, E. A. (1996). New techniques for predicting solar proton fluences for radiation effects applications. *IEEE Trans Nucl Sci*, *43*(6), 2772-2777. https://doi.org/10.1109/23.556865

Xi, L., Smith, C. A., Saupe, K. W., & Dempsey, J. A. (1993). Effects of memory from vagal feedback on short-term potentiation of ventilation in conscious dogs. *J Physiol*, *462*, 547-561. https://doi.org/10.1113/jphysiol.1993.sp019568

Xie, L., Li, Y., Jiang, X., Zhao, J., & Xiao, T. (2019). A 10-year-old boy with dyspnea and hypoxia: abernathy malformation masquerading as pulmonary arteriovenous fistula. *BMC Pediatr*, *19*(1), 55. https://doi.org/10.1186/s12887-019-1422-x

Xu, Z. Y., Wang, X., Si, Y. Y., Wu, J. C., Zuo, Y. X., Xue, F. S., & Liu, J. (2008). Intravenous remifentanil and propofol for gastroscopy. *J Clin Anesth*, *20*(5), 352-355. https://doi.org/10.1016/j.jclinane.2008.03.006

Yamanashi, Y., Mori, M., Terajima, K., Tsueshita, T., Horinouchi, H., Sakai, H., & Sakamoto, A. (2009). A transient inflammatory reaction in the lung after experimental hemorrhagic

shock and resuscitation with a hemoglobin-vesicles solution compared with rat RBC transfusion. *Asaio j*, *55*(5), 478-483. https://doi.org/10.1097/MAT.0b013e3181b17f34
Yamazaki, H., Okazaki, M., Takeda, R., & Haji, A. (2002). Hypercapnic and hypoxic ventilatory responses in long-term streptozotocin-diabetic rats during conscious and pentobarbital-induced anesthetic states. *Life Sci*, *72*(1), 79-89. https://doi.org/10.1016/s0024-3205(02)02201-4
Yang, J., Shanahan, K. J., Shriver, L. P., & Luciano, M. G. (2016). Exercise-induced changes of cerebrospinal fluid vascular endothelial growth factor in adult chronic hydrocephalus patients. *J Clin Neurosci*, *24*, 52-56. https://doi.org/10.1016/j.jocn.2015.08.019
Yang, Y., Wang, L., Han, J., Tang, X., Ma, M., Wang, K., Zhang, X., Ren, Q., Chen, Q., & Qiu, Q. (2015). Comparative transcriptomic analysis revealed adaptation mechanism of Phrynocephalus erythrurus, the highest altitude Lizard living in the Qinghai-Tibet Plateau. *BMC Evol Biol*, *15*, 101. https://doi.org/10.1186/s12862-015-0371-8
Yates, K. B., Tonnerre, P., Martin, G. E., Gerdemann, U., Al Abosy, R., Comstock, D. E., Weiss, S. A., Wolski, D., Tully, D. C., Chung, R. T., Allen, T. M., Kim, A. Y., Fidler, S., Fox, J., Frater, J., Lauer, G. M., Haining, W. N., & Sen, D. R. (2021). Epigenetic scars of CD8(+) T cell exhaustion persist after cure of chronic infection in humans. *Nat Immunol*, *22*(8), 1020-1029. https://doi.org/10.1038/s41590-021-00979-1
Yavaşcaoğlu, B., Cebelli, V., Kelebek, N., Uçkunkaya, N., & Kutlay, O. (2002). Comparison of different priming techniques on the onset time and intubating conditions of rocuronium. *Eur J Anaesthesiol*, *19*(7), 517-521. https://doi.org/10.1017/s0265021502000844
Yee, T., Gronner, A., & Knight, R. T. (1994). CT findings of hypoxic basal ganglia damage. *South Med J*, *87*(6), 624-626. https://doi.org/10.1097/00007611-199406000-00009
Yoneda, I., Tomoda, M., Tokumaru, O., Sato, T., & Watanabe, Y. (2000). Time of useful consciousness determination in aircrew members with reference to prior altitude chamber experience and age. *Aviat Space Environ Med*, *71*(1), 72-76.
Yoneda, I., & Watanabe, Y. (1997). Comparisons of altitude tolerance and hypoxia symptoms between nonsmokers and habitual smokers. *Aviat Space Environ Med*, *68*(9), 807-811.
Yonelinas, A. P., Kroll, N. E., Quamme, J. R., Lazzara, M. M., Sauvé, M. J., Widaman, K. F., & Knight, R. T. (2002). Effects of extensive temporal lobe damage or mild hypoxia on recollection and familiarity. *Nat Neurosci*, *5*(11), 1236-1241. https://doi.org/10.1038/nn961
Yoshida, S., Nabeshima, T., Kinbara, K., & Kameyama, T. (1992). Effects of NIK-247 on CO-induced impairment of passive avoidance in mice. *Eur J Pharmacol*, *214*(2-3), 247-252. https://doi.org/10.1016/0014-2999(92)90125-n
Yoshii, S., Suzuki, S., Osawa, H., Hosaka, S., Honda, Y., Abraham, S. J., Tada, Y., Sugiyama, H., Tan, T., Kadono, T., Hoshiai, M., & Komai, T. (2002). Accessory hepatic vein complicating extra-cardiac total cavopulmonary connection. *Ann Thorac Cardiovasc Surg*, *8*(2), 112-114.
Zabel, T. A., Slomine, B., Brady, K., & Christensen, J. (2005). Neuropsychological profile following suicide attempt by hanging: two adolescent case reports. *Child Neuropsychol*, *11*(4), 373-388. https://doi.org/10.1080/09297040490916965
Zamponi, N., Rychlicki, F., Corpaci, L., Cesaroni, E., & Trignani, R. (2008). Vagus nerve stimulation (VNS) is effective in treating catastrophic 1 epilepsy in very young children. *Neurosurg Rev*, *31*(3), 291-297. https://doi.org/10.1007/s10143-008-0134-8
Zandbergen, E. G., Hijdra, A., de Haan, R. J., van Dijk, J. G., Ongerboer de Visser, B. W., Spaans, F., Tavy, D. L., & Koelman, J. H. (2006). Interobserver variation in the interpretation of SSEPs in anoxic-ischaemic coma. *Clin Neurophysiol*, *117*(7), 1529-1535. https://doi.org/10.1016/j.clinph.2006.03.018
Zattara-Hartmann, M. C., & Jammes, Y. (1996). Acute hypoxemia depresses the cardiorespiratory response during phase I constant load exercise and unloaded cycling.

*Arch Physiol Biochem*, *104*(2), 212-219. https://doi.org/10.1076/apab.104.2.212.12885
Zemore, Z., Sharma, A., Carter, K., & Baghdassarian, A. (2018). Delayed Presentation of Tetralogy of Fallot with Isolated Cyanosis. *Case Rep Pediatr*, *2018*, 7412869. https://doi.org/10.1155/2018/7412869
Zeymer, U., Dechend, R., Riemer, T., Deeg, E., Senges, J., Pittrow, D., & Schmieder, R. (2016). Two-Year Outcomes of Patients Treated With Aliskiren Under Clinical Practice Conditions: Non-Interventional Prospective Study. *J Clin Hypertens (Greenwich)*, *18*(7), 647-654. https://doi.org/10.1111/jch.12725
Zhai, X., Jiao, R., Ni, A., & Wang, X. (2023). Case report: Anxiety and depression as initial symptoms in a patient with acute hypoxia and patent foramen ovale. *Front Psychiatry*, *14*, 1229995. https://doi.org/10.3389/fpsyt.2023.1229995
Zhang, B., Tanaka, J., Yang, L., Yang, L., Sakanaka, M., Hata, R., Maeda, N., & Mitsuda, N. (2004). Protective effect of vitamin E against focal brain ischemia and neuronal death through induction of target genes of hypoxia-inducible factor-1. *Neuroscience*, *126*(2), 433-440. https://doi.org/10.1016/j.neuroscience.2004.03.057
Zhang, C., Wei, C. J., Jin, Z., Ma, J., Shen, Y. E., Yu, Q., Fan, Y. B., Xiong, H., & Que, C. L. (2023). Characteristics and feasibility of ambulatory respiratory assessment of paediatric neuromuscular disease: an observational retrospective study. *Int J Neurosci*, *133*(9), 1045-1054. https://doi.org/10.1080/00207454.2022.2042691
Zhang, J. X., Chen, X. Q., Du, J. Z., Chen, Q. M., & Zhu, C. Y. (2005). Neonatal exposure to intermittent hypoxia enhances mice performance in water maze and 8-arm radial maze tasks. *J Neurobiol*, *65*(1), 72-84. https://doi.org/10.1002/neu.20174
Zhang, J. X., Lu, X. J., Wang, X. C., Li, W., & Du, J. Z. (2006). Intermittent hypoxia impairs performance of adult mice in the two-way shuttle box but not in the Morris water maze. *J Neurosci Res*, *84*(1), 228-235. https://doi.org/10.1002/jnr.20860
Zhang, W., Wang, L., Zhu, N., Wu, W., & Liu, H. (2024). A prospective, randomized, single-blinded study comparing the efficacy and safety of dexmedetomidine and propofol for sedation during endoscopic retrograde cholangiopancreatography. *BMC Anesthesiol*, *24*(1), 191. https://doi.org/10.1186/s12871-024-02572-z
Zhang, W., Yin, H., Xu, Y., Fang, Z., Wang, W., Zhang, C., Shi, H., & Wang, X. (2022). The effect of varying inhaled oxygen concentrations of high-flow nasal cannula oxygen therapy during gastroscopy with propofol sedation in elderly patients: a randomized controlled study. *BMC Anesthesiol*, *22*(1), 335. https://doi.org/10.1186/s12871-022-01879-z
Zhao, P., Peng, L., Li, L., Xu, X., & Zuo, Z. (2007). Isoflurane preconditioning improves long-term neurologic outcome after hypoxic-ischemic brain injury in neonatal rats. *Anesthesiology*, *107*(6), 963-970. https://doi.org/10.1097/01.anes.0000291447.21046.4d
Zhao, Y., Packer, C. S., & Rhoades, R. A. (1993). Pulmonary vein contracts in response to hypoxia. *Am J Physiol*, *265*(1 Pt 1), L87-92. https://doi.org/10.1152/ajplung.1993.265.1.L87
Zhou, L. J., Fang, X. Z., Gao, J., Zhangm, Y., & Tao, L. J. (2016). Safety and Efficacy of Dexmedetomidine as a Sedative Agent for Performing Awake Intubation: A Meta-analysis. *Am J Ther*, *23*(6), e1788-e1800. https://doi.org/10.1097/mjt.0000000000000319
Zhou, R., Fu, L., Liu, S., Gao, S., Zhao, Z., Jiang, W., Liu, L., Ren, W., Xiang, D., You, X., Tang, C., Zhou, Y., Song, Y., Xie, J., Xie, L., Yu, R., Zhang, X., Zhou, D., Han, J., . . . Xiong, L. (2024). Influences of Propofol, Ciprofol and Remimazolam on Dreaming During Anesthesia for Gastrointestinal Endoscopy: A Randomized Double-Blind Parallel-Design Trial. *Drug Des Devel Ther*, *18*, 1907-1915. https://doi.org/10.2147/dddt.S455915
Zhou, Z., Wu, L., Zhong, Y., Fang, X., Liu, Y., Chen, H., & Zhang, W. (2017). Gelsemium elegans Poisoning: A Case with 8 Months of Follow-up and Review of the Literature.

*Front Neurol*, *8*, 204. https://doi.org/10.3389/fneur.2017.00204
Zhu, A., Wu, Z., Zhong, X., Ni, J., Li, Y., Meng, J., Du, C., Zhao, X., Nakanishi, H., & Wu, S. (2018). Brazilian Green Propolis Prevents Cognitive Decline into Mild Cognitive Impairment in Elderly People Living at High Altitude. *J Alzheimers Dis*, *63*(2), 551-560. https://doi.org/10.3233/jad-170630
Zinman, R., Corey, M., Coates, A. L., Canny, G. J., Connolly, J., Levison, H., & Beaudry, P. H. (1989). Nocturnal home oxygen in the treatment of hypoxemic cystic fibrosis patients. *J Pediatr*, *114*(3), 368-377. https://doi.org/10.1016/s0022-3476(89)80553-0
Zonnenberg, I. A., Dijk, J. V., Dungen, F., Vermeulen, R. J., & Weissenbruch, M. M. V. (2019). The prognostic value of NIRS in preterm infants with (suspected) late-onset sepsis in relation to long term outcome: A pilot study. *PLoS One*, *14*(7), e0220044. https://doi.org/10.1371/journal.pone.0220044

**Seed PMID References**


Asmaro, D., Mayall, J., & Ferguson, S. (2013). Cognition at altitude: impairment in executive and memory processes under hypoxic conditions. *Aviat Space Environ Med*, *84*(11), 1159-1165. https://doi.org/10.3357/asem.3661.2013

Biswal, S., Sharma, D., Kumar, K., Nag, T. C., Barhwal, K., Hota, S. K., & Kumar, B. (2016). Global hypoxia induced impairment in learning and spatial memory is associated with precocious hippocampal aging. *Neurobiology of Learning and Memory*, *133*, 157-170. https://doi.org/https://doi.org/10.1016/j.nlm.2016.05.011

Bondi, D., Verratti, V., Nori, R., Piccardi, L., Prete, G., Pietrangelo, T., & Tommasi, L. (2021). Spatial Abilities at High Altitude: Exploring the Role of Cultural Strategies and Hypoxia. *High Alt Med Biol*, *22*(2), 157-165. https://doi.org/10.1089/ham.2020.0115

Bonnon, M., Noel-Jorand, M. C., & Therme, P. (2000). Effects of different stay durations on attentional performance during two mountain expeditions. *Aviat Space Environ Med*, *71*(7), 678-684. https://www.ncbi.nlm.nih.gov/pubmed/10902930

Guo, F., Wang, C., Tao, G., Ma, H., Zhang, J., & Wang, Y. (2024). A longitudinal study on the impact of high-altitude hypoxia on perceptual processes. *Psychophysiology*, *61*(6), e14548. https://doi.org/10.1111/psyp.14548

Hencz, A., Magony, A., Thomas, C., Kovacs, K., Szilagyi, G., Pal, J., & Sik, A. (2023). Mild hypoxia-induced structural and functional changes of the hippocampal network. *Front Cell Neurosci*, *17*, 1277375. https://doi.org/10.3389/fncel.2023.1277375

Hutcheon, E. A., Vakorin, V. A., Nunes, A. S., Ribary, U., Ferguson, S., Claydon, V. E., & Doesburg, S. M. (2023). Comparing neuronal oscillations during visual spatial attention orienting between normobaric and hypobaric hypoxia. *Scientific Reports*, *13*(1), 18021. https://doi.org/10.1038/s41598-023-45308-8

Issa, A. N., Herman, N. M., Wentz, R. J., Taylor, B. J., Summerfield, D. C., & Johnson, B. D. (2016). Association of Cognitive Performance with Time at Altitude, Sleep Quality, and Acute Mountain Sickness Symptoms. *Wilderness Environ Med*, *27*(3), 371-378. https://doi.org/10.1016/j.wem.2016.04.008

Kramer, A. F., Coyne, J. T., & Strayer, D. L. (1993). Cognitive function at high altitude. *Hum Factors*, *35*(2), 329-344. https://doi.org/10.1177/001872089303500208

Saputra, N. M., Widyahening, I. S., Mulijadi, H., Mustopo, W. I., Werdhani, R. A., Ibrahim, N., & Wibawanti, R. (2022). Effect of altitude zone exposure on visuospatial function in military aircrew member. *Pan Afr Med J*, *41*, 235. https://doi.org/10.11604/pamj.2022.41.235.22274

Shanjun, Z., Shenwei, X., Bin, X., Huaijun, T., Simin, Z., & Peng, L. (2020). Individual chronic mountain sickness symptom is an early warning sign of cognitive impairment. *Physiol Behav*, *214*, 112748. https://doi.org/10.1016/j.physbeh.2019.112748

Turner, C. E., Barker-Collo, S. L., Connell, C. J., & Gant, N. (2015). Acute hypoxic gas breathing severely impairs cognition and task learning in humans. *Physiol Behav*, *142*, 104-110. https://doi.org/10.1016/j.physbeh.2015.02.006

Yan, X., Zhang, J., Gong, Q., & Weng, X. (2011). Prolonged high-altitude residence impacts verbal working memory: an fMRI study. *Experimental Brain Research*, *208*(3), 437-445. https://doi.org/10.1007/s00221-010-2494-x
