## Supplementary Material 3 for "Neurocognitive Vulnerability to Hypoxia Across Cognitive Domains: A Systematic Review and Meta-Analysis": Supplementary Material 3.html

Neurocognitive Vulnerability During Hypoxia: Differential Effects Across Cognitive Domains


### Neurocognitive Vulnerability During Hypoxia: Differential Effects Across Cognitive Domains

###### Daniel J McKeown

#### 2026-09-08

This report presents the results of a series of meta-analytic models
examining the effects of hypoxia on cognitive performance. For each
cognitive domain, two models are presented, as well as a funnel plot and
a model accounting for publication bias:

- **Main effect model**: Estimates the overall effect of
  hypoxia on the cognitive outcome across studies, without accounting for
  additional predictors.
- **Moderator model**: Includes study-level moderators
  (e.g., severity , duration, and type of exposure, cognitive domain/task
  measure, and participant age) to examine whether these factors explain
  variability in effect sizes across studies.
- **Funnel plot**: Funnel plots are also included to
  visualise the distribution of effect sizes included in each model.
- **Corrected model**: To control for potentional
  publication bias, the main effect model was corrected using the trimfill
  function of the metafor package.

All models were fit using the `metafor` package in R, with
standardized mean differences (SMDH) as the effect size metric and
random effects specified at the study level.

---

#Categorise variables where needed

Effect Sizes and Studies by CHC Broad Ability

| chc\_code | k | studies |
| --- | --- | --- |
| gf | 30 | 14 |
| glr | 11 | 6 |
| gps | 11 | 7 |
| gs | 27 | 16 |
| gsm | 44 | 22 |
| gt | 21 | 9 |
| gv | 8 | 4 |

#Overall cognitive ability#

Random-Effects Model: Overall Cognitive Ability

|  | Estimate | SE | zval | pval | CI\_Lower | CI\_Upper |
| --- | --- | --- | --- | --- | --- | --- |
| Overall Effect | -0.418 | 0.055 | -7.527 | 0 | -0.526 | -0.309 |

Heterogeneity Statistics

| Effect\_Sizes | Tau2 | Tau | I2 | H2 | Q | df | p\_value |
| --- | --- | --- | --- | --- | --- | --- | --- |
| 152 | 0.382 | 0.618 | 87.122 | 7.765 | 886.527 | 151 | 0 |

Pooled Effect and 95 Percent Prediction Interval

| Pooled\_Effect | CI\_Lower | CI\_Upper | PI\_Lower | PI\_Upper |
| --- | --- | --- | --- | --- |
| -0.418 | -0.526 | -0.309 | -1.634 | 0.799 |

Multilevel Moderator Model

| Term | Estimate | SE | zval | pval | CI\_Lower | CI\_Upper |
| --- | --- | --- | --- | --- | --- | --- |
| Intercept | -0.693 | 0.186 | -3.714 | 0.000 | -1.058 | -0.327 |
| Hypoxia Severity | 8.322 | 2.540 | 3.277 | 0.001 | 3.344 | 13.300 |
| Log Duration | -0.011 | 0.054 | -0.211 | 0.833 | -0.118 | 0.095 |
| Participant Age | -0.016 | 0.015 | -1.119 | 0.263 | -0.045 | 0.012 |
| Cognitive Ability: fluid reasoning | 0.326 | 0.091 | 3.590 | 0.000 | 0.148 | 0.504 |
| Cognitive Ability: long-term storage and retrieval | 0.849 | 0.165 | 5.137 | 0.000 | 0.525 | 1.172 |
| Cognitive Ability: processing speed | 0.235 | 0.097 | 2.426 | 0.015 | 0.045 | 0.424 |
| Cognitive Ability: psychomotor speed | -0.035 | 0.134 | -0.264 | 0.791 | -0.297 | 0.226 |
| Cognitive Ability: short-term memory | 0.216 | 0.077 | 2.817 | 0.005 | 0.066 | 0.367 |
| Cognitive Ability: visual processing | 0.491 | 0.124 | 3.954 | 0.000 | 0.247 | 0.734 |
| Hypoxia Type: intermittent hypoxia | 0.997 | 0.477 | 2.088 | 0.037 | 0.061 | 1.932 |
| Hypoxia Type: normobaric hypoxia | 0.133 | 0.241 | 0.553 | 0.580 | -0.339 | 0.605 |

Omnibus Test of Moderators

| QM | df | p\_value |
| --- | --- | --- |
| 65.49 | 11 | 0 |

Residual Heterogeneity in the Moderator Model

| QE | df | p\_value |
| --- | --- | --- |
| 680.204 | 129 | 0 |

Trim-and-Fill Corrected Model

|  | Estimate | SE | zval | pval | CI\_Lower | CI\_Upper |
| --- | --- | --- | --- | --- | --- | --- |
| Trim-and-Fill Adjusted Effect | -0.418 | 0.055 | -7.527 | 0 | -0.526 | -0.309 |

Trim-and-Fill Imputation Results

| Original\_Effect\_Sizes | Imputed\_Effect\_Sizes | Total\_Effect\_Sizes |
| --- | --- | --- |
| 152 | 0 | 152 |

#FIGURE 1#

```
## png 
##   2
```

```
## png 
##   2
```

#FIGURE 2#

### Fluid reasoning ability (Gf)

Main Effect Model: Fluid Reasoning

|  | Estimate | SE | zval | pval | CI\_Lower | CI\_Upper |
| --- | --- | --- | --- | --- | --- | --- |
| Overall Effect | -0.359 | 0.1 | -3.584 | 0 | -0.555 | -0.163 |

Heterogeneity and Model Fit Statistics: Fluid Reasoning

| Effect\_Sizes | Studies | QE | QE\_df | QE\_p | I2 | H2 | Tau2 | Prediction\_Lower | Prediction\_Upper | AIC | BIC |
| --- | --- | --- | --- | --- | --- | --- | --- | --- | --- | --- | --- |
| 30 | 14 | 141.669 | 29 | 0 | 83.089 | 5.913 | 0.234 | -1.328 | 0.61 | 53.267 | 56.002 |

Exploratory Univariable Moderator Models: Fluid Reasoning

|  | Moderator\_Model | Term | Estimate | SE | zval | pval | CI\_Lower | CI\_Upper |
| --- | --- | --- | --- | --- | --- | --- | --- | --- |
| intrcpt…1 | Severity | Intercept | -0.358 | 0.108 | -3.320 | 0.001 | -0.569 | -0.147 |
| Severity\_c | Severity | Severity | 10.482 | 4.433 | 2.364 | 0.018 | 1.793 | 19.171 |
| intrcpt…3 | Duration | Intercept | -0.438 | 0.114 | -3.844 | 0.000 | -0.661 | -0.215 |
| log\_duration\_c | Duration | Log Duration | -0.030 | 0.042 | -0.722 | 0.471 | -0.113 | 0.052 |
| intrcpt…5 | Age | Intercept | -0.378 | 0.127 | -2.982 | 0.003 | -0.627 | -0.130 |
| Age\_c | Age | Age | 0.005 | 0.018 | 0.299 | 0.765 | -0.030 | 0.041 |
| intrcpt…7 | Hypoxia\_Type | Intercept | -0.372 | 0.152 | -2.455 | 0.014 | -0.669 | -0.075 |
| T.2normobaric hypoxia | Hypoxia\_Type | Normobaric vs Hypobaric | -0.001 | 0.169 | -0.009 | 0.993 | -0.334 | 0.331 |

Model Fit Statistics for Exploratory Univariable Moderator Models

| Moderator\_Model | Effect\_Sizes | Studies | QM | QM\_df | QM\_p | QE | QE\_df | QE\_p | Tau2 | AIC | BIC |
| --- | --- | --- | --- | --- | --- | --- | --- | --- | --- | --- | --- |
| Severity | 30 | 14 | 5.591 | 1 | 0.018 | 102.824 | 28 | 0 | 0.127 | 51.884 | 55.881 |
| Duration | 29 | 13 | 0.521 | 1 | 0.471 | 122.232 | 27 | 0 | 0.131 | 53.042 | 56.929 |
| Age | 30 | 14 | 0.089 | 1 | 0.765 | 141.609 | 28 | 0 | 0.185 | 56.327 | 60.324 |
| Hypoxia Type | 30 | 14 | 0.000 | 1 | 0.993 | 136.647 | 28 | 0 | 0.176 | 57.005 | 61.002 |

Data Contributing to Exploratory Moderator Models

| Model | Effect\_Sizes | Studies |
| --- | --- | --- |
| Severity | 30 | 14 |
| Duration | 29 | 13 |
| Age | 30 | 14 |
| Hypoxia Type | 30 | 14 |

Trim-and-Fill Corrected Model: Fluid Reasoning

|  | Estimate | SE | zval | pval | CI\_Lower | CI\_Upper |
| --- | --- | --- | --- | --- | --- | --- |
| Trim-and-Fill Effect | -0.27 | 0.108 | -2.492 | 0.013 | -0.482 | -0.058 |

Trim-and-Fill Imputation Results: Fluid Reasoning

| Original\_Effect\_Sizes | Imputed\_Effect\_Sizes | Total\_Effect\_Sizes |
| --- | --- | --- |
| 30 | 3 | 33 |

### Long-term storage and retrieval ability (Glr)

Main Effect Model: Long-Term Storage and Retrieval

|  | Estimate | SE | zval | pval | CI\_Lower | CI\_Upper |
| --- | --- | --- | --- | --- | --- | --- |
| Overall Effect | -0.601 | 0.221 | -2.717 | 0.007 | -1.034 | -0.167 |

Heterogeneity and Model Fit Statistics: Long-Term Storage and Retrieval

| Effect\_Sizes | Studies | QE | QE\_df | QE\_p | I2 | H2 | Tau2 | Prediction\_Lower | Prediction\_Upper | AIC | BIC |
| --- | --- | --- | --- | --- | --- | --- | --- | --- | --- | --- | --- |
| 11 | 6 | 50.525 | 10 | 0 | 83.351 | 6.006 | 0.405 | -1.921 | 0.72 | 26.73 | 27.336 |

Exploratory Univariable Moderator Models: Long-Term Storage and
Retrieval

|  | Moderator\_Model | Term | Estimate | SE | zval | pval | CI\_Lower | CI\_Upper |
| --- | --- | --- | --- | --- | --- | --- | --- | --- |
| intrcpt…1 | Severity | Intercept | -0.775 | 0.322 | -2.411 | 0.016 | -1.406 | -0.145 |
| Severity\_c | Severity | Severity | 9.735 | 10.344 | 0.941 | 0.347 | -10.538 | 30.009 |
| intrcpt…3 | Duration | Intercept | -0.701 | 0.385 | -1.821 | 0.069 | -1.455 | 0.053 |
| log\_duration\_c | Duration | Log Duration | 0.169 | 0.156 | 1.083 | 0.279 | -0.137 | 0.476 |
| intrcpt…5 | Age | Intercept | -0.586 | 0.388 | -1.510 | 0.131 | -1.347 | 0.175 |
| Age\_c | Age | Age | -0.028 | 0.031 | -0.914 | 0.361 | -0.089 | 0.032 |

Model Fit Statistics for Exploratory Univariable Moderator Models:
Long-Term Storage and Retrieval

| Moderator\_Model | Effect\_Sizes | Studies | QM | QM\_df | QM\_p | QE | QE\_df | QE\_p | Tau2 | AIC | BIC |
| --- | --- | --- | --- | --- | --- | --- | --- | --- | --- | --- | --- |
| Severity | 11 | 6 | 0.886 | 1 | 0.347 | 42.176 | 9 | 0.000 | 0.542 | 23.347 | 23.939 |
| Duration | 9 | 4 | 1.173 | 1 | 0.279 | 22.658 | 7 | 0.002 | 0.186 | 14.244 | 14.082 |
| Age | 11 | 6 | 0.835 | 1 | 0.361 | 47.083 | 9 | 0.000 | 0.546 | 21.662 | 22.253 |

Trim-and-Fill Corrected Model: Long-Term Storage and Retrieval

|  | Estimate | SE | zval | pval | CI\_Lower | CI\_Upper |
| --- | --- | --- | --- | --- | --- | --- |
| Trim-and-Fill Effect | -0.601 | 0.221 | -2.717 | 0.007 | -1.034 | -0.167 |

Trim-and-Fill Imputation Results: Long-Term Storage and Retrieval

| Original\_Effect\_Sizes | Imputed\_Effect\_Sizes | Total\_Effect\_Sizes |
| --- | --- | --- |
| 11 | 0 | 11 |

#Psychomotor speed#

Main Effect Model: Psychomotor Speed

|  | Term | Estimate | SE | zval | pval | CI\_Lower | CI\_Upper |
| --- | --- | --- | --- | --- | --- | --- | --- |
| intrcpt | Overall Effect | -0.503 | 0.231 | -2.184 | 0.029 | -0.955 | -0.052 |

Heterogeneity and Prediction Interval Statistics

| Effect\_Sizes | Studies | QE | QE\_df | QE\_p | I2 | Tau2 | Prediction\_Lower | Prediction\_Upper |
| --- | --- | --- | --- | --- | --- | --- | --- | --- |
| 11 | 7 | 65.331 | 10 | 0 | 86.88 | 0.487 | -1.944 | 0.937 |

Exploratory Univariable Moderator Models: Psychomotor Speed

|  | Moderator\_Model | Term | Estimate | SE | zval | pval | CI\_Lower | CI\_Upper |
| --- | --- | --- | --- | --- | --- | --- | --- | --- |
| intrcpt…1 | Severity | Intercept | -0.515 | 0.279 | -1.847 | 0.065 | -1.062 | 0.031 |
| Severity\_c | Severity | Severity | -0.909 | 12.786 | -0.071 | 0.943 | -25.969 | 24.151 |
| intrcpt…3 | Age | Intercept | -0.381 | 0.321 | -1.186 | 0.236 | -1.010 | 0.248 |
| Age\_c | Age | Age | -0.024 | 0.029 | -0.810 | 0.418 | -0.081 | 0.034 |

Model Fit Statistics for Exploratory Moderator Models

| Model | Effect\_Sizes | Studies | QM | QM\_df | QM\_p | QE | QE\_df | QE\_p | Tau2 |
| --- | --- | --- | --- | --- | --- | --- | --- | --- | --- |
| Severity | 11 | 7 | 0.005 | 1 | 0.943 | 57.474 | 9 | 0 | 0.478 |
| Age | 11 | 7 | 0.655 | 1 | 0.418 | 56.779 | 9 | 0 | 0.473 |

Data Contributing to Exploratory Moderator Models

| Model | Effect\_Sizes | Studies |
| --- | --- | --- |
| Severity | 11 | 7 |
| Age | 11 | 7 |

Trim-and-Fill Corrected Model: Psychomotor Speed

|  | Term | Estimate | SE | zval | pval | CI\_Lower | CI\_Upper |
| --- | --- | --- | --- | --- | --- | --- | --- |
| intrcpt | Trim-and-Fill Effect | -0.598 | 0.233 | -2.57 | 0.01 | -1.053 | -0.142 |

Trim-and-Fill Imputation Results

| Original\_Effect\_Sizes | Imputed\_Effect\_Sizes | Total\_Effect\_Sizes |
| --- | --- | --- |
| 11 | 1 | 12 |

### Processing speed

Main Effect Model: Processing Speed

|  | Term | Estimate | SE | zval | pval | CI\_Lower | CI\_Upper |
| --- | --- | --- | --- | --- | --- | --- | --- |
| intrcpt | Overall Effect | -0.615 | 0.184 | -3.343 | 0.001 | -0.976 | -0.255 |

Heterogeneity and Prediction Interval Statistics

| Effect\_Sizes | Studies | QE | QE\_df | QE\_p | I2 | Tau2 | Prediction\_Lower | Prediction\_Upper |
| --- | --- | --- | --- | --- | --- | --- | --- | --- |
| 27 | 16 | 211.202 | 26 | 0 | 92.07 | 0.803 | -2.408 | 1.178 |

Exploratory Univariable Moderator Models: Processing Speed

|  | Moderator\_Model | Term | Estimate | SE | zval | pval | CI\_Lower | CI\_Upper |
| --- | --- | --- | --- | --- | --- | --- | --- | --- |
| intrcpt…1 | Severity | Intercept | -0.474 | 0.208 | -2.276 | 0.023 | -0.882 | -0.066 |
| Severity\_c | Severity | Severity | 11.328 | 10.542 | 1.075 | 0.283 | -9.333 | 31.989 |
| intrcpt…3 | Duration | Intercept | -0.535 | 0.238 | -2.251 | 0.024 | -1.000 | -0.069 |
| log\_duration\_c | Duration | Log Duration | -0.045 | 0.063 | -0.709 | 0.478 | -0.169 | 0.079 |
| intrcpt…5 | Age | Intercept | -0.476 | 0.219 | -2.171 | 0.030 | -0.905 | -0.046 |
| Age\_c | Age | Age | 0.000 | 0.017 | 0.020 | 0.984 | -0.034 | 0.034 |

Model Fit Statistics for Exploratory Moderator Models

| Model | Effect\_Sizes | Studies | QM | QM\_df | QM\_p | QE | QE\_df | QE\_p | Tau2 |
| --- | --- | --- | --- | --- | --- | --- | --- | --- | --- |
| Severity | 27 | 16 | 1.155 | 1 | 0.283 | 205.946 | 25 | 0 | 0.624 |
| Duration | 25 | 14 | 0.503 | 1 | 0.478 | 198.278 | 23 | 0 | 0.707 |
| Age | 27 | 16 | 0.000 | 1 | 0.984 | 210.551 | 25 | 0 | 0.645 |

Data Contributing to Exploratory Moderator Models

| Model | Effect\_Sizes | Studies |
| --- | --- | --- |
| Severity | 27 | 16 |
| Duration | 25 | 14 |
| Age | 27 | 16 |

Trim-and-Fill Corrected Model: Processing Speed

|  | Term | Estimate | SE | zval | pval | CI\_Lower | CI\_Upper |
| --- | --- | --- | --- | --- | --- | --- | --- |
| intrcpt | Trim-and-Fill Effect | -0.615 | 0.184 | -3.343 | 0.001 | -0.976 | -0.255 |

Trim-and-Fill Imputation Results

| Original\_Effect\_Sizes | Imputed\_Effect\_Sizes | Total\_Effect\_Sizes |
| --- | --- | --- |
| 27 | 0 | 27 |

### Short-term memory (Gsm)

Main Effect Model: Short-Term Memory (Gsm)

|  | Term | Estimate | SE | zval | pval | CI\_Lower | CI\_Upper |
| --- | --- | --- | --- | --- | --- | --- | --- |
| intrcpt | Overall Effect | -0.355 | 0.104 | -3.419 | 0.001 | -0.559 | -0.152 |

Model Fit and Heterogeneity Statistics: Short-Term Memory (Gsm)

| Statistic | Value |
| --- | --- |
| Effect Sizes (k) | 44.000 |
| Independent Studies | 22.000 |
| Q | 247.944 |
| Q df | 43.000 |
| Q p-value | 0.000 |
| I² (%) | 87.227 |
| Tau² | 0.383 |
| Tau | 0.618 |
| 95% Prediction Interval Lower | -1.585 |
| 95% Prediction Interval Upper | 0.874 |

Exploratory Univariable Moderator Models: Short-Term Memory (Gsm)

|  | Moderator\_Model | Term | Estimate | SE | zval | pval | CI\_Lower | CI\_Upper |
| --- | --- | --- | --- | --- | --- | --- | --- | --- |
| intrcpt…1 | Severity | Intercept | -0.401 | 0.155 | -2.592 | 0.010 | -0.704 | -0.098 |
| Severity\_c | Severity | Severity | 4.665 | 4.323 | 1.079 | 0.281 | -3.808 | 13.138 |
| intrcpt…3 | Duration | Intercept | -0.319 | 0.193 | -1.657 | 0.098 | -0.697 | 0.059 |
| log\_duration\_c | Duration | Log Duration | -0.020 | 0.034 | -0.596 | 0.551 | -0.086 | 0.046 |
| intrcpt…5 | Age | Intercept | -0.411 | 0.160 | -2.574 | 0.010 | -0.725 | -0.098 |
| Age\_c | Age | Age | -0.022 | 0.020 | -1.120 | 0.263 | -0.060 | 0.016 |
| intrcpt…7 | Hypoxia Type | Intercept | -0.494 | 0.167 | -2.952 | 0.003 | -0.822 | -0.166 |
| T.2normobaric hypoxia | Hypoxia Type | Hypoxia Type: normobaric hypoxia | 0.151 | 0.144 | 1.053 | 0.292 | -0.130 | 0.433 |

Model Fit Statistics for Exploratory Moderator Models: Short-Term Memory
(Gsm)

| Model | Effect\_Sizes | Studies | QE | QE\_df | QE\_p | QM | QM\_df | QM\_p | AIC | BIC | LogLik | Tau2 |
| --- | --- | --- | --- | --- | --- | --- | --- | --- | --- | --- | --- | --- |
| Severity | 44 | 22 | 202.848 | 42 | 0 | 1.164 | 1 | 0.281 | 57.792 | 63.005 | -25.896 | 0.442 |
| Duration | 40 | 18 | 226.326 | 38 | 0 | 0.355 | 1 | 0.551 | 52.502 | 57.415 | -23.251 | 0.575 |
| Age | 44 | 22 | 243.416 | 42 | 0 | 1.254 | 1 | 0.263 | 56.895 | 62.108 | -25.448 | 0.471 |
| Hypoxia Type | 42 | 21 | 227.746 | 40 | 0 | 1.110 | 1 | 0.292 | 55.479 | 60.546 | -24.740 | 0.442 |

Data Contributing to Exploratory Moderator Models: Short-Term Memory
(Gsm)

| Model | Effect\_Sizes | Studies |
| --- | --- | --- |
| Severity | 44 | 22 |
| Duration | 40 | 18 |
| Age | 44 | 22 |
| Hypoxia Type | 42 | 21 |

Trim-and-Fill Corrected Model: Short-Term Memory (Gsm)

|  | Term | Estimate | SE | zval | pval | CI\_Lower | CI\_Upper |
| --- | --- | --- | --- | --- | --- | --- | --- |
| intrcpt | Trim-and-Fill Effect | -0.355 | 0.104 | -3.419 | 0.001 | -0.559 | -0.152 |

Trim-and-Fill Imputation Summary: Short-Term Memory (Gsm)

| Statistic | Value |
| --- | --- |
| Observed Effect Sizes | 44 |
| Imputed Effect Sizes | 0 |
| Total Effect Sizes After Trim-and-Fill | 44 |

### Decision speed / reaction time (Gt)

Main Effect Model: Decision Speed / Reaction Time (Gt)

|  | Term | Estimate | SE | zval | pval | CI\_Lower | CI\_Upper |
| --- | --- | --- | --- | --- | --- | --- | --- |
| intrcpt | Overall Effect | -0.333 | 0.108 | -3.071 | 0.002 | -0.545 | -0.12 |

Model Fit and Heterogeneity Statistics: Decision Speed / Reaction Time
(Gt)

| Statistic | Value |
| --- | --- |
| Effect Sizes (k) | 21.000 |
| Independent Studies | 9.000 |
| Q | 95.890 |
| Q df | 20.000 |
| Q p-value | 0.000 |
| I² (%) | 78.771 |
| Tau² | 0.189 |
| Tau | 0.435 |
| 95% Prediction Interval Lower | -1.211 |
| 95% Prediction Interval Upper | 0.546 |

Exploratory Univariable Moderator Models: Decision Speed / Reaction Time
(Gt)

|  | Moderator\_Model | Term | Estimate | SE | zval | pval | CI\_Lower | CI\_Upper |
| --- | --- | --- | --- | --- | --- | --- | --- | --- |
| intrcpt…1 | Severity | Intercept | -0.268 | 0.086 | -3.105 | 0.002 | -0.437 | -0.099 |
| Severity\_c | Severity | Severity | 7.760 | 3.972 | 1.954 | 0.051 | -0.026 | 15.545 |
| intrcpt…3 | Duration | Intercept | -0.258 | 0.106 | -2.433 | 0.015 | -0.467 | -0.050 |
| log\_duration\_c | Duration | Log Duration | -0.005 | 0.043 | -0.111 | 0.911 | -0.089 | 0.080 |
| intrcpt…5 | Age | Intercept | -0.259 | 0.082 | -3.174 | 0.002 | -0.419 | -0.099 |
| Age\_c | Age | Age | 0.013 | 0.007 | 2.016 | 0.044 | 0.000 | 0.026 |
| intrcpt…7 | Hypoxia Type | Intercept | -0.209 | 0.122 | -1.711 | 0.087 | -0.448 | 0.030 |
| T.2normobaric hypoxia | Hypoxia Type | Hypoxia Type: normobaric hypoxia | -0.184 | 0.148 | -1.239 | 0.215 | -0.475 | 0.107 |

Model Fit Statistics for Exploratory Moderator Models: Decision Speed /
Reaction Time (Gt)

| Model | Effect\_Sizes | Studies | QE | QE\_df | QE\_p | QM | QM\_df | QM\_p | AIC | BIC | LogLik | Variance\_Component |
| --- | --- | --- | --- | --- | --- | --- | --- | --- | --- | --- | --- | --- |
| Severity | 21 | 9 | 87.949 | 19 | 0 | 3.816 | 1 | 0.051 | 66.789 | 69.622 | -30.395 | 0.033 |
| Duration | 21 | 9 | 90.906 | 19 | 0 | 0.012 | 1 | 0.911 | 69.885 | 72.718 | -31.943 | 0.055 |
| Age | 21 | 9 | 86.614 | 19 | 0 | 4.064 | 1 | 0.044 | 66.421 | 69.255 | -30.211 | 0.027 |
| Hypoxia Type | 19 | 8 | 77.856 | 17 | 0 | 1.535 | 1 | 0.215 | 65.097 | 67.596 | -29.548 | 0.014 |

Data Contributing to Exploratory Moderator Models: Decision Speed /
Reaction Time (Gt)

| Model | Effect\_Sizes | Studies |
| --- | --- | --- |
| Severity | 21 | 9 |
| Duration | 21 | 9 |
| Age | 21 | 9 |
| Hypoxia Type | 19 | 8 |

Trim-and-Fill Corrected Model: Decision Speed / Reaction Time (Gt)

|  | Term | Estimate | SE | zval | pval | CI\_Lower | CI\_Upper |
| --- | --- | --- | --- | --- | --- | --- | --- |
| intrcpt | Trim-and-Fill Effect | -0.141 | 0.113 | -1.252 | 0.211 | -0.361 | 0.08 |

Trim-and-Fill Imputation Summary: Decision Speed / Reaction Time (Gt)

| Statistic | Value |
| --- | --- |
| Observed Effect Sizes | 21 |
| Imputed Effect Sizes | 6 |
| Total Effect Sizes After Trim-and-Fill | 27 |

### Visual processing (Gv)

Main Effect Model: Visual Processing

| Term | Estimate | SE | zval | pval | CI\_Lower | CI\_Upper |
| --- | --- | --- | --- | --- | --- | --- |
| Overall effect | -0.289 | 0.252 | -1.147 | 0.251 | -0.782 | 0.205 |

Heterogeneity Statistics: Visual Processing

| Statistic | Value |
| --- | --- |
| Effect sizes | 8 |
| Independent studies | 4 |
| Tau squared | 0.448 |
| Tau | 0.669 |
| I squared (%) | 89.237 |
| Q | 52.997 |
| Q df | 7 |
| Q p-value | <0.001 |

Prediction Interval: Visual Processing

| Estimate | PI\_Lower | PI\_Upper |
| --- | --- | --- |
| -0.289 | -1.69 | 1.113 |

Trim-and-Fill Corrected Model: Visual Processing

| Term | Estimate | SE | zval | pval | CI\_Lower | CI\_Upper |
| --- | --- | --- | --- | --- | --- | --- |
| Trim-and-fill adjusted effect | -0.289 | 0.252 | -1.147 | 0.251 | -0.782 | 0.205 |

Trim-and-Fill Diagnostics: Visual Processing

| Statistic | Value |
| --- | --- |
| Observed effect sizes | 8 |
| Imputed effect sizes | 0 |
| Total effect sizes | 8 |
